# High-throughput transcriptomic analysis of human primary hepatocyte spheroids exposed to per- and polyfluoroalkyl substances (PFAS) as a platform for relative potency characterization

**DOI:** 10.1101/2020.10.15.341362

**Authors:** A. Rowan-Carroll, A. Reardon, K. Leingartner, R. Gagné, A. Williams, M.J. Meier, B. Kuo, J. Bourdon-Lacombe, I. Moffat, R. Carrier, A. Nong, L. Lorusso, S.S. Ferguson, E. Atlas, C. Yauk

## Abstract

Per- and poly-fluoroalkyl substances (PFAS) are widely found in the environment because of their extensive use and persistence. Although several PFAS are well studied, most lack toxicity data to inform human health hazard and risk assessment. This study focussed on four model PFAS: perfluorooctanoic acid (PFOA; 8 carbon), perfluorobutane sulfonate (PFBS; 4 carbon), perfluorooctane sulfonate (PFOS; 8 carbon), and perfluorodecane sulfonate (PFDS; 10 carbon). Human primary liver cell spheroids (pooled from 10 donors) were exposed to 10 concentrations of each PFAS and analyzed at four time-points. The approach aimed to: (1) identify gene expression changes mediated by the PFAS; (2) identify similarities in biological responses; (3) compare PFAS potency through benchmark concentration analysis; and (4) derive bioactivity exposure ratios (ratio of the concentration at which biological responses occur, relative to daily human exposure). All PFAS induced transcriptional changes in cholesterol biosynthesis and lipid metabolism pathways, and predicted PPARα activation. PFOS exhibited the most transcriptional activity and had a highly similar gene expression profile to PFDS. PFBS induced the least transcriptional changes and the highest benchmark concentration (i.e., was the least potent). The data indicate that these PFAS may have common molecular targets and toxicities, but that PFOS and PFDS are the most similar. The transcriptomic bioactivity exposure ratios derived here for PFOA and PFOS were comparable to those derived using rodent apical endpoints in risk assessments. These data provide a baseline level of toxicity for comparison with other known PFAS using this testing strategy.

## Introduction

Per- and poly-fluoroalkyl substances (PFAS) are a class of chemicals ubiquitously found in the environment. Their unique hydrophobic and lipophobic properties have led to their use in many consumer products and their persistence, and bioaccumulation potential, led to their detection in wildlife and humans worldwide (Giesy and Kanna 2001; Armitage *et al*., 2006; 2009a, Wania, 2007). Although perfluorooctane sulfonate (PFOS) and perfluorooctanoate (PFOA) have been discontinued from North American production, they continue to be found in drinking water and at contaminated sites within Canada (Health Canada., 2018a and b). Moreover, there are thousands of PFAS that remain in production, with little toxicological data available for the vast majority (OECD/UNEP Global PFC Group, 2018).

The adverse health effects of PFOS and PFOA in humans are well documented, including increased cholesterol levels (Erikson *et al*., 2013; Geiger *et al*., 2014), decreased human fertility (Bach *et al*., 2016), and reduced immune response (NTP Monograph, 2016; Kielsen *et al*., 2016; Health Canada, 2018a and b). Recently, the US Environmental Protection Agency developed an action plan to address PFAS in drinking water, general exposure reduction, safety in commerce, and support for communications (EPA, 2020).

The effects of several PFAS have been extensively studied in animal models. For example, wild-type and PPARα-null mice treated with several PFAS exhibit steatosis, α which is characterised by an accumulation of lipids in hepatocytes, leading to liver disease (Das *et al*., 2017). Furthermore, rats dosed with PFOS (up to 1.14 mg/kg/day) and offspring of mice dosed with PFOA (up to 5 mg/kg/day) developed liver cancer (Bjork *et al*., 2011; Filgo *et al*., 2015).

Increased exposure to PFAS has been associated with adverse liver outcomes and the development of steatosis. Using NHANES data, increased PFAS exposure was associated with alterations in select liver biomarkers, suggesting they play a role in the progression of non-alcoholic fatty liver disease (Jain and Ducatman, 2019). Both *in vitro* and *in vivo* studies suggest that the mechanism by which these compounds act involves increased intracellular triglycerides levels and activation of nuclear receptors (Louisse *et al*. 2020, Schlezinger *et al.,* 2020). Given these noted adverse liver effects, the liver was identified as the critical epidemiological endpoint in Health Canada’s existing regulatory guidelines on PFAS (Health Canada, 2018a and b). In humans, median serum levels can reach 80 ng/ml for PFOA and 22 ng/ml for PFOS and these exposure levels are correlated with increased blood lipid levels (Steenland *et al*., 2009). Conversely, a review from Chang et al. (2014) found no apparent causal association with cancer in human occupational exposures to PFOA that are an order of magnitude higher (serum levels ranging from 160 to 2880 ng/ml) than environmental exposures.

Acquiring information on data-poor substances for risk assessment has been a challenge for regulatory agencies worldwide given the cost and time required to conduct traditional toxicological research. To accelerate the pace of chemical risk assessments, global efforts have focused on increasing the use of New Approach Methodologies (NAMs) (Kavlock *et al*., 2018; Krewski *et al*., 2020). *In silico* predictions, high-throughput screening, novel *in vitro* models, *in vitro* to *in vivo* extrapolation (IVIVE), and read-across are some of the NAMs with increasing use in risk assessment for chemicals with limited data. High-throughput transcriptomics is a particularly powerful method for obtaining information about the broad scope of biological perturbations that result from chemical exposures. New technologies now enable gene expression analyses directly from cell lysates, which significantly increases the ease and speed with which transcriptomic data can be produced (Trejo *et al*., 2019).

To address the need for efficient assessment of data-poor PFAS, we sought to explore transcriptional responses across exposure time with a broad concentration range in order to refine a streamlined set of exposure conditions for PFAS evaluations.

We developed an approach to broadly screen for PFAS-induced biological perturbations through global gene expression profiling of a primary human liver cell spheroid model. A main objective was to establish baseline transcriptomic profiles and an analytical pipeline for future evaluations of data-poor PFAS. Spheroids were exposed to nine concentrations of PFAS and sampled at four time points. To establish our approach, we focused on four prototype PFAS: short-chain perfluorobutane sulfonate (PFBS), long-chain perfluorodecane sulfonate (PFDS), and PFOA and PFOS. Microscopy was used to evaluate phenotypic changes in the spheroids, and the lactose dehydrogenase activity assay was used to assess cytotoxicity over time. The TempO-Seq platform from BioSpyder using the NIEHS S1500 gene set was applied for transcriptomic analysis (House *et al*., 2017; Mav *et al*., 2018). PFAS were compared based on the induction of differentially expressed genes, transcriptional patterns measured by hierarchical clustering and principal component analysis, pathway enrichment, upstream regulator enrichment, and transcriptomic benchmark concentrations. Bioactivity-exposure ratios for PFOS and PFOA were calculated by comparing human administered equivalent dose (the daily dose required to achieve the *in vitro* concentration of interest) to human exposure levels. A small bioactivity exposure ratio suggests that biological perturbations occur at concentrations that approach external human exposure levels. This initial research project was used to inform experimental design and analytical approach for analysis of a second larger group of data-poor PFAS (Reardon *et al*., BioRxiv **doi**: https://doi.org/10.1101/2020.10.20.347328).

## Methods

### Chemicals and preparation of working solutions

PFOS (95% purity CAS 1763-23-1), PFOA (95% purity CAS 335-67-1), PFBS (95% purity CAS 375-73-5) and Cyclosporine A (CsA) were purchased from Sigma-Aldrich (Oakville, ON) and PFDS was purchased from Toronto Research Chemicals (95% purity CAS 335-77-3 Toronto, ON). Chemicals were dissolved in dimethylsulfoxide (DMSO, Sigma-Aldrich, Oakville, ON) to prepare working stock solutions ranging in concentration from 0.02 to 30 mM. Each PFAS was tested at 10 different concentrations (0 to 100 µM), in quadruplicate experiments. For microscopic characterization, a 30 mM working stock of CsA in DMSO was also prepared. The final concentration of DMSO in the media was 0.1% for all PFAS at all concentrations up to 50 μM; the DMSO was μ M and 0.3% for 100 μM. The %DMSO varied at the two top μ concentrations because chemical stocks were diluted to 30 mM for all PFAS due to solubility. All experimental concentrations of PFAS were matched to vehicle only (DMSO) time-matched and concentration-matched controls; e.g., PFAS at 100 µM (0.3% DMSO) was matched to 0.3% DMSO controls from the same time point.

### Cell Culture

3D InSight™ Human Liver Microtissues were purchased from InSphero (Brunswick, ME) in a 96 well format, with a single spheroid per well. These spheroids are a co-culture model pooled from 10 different human liver donors and are a metabolically active system comprised of isolated primary hepatocytes and Kupffer cells. Upon arrival, the media were replaced with serum free cell medium (InSphero Human Liver Maintenance Medium–Tox) and spheroids were allowed to acclimate for 24 hours prior to PFAS exposures. The Human Liver Maintenance Medium–Tox contains no additives or serum and is designed for testing of drug induced liver injury (DILI) compounds. The media contains no proteins or albumin; however, the spheroids themselves do secrete albumin into the media.

Liver spheroids were cultured under sterile conditions with incubation at 37 ^°^C, 5% CO_2_ as per the materials and instructions provided by the manufacturer. Liver spheroids were exposed for 1, 4, 10 or 14 days to PFBS, PFOA, PFOS, PFDS, or DMSO control diluted in culture medium. Final exposure concentrations were 0.02, 0.1, 0.2, 1, 2, 10, 20, 50, and 100 μM of each of the four PFAS chemicals. The entire experiment included eight 96-well plates of spheroids. Each plate contained every concentration of each PFAS alongside a matched DMSO control for every sample. On each plate there were also 18 DMSO controls: four 0.3% DMSO (for 100 μM concentrations), four 0.17% DMSO (for 50 μM concentrations), and ten 0.1% DMSO (for all of the remaining concentrations). There were two 96-well plates for every time point. PFAS or DMSO, at the desired concentrations, were replenished each time the media were changed (every 3 days).

The highest concentration tested was 100 µM, as per the US EPA’s ToxCast program. Although the top concentration is high, it is useful to have a top concentration yielding a robust transcriptomic response for benchmark concentration modeling and this provides important context analogous to LD_50_ for guideline toxicology studies. From the top concentration we scaled down in total by 3 logs to the lowest, most environmentally relevant concentration (Health Canada., 2017; 2018a and b). Media were replaced every 3 days with freshly diluted chemical, and spent media were reserved for cytotoxicity analysis. Spheroids designated for microscopy were treated with 0.1% DMSO; 48 hours prior to the day-14 time point half of these spheroids were exposed to 30 µM CsA for immunostaining. Spheroids designated for TempO-Seq library building and sequencing were washed with Dulbecco’s phosphate-buffered saline (DPBS; Thermo Fisher Scientific Waltham, MA) and lysed using 5-7 µl of 2X TempO-Seq lysis buffer (BioSpyder Technologies Inc Carlsbad, CA). Liver spheroids were triturated and allowed to complete lysis for approximately 10 minutes before freezing at -80 ^°^C. Spheroids designated for TempO-Seq library building and sequencing analysis were collected after exposure on day-1, day-4, day-10, and day-14.

### Cytotoxicity Assessment

Cytotoxicity was measured from spent culture media collected every 3 days during media replenishment (i.e., on days 3, 6, 9, 12, and then on day-14 at study termination, versus days 1, 4, 10, and 14 for transcriptomic analyses below) using the lactate dehydrogenase assay (LDH-Glo Cytotoxicity Assay Promega Corp Madison, USA), as per manufacturer’s instructions. Briefly, media were diluted 1:10 in storage buffer (200 mM Tris-HCl, pH 7.3, containing 10% glycerol and 1% bovine serum albumin) and kept frozen at -80 ^°^C until analysis. For the assay, thawed samples were diluted with an equal volume of LDH Detection Reagent and equilibrated for 60 minutes at room temperature before reading the luminescence (GloMax 96 Microplate Luminometer, Promega Corp Madison, WI USA) in relative luminescence units (RLU). Cytotoxicity was determined after performing statistical analysis of the RLU data after blank subtraction in Sigma Plot 13™, using the Kruskal-Wallis one-way analysis of variance on ranks, followed by the Dunn’s multiple comparison versus the control group for each time point, with n = 4 for all time points, with exceptions on Day-14 (n = 2 in some cases as indicated). The sample RLU was converted to a ratio by division with the average RLU of the DMSO controls and the mean and standard deviations of these ratios were calculated for each treatment and time point. The release of LDH was expressed as a fold-change over DMSO controls. Overall cytotoxicity was determined by combining three metrics of cell viability: LDH release greater than 10-fold over the control, low read counts from Next Generation Sequencing (described further below), and visual examination of cell morphology.

### Microscopy

Spheroid whole mounts were prepared on Day-4 and Day-14 according to a protocol provided by InSphero Inc. (Technical Protocol 008 c. 2015), with modifications. Briefly, spheroids were fixed with 4% paraformaldehyde in phosphate buffered saline (PBS) for 1 hour, permeabilized with 0.2% Triton-X100 in PBS for 30 minutes, and blocked with 5% bovine serum albumin in PBS for 1 hour at room temperature. On day-4, the spheroids were incubated with CD-68 antibody (1:100; Santa Cruz CD68 (E-11): sc-17832 conjugated to AlexaFluor 594) overnight at 4 °C, followed by incubation with 1:40 phalloidin (Life Technologies 12379 conjugated to AlexaFluor 488 Carlsbad, CA) and DAPI (2 µg/ml; Sigma D8417 Oakville, ON) in PBS for 45 minutes at room temperature. This was performed to ensure spheroid integrity and cell composition.

On Day-14, spheroids were stained with CYP3A4 primary antibody (1:00; Santa Cruz CYP3A4 (HL3): sc-53850 conjugated to AlexaFluor 647 Dallas, TX) phalloidin and DAPI as described above, or with Nile Red (1 µg/ml; Sigma 19123 Oakville, ON) and DAPI (2 µg/ml; Sigma D8417 Oakville, ON) in PBS for 30 minutes at room temperature. After antibody incubation and three washes with PBS, spheroids were transferred to glass microscope slides and mounted with Prolong Gold Mounting Medium (Invitrogen P36934, Burlington, ON). Confocal laser-scanning immunofluorescence microscopy was performed using a Leica TCS SP8 microscope.

### TempO-Seq Library Building and Next Generation Sequencing

Gene expression was measured using the human TempO-Seq S1500 panel (BioSpyder Technologies Inc. Carlsbad, CA) (Mav *et al*., 2018). This targeted RNA-seq panel is comprised of 3000 genes selected by the NIEHS to represent a biologically diverse set of pathways that are toxicologically responsive (https://federalregister.gov/a/2015-08529). For TempO-Seq analysis, liver spheroids were exposed to nine concentrations of the four PFAS and solvent control (DMSO), over the four time points (days 1, 4, 10, and 14). Spheroids were lysed *in situ* using 2 x TempO-Seq lysis buffer diluted with an equal amount of PBS. Lysates and positive technical controls (Human Universal Reference RNA - uhrRNA Agilent Cat# 740000, Santa Clara, California and Human Brain Total RNA brRNA - ThermoFisher AM7962, Waltham, Massachusetts), as well as no-cell negative controls (1X TempO-Seq lysis buffer alone) were hybridized to the detector oligo mix following the manufacturer’s instructions (TEMPO-SEQ Human Tox +Surrogate with Standard Attenuation Transcriptome Kit (96 Samples) BioSpyder Technologies, Inc. Hybridization was followed by nuclease digestion of excess oligos, detector oligo ligation, and amplification of the product with the tagged primers according to manufacturer instructions. At the time of amplification, each sample was ligated to a sample-specific bar code that allows for combining samples for sequencing purposes. Labelled amplicons were then pooled and purified using NucleoSpin Gel and PCR Clean-up kits, (Takara Bio USA, Inc. Mountain View, CA). Libraries were sequenced in-house using a NextSeq 500 High-Throughput Sequencing System (Illumina San Diego, California) using 50 cycles from a 75-cycle high throughput flow cell. A median read depth of 2 million reads/sample was achieved.

### Data quality overview and removal of transcriptomic outliers

The gene expression data are available in the NCBI’s Gene Expression Omnibus (Bioproject number PRJNA604830). Initial quality assessment was done on water-only controls and reference standards included on every TempO-Seq plate. We first confirmed very little signal (< 0.3%) from samples with no cells/RNA. In addition, we confirmed that commercial human universal reference RNA (two per plate – 16 in total) showed high Spearman correlation coefficients (> 0.94) across all plate comparisons indicating a high degree of technical reproducibility across the plates.

The median read depth for the study was two million reads with a minimum read depth of 400,000 for retained samples. The median range of % mapped reads across experimental space using BioSpyder’s script was: day-1: 71%, day-4: 62%, day-10: 49%, and day-14: 53%. The cytotoxicity information was paired with the number of reads recovered during the TempO-Seq experiments to identify overtly cytotoxic concentrations for removal from the transcriptomic data analysis. The conditions removed from subsequent analyses were: 100 µM PFOA on days 10 and 14, and 50 µM on day-14; and 50 and 100 µM PFOS from all time points.

### Data processing and identification of differentially expressed genes

Reads were extracted from the bcl files and demultiplexed (i.e., assigned to respective sample files) with bcl2fastq v. 2.20.0.42. The fastq files were processed with the “pete. star. script_v3.0” supplied by BioSpyder. Briefly, the script uses star v.2.5 to align the reads and the count function from QuasR to extract the feature counts specified in a gtf file from the aligned reads.

Samples with low read counts (library sizes < 100,000 reads) were removed from the study. As described above, read counts were also used to identify overtly cytotoxic concentrations alongside LDH assay results as samples with low read counts were often associated with high levels of cytotoxicity as measured in the LDH assay. Probes with low counts were flagged as absent genes and removed (probes that did not exceed a median of 5 counts in at least one group). Dendrogram plots (Supplemental Files1-1 through 1-9) were generated on all the remaining genes in R using the hclust () function with complete-linkage for each time point using a distance metric defined as 1- Spearman correlation. Samples that clustered as singletons when cutting the dendrograms at the 0.1 dissimilarity were removed from the study. To account for differences in the percentage of DMSO used, the data were normalized to matched controls and rescaled to the average of the 0.1% DMSO dose group. Boxplots of the log2 CPM (counts per million) were plotted to ensure that the samples within each treatment group had similar medians and interquartile ranges (Supplemental Files 2 through 5). This analysis led to the removal of a total of 105 samples, such that the sample size of the experiment was at least n = 2 (four concentrations in total had n = 2; 41 had n = 3; and 70 had n = 4) for every concentration tested (Supplemental File 6A) with 18 DMSO controls per plate. Supplemental File 6B summarizes the number of controls pre- and post-filtering. Analyses to identify differentially expressed genes were conducted on probes using raw counts with the default parameters of DESeq2 v1.30. Here, default parameters include using Cook’s distance to suppress p-values for probes that had one or more samples deemed to be an outlier or an influential observation at the 0.99 cut- off. Time points were analyzed independently, as data across time points were generated from different sequencing runs. The full model used in each analysis included a treatment effect and a blocking effect indicating the index plate. Using the full model, pairwise comparisons were conducted for each chemical concentration combination to the appropriate DMSO control (i.e., matched to the relevant %DMSO). Probes reaching the threshold of a false discovery rate (FDR) adjusted p-value < 0.05 and an absolute fold change > 1.5 (on a linear scale) were flagged (probes passing filters) and retained for further analyses. Post-hoc collapsing of probes mapping to the same gene was conducted to generate the table of differentially expressed genes to represent unique genes only.

### Hierarchal clustering and principal component analysis of differentially expressed genes

Heatmaps were generated with the pheatmapfunction from R with both the row and column distances defined as 1-Spearman correlation. Values were averaged between replicates; CPM mean values of probes passing filters were averaged between replicates within each group. Resulting values were subsequently log2 transformed. The principal components were calculated with the prcompR function. The percentages of variation on the axes were gathered from the “importance” field produced by the summary function applied to the resulting object of the prcomp function. The work above was done with R v.3.6.1. Further details on pipeline analysis are available in Supplemental File 7. Principal component analysis plots were generated using the probes passing filters (Supplemental File 8A). This led to the identify of an additional DMSO outlier. The DeSeq2 analysis was repeated and the PCA was replotted (Supplemental File 8B; Supplemental File 6B has the total number DMSO controls used in the study). Hierarchical clustering plots for each time point are also included as Supplemental Files 8 C and 8D. Heatmaps were generated with the pheatmapfunction from R with both the row and column distances defined as 1- Spearman correlation with complete linkage. Note that the summary of the number of replicates (n) included for the benchmark concentration analysis are the same as those used to identify differentially expressed genes (Supplemental File 6A and B).

### Functional analysis using Ingenuity Pathway Analysis

Functional analysis of the differentially expressed genes was conducted using Ingenuity Pathway Analysis (IPA, Qiagen, Redwood City, CA, USA) using probes with FDR-corrected p-value < 0.05, absolute fold change > 1.5 and user supplied probes for background. For probes that mapped to the same gene, the probe with the maximum absolute fold change was used to represent the gene. Right-tailed Fisher’s exact tests were used to calculate the probability of enrichment of biological functions, canonical pathways, and upstream regulators. The background tab was set to “user-defined”, which allows for enrichment comparisons against the TempO-Seq panel only. The z-score, a statistical predictor of relationship between a direct observation and a prediction of activation/suppression of a particular pathway was set to +/- |2.0|, which is considered a significant cut-off (Ek *et al*., 2015).

### Benchmark concentration – Gene-level analysis

Best practices as recommended by the US EPA’s expert panel on genomic benchmark dose analysis were followed (National Toxicology Program Report, 2018). Benchmark concentration modeling was conducted using BMDExpress v.2.3, a free software package that is available for download (https://github.com/auerbachs/BMDExpress-2/releases). Additional information on BMDExpress is available (https://github.com/auerbachs/BMDExpress-2/wiki/Benchmark-Dose-Analysis). Normalized counts per million of the mapped reads for genes were used for this analysis. Samples identified to be cytotoxic or outliers were removed prior to benchmark concentration analysis. Data were imported into BMDExpress and subject to pre-filtering using the Williams’ Trend Test with 100 permutations and a 1.5-fold change (FC) cutoff to select genes that exhibit significant monotonically increasing or monotonically decreasing concentration-response (p < 0.05). Data exhibiting an increasing or decreasing change in response were modeled using Hill, Power, Linear, Polynomial (2° and 3°), and Exponential (2, 3, 4, 5) models to identify potential concentration-response relationships and best fit. For linear and polynomial models, the Nested Chi-square test was used to determine best-fit models followed by a comparison of Akaike information criterion (AIC) for nested models (Hill and Power) with a goodness of fit p-value of 0.1. Hill models were flagged if the ‘k’ parameter was less than 1/3 the lowest (non-solvent control) concentration, and in the case of a flagged Hill model, the next best-fit model with a goodness of fit p-value > 0.05 was selected (Hill *et al*., 2018). Additional parameters included a benchmark response factor of 1 standard deviation (SD) relative to the response of controls, a confidence level of 0.95, power restricted to ≥ 1, and maximum of 250 iterations (maximum iterations represent a convergence criterion for the model). Applied filters to best-fit models excluded data with highly uncertain benchmark concentrations (p < 0.1, BMC Upper threshold/ BMC Lower Threshold ≥ 40) as well as data producing benchmark concentration values > the highest non-cytotoxic concentration (e.g., 100 µM). Gene accumulation plots showing the distribution of gene-level benchmark concentrations across the range of concentrations for best-fit models were exported from BMDExpress. The BMD “project file (.bm2)” files are available through a link within the supplementary information (Supplementary File 9).

Potency comparison of the PFAS was conducted using the median benchmark concentration of all filtered genes adhering to best-fit models and a bootstrap R-script producing 95% confidence intervals. Potencies were also compared through visual inspection of gene accumulation plots. In addition, both the 5th percentile gene benchmark concentration and the lowest pathway median gene benchmark concentration were used to derive a bioactivity exposure ratio for PFOS and PFOA (described below).

### Derivation of the Bioactivity Exposure Ratio

Administered equivalent doses (AEDs) for PFOS and PFOA were estimated from toxicokinetics information described in Wambaugh et al. (2013). Wambaugh et al. performed simulations using a three compartmental model based on points of departure collected from various studies to obtain the steady-state concentrations (Css). Their population Monte Carlo simulation included a distributed oral uptake rate to consider for variability. We chose the Css and lowest observe effect level (LOEL) values for each of the PFOS and PFOA from non-cancer liver rodent points of departure, which were most consistent with the toxicokinetics analyses in this manuscript.

Transcriptional benchmark concentration values were converted to administered equivalent doses (AED) using the following equation:\

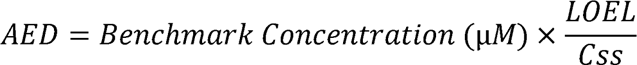

For ease of calculation, the formula was reduced to:

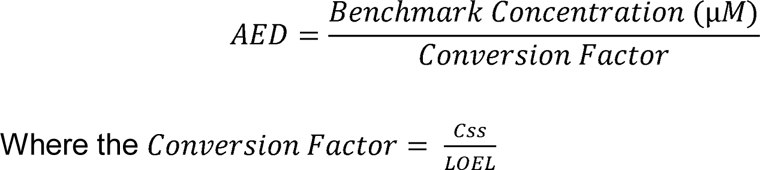

The resulting conversion factors to compute administered equivalent dose for each of the PFOS and PFOA are 19.47 and 46.4 mg/L per mg/kg/day, respectively.

To derive a bioactivity exposure ratio, the administered equivalent dose (mg/kg- BW/day) were compared to the upper limit of the daily US population exposure (95^th^ percentile) for PFOA (4.50×10^-04^ mg/kg-BW/day) and PFOS (4.76×10^-04^ mg/kg-BW/day) obtained from the US EPA comptox dashboard (https://comptox.epa.gov/dashboard, accessed: January 2020). The other PFAS within the current investigation (PFBS and PFDS) were excluded due to limited data availability.

Two approaches to bioactivity exposure ratio derivation were used. In **Approach 1**, the ratio was calculated using the lower bound 5^th^ percentile administered equivalent dose from the gene-level benchmark concentration analysis that was rounded to the nearest gene with an integer of greater than or equal value (i.e., the ceiling) and compared to the population exposure. This methodology has been used in previous work to represent a lower bound estimate of *in vivo* adverse effects within the ToxCast database in order to derive a bioactivity exposure ratio (Paul Friedman *et al*., 2020). In **Approach 2**, bioactivity exposure ratio was derived based on data demonstrating a strong correlation between the dose at which apical effects *in vivo* occur and the lowest (most sensitive) *in vivo* pathway benchmark concentration (Thomas *et al*., 2013). The pathway level analysis was conducted using the built-in Defined Category Analysis function within BMDExpress. Pathway-level estimates were derived from genes meeting the criteria of best-fit models mapped to probes from the IPA database (downloaded October 2019). Estimates were removed if they contained genes with a benchmark concentration > highest concentration from the category descriptive statistics, benchmark concentration with a goodness of fit p-value < 0.1, or a BMCU/BMCL value > 40. Exported pathway level estimates included additional filters limiting probe-sets to pathways with ≥ 3 genes passing all previous filters and with ≥ 5% of pathway genes affected. Each approach was then compared to reference administered equivalent dose values derived from *in vivo* animal data that were used to determine drinking water guidelines for PFOA and PFOS (Health Canada, 2018 a and b) for the purpose of risk assessment.

## Results

### Human primary liver cell spheroids microscopic evaluation

In order to assess the composition of the liver spheroid model, approximately 25 spheroids were stained with CD68, a marker of Kupffer cells, and phalloidin to stain for actin (Figure 1A). As anticipated, the spheroids stained positive for CD68, confirming the presence of this native cell type that is part of the normal liver composition. Actin staining showed the intact morphology of the spheroids and the hepatocytes within. The spheroids were likewise treated with a known inducer of steatosis, Cyclosporine A (CsA) for 48 hrs, and then processed for microscopic evaluation (Figure 1A). Treatment with 30 µM CsA induced expression of CYP3A4 (Figure 1B), a member of the cytochrome P450 oxidizing family of enzymes responsible for the oxidation of toxicants (Zanger and Schwab, 2013), and increased lipid accumulation as assessed by Nile Red staining (Figure 1B panel D). These results confirm that the spheroid hepatocytes were responding as expected.

**Figure 1.**
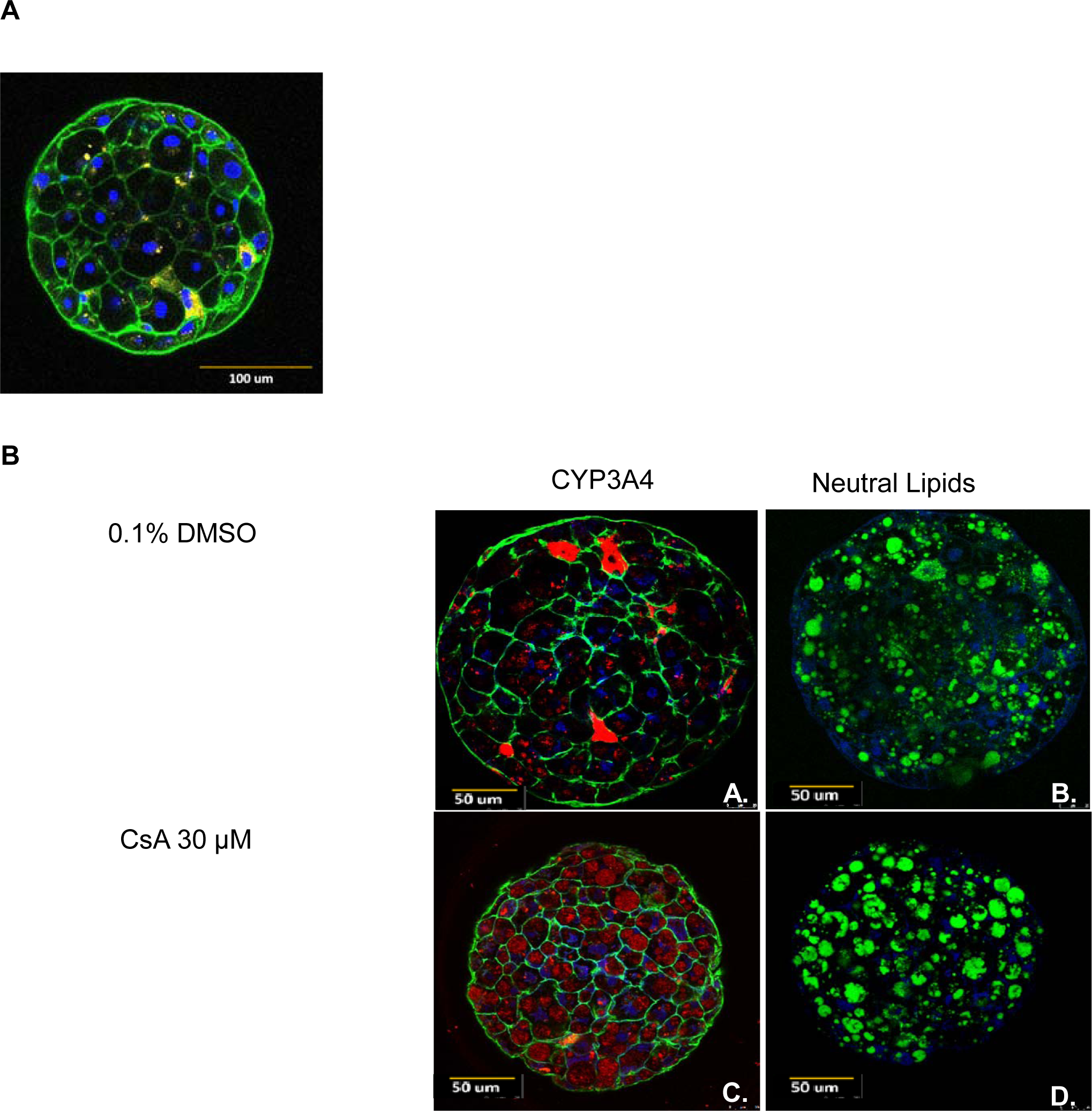
(A) Liver spheroid morphology after 4 days of treatment with 0.1% DMSO. Immunofluorescence staining with CD-68 for Kupffer cells (yellow), phalloidin for F-actin filaments (green), and DAPI for nuclei (blue), scale = 100 µm. (B) Effects of 48-hour exposure to Cyclosporine A on spheroid morphology compared to DMSO control. Microscopy images are reflective of expected detoxification and increased activity of the detoxification/biotransformation of the spheroids. (A and C) Immunofluorescence staining for: CYP3A4–(red), F-actin filaments stained with phalloidin (green), and nuclei with DAPI (blue). (B and D) Immunofluorescence staining for: neutral lipids (green), and nuclei with DAPI (blue). 400x magnification.

### Cytotoxicity assessment

The LDH-Glo assay was used to measure cytotoxicity over the course of the experiment. Cytotoxicity was evaluated at every media change (Days 3, 6, 9, 12, and 14). At each time point, concentrations that surpassed a threshold of 10-fold LDH release compared to the solvent controls were considered overtly cytotoxic. In addition to the LDH-Glo assay, cytotoxicity was evaluated based on read counts (TempO-Seq) and spheroid morphology (microscopy). Once LDH thresholds were surpassed, this cytotoxic concentration was eliminated from all time points thereafter.

Exposure of the liver cell spheroids to PFOA, the sole carboxylic acid of the group, did not induce significant cytotoxicity over time at 0.02 – 20 µM concentrations (Figure 2A). The 100 µM PFOA treatment resulted in an increase in LDH release in the supernatants beginning on day-6 (13.9-fold over control). By day-9, LDH levels decreased due to the decline in viable cells remaining. Data from the day-10 and day-14 time points of the 100 µM PFOA dose were excluded from further analyses based on these cytotoxicity results. The 50 µM concentration at day-14 was also eliminated because visual inspection revealed disintegration of the spheroids at this concentration and time point.

**Figure 2.**
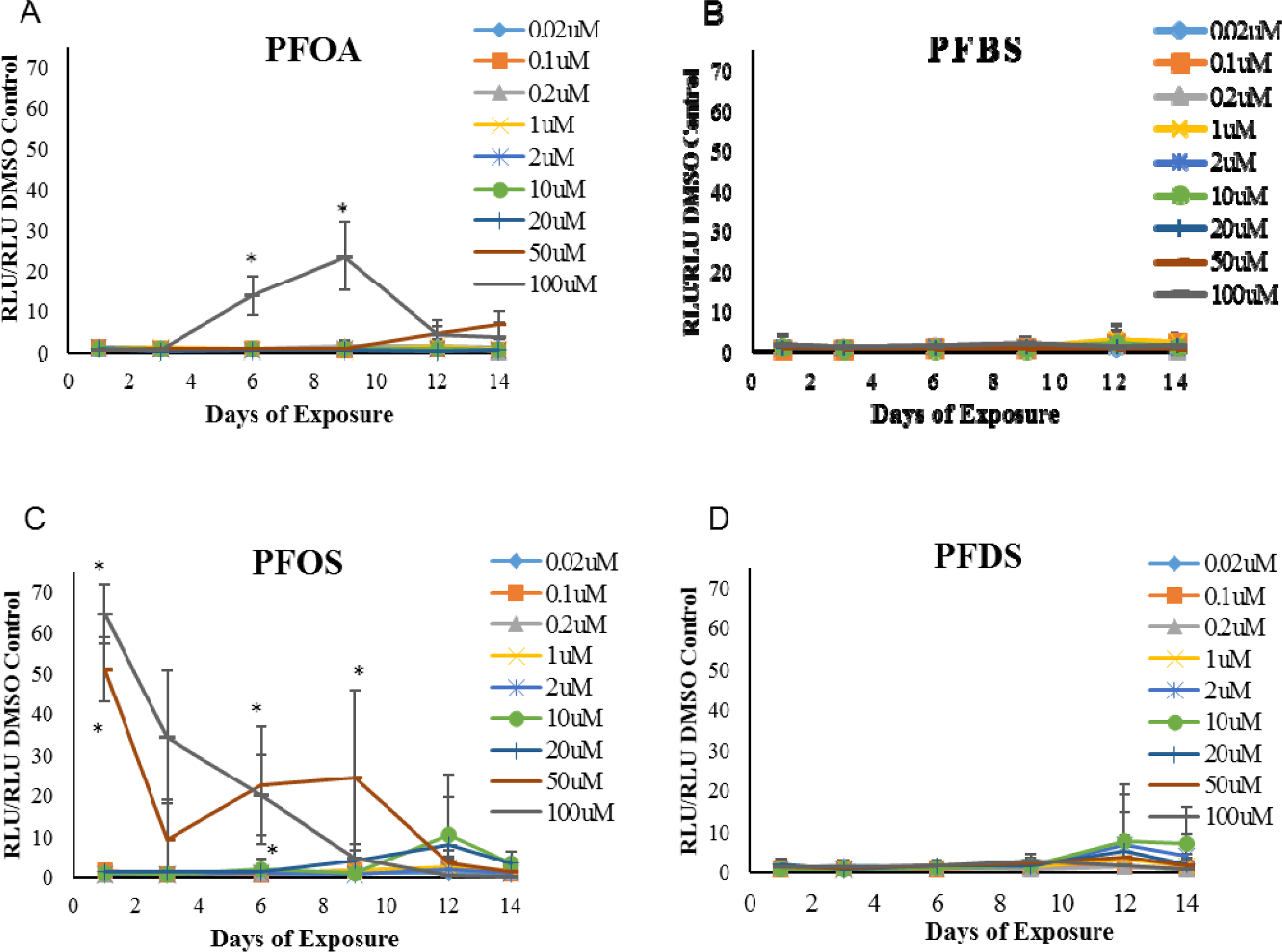
Lactate dehydrogenase (LDH) release as an indicator of cytotoxicity following treatment with a) PFOA, b) PFBS, c) PFOS, or d) PFDS over a 14-day time course (n = 4 with the exception of day-14, n =2). Plots demonstrate the means +/- standard deviation of the ratio of sample relative luminescence units (RLU) to mean DMSO RLU. * indicates the RLU was significantly higher than the DMSO control group (p < 0.05). Note that after reaching a peak, LDH levels decline over time consistent with reduced numbers of viable cells or evolving tolerance to chemical exposures in the spheroids.

PFOS was the most cytotoxic of the chemicals tested based on the magnitude of LDH leakage and the speed at which cell death occurred at the high concentrations (Figure 2C). The onset of cytotoxic effects occurred at 50 µM and 100 µM PFOS, causing dramatic cell death as early as 24 hrs post-exposure. The RLU ratios declined at later time points because of fewer viable cells as cell death progressed. After day-6, the LDH release for this concentration was no longer different from the control, due to the low number of surviving cells (visual qualitative inspection, data not shown). For the 50 µM PFOS treatment, an initial 50-fold increase in cytotoxicity was observed after day-1. Thus, the 50 µM and 100 µM concentrations were both excluded from the gene expression and benchmark concentration analysis. No overt cytotoxicity was observed for the shorter (PFBS) or longer (PFDS) chain PFAS (Figures 2B and 2D respectively).

### Identification of differentially expressed genes and hierarchical clustering

Table 1 summarizes the numbers of differentially expressed genes (FDR adj p-value ≤ 0.05 and |FC| ≥ 1.5) for each PFAS. As anticipated, the number of differentially expressed genes generally showed a trend toward increasing response with concentration. PFOS exposure induced the greatest number of differentially expressed genes; whereas, PFBS induced very few differentially expressed genes across time, even at the highest concentration (largest number of differentially expressed genes was 76 on day-10 at 100 µM). Comparing all PFAS at 20 µM, the highest non-cytotoxic concentration in common across all of the chemicals, revealed that the rank order for which PFAS induced the most transcriptional changes was consistent across time in the following order: PFOS > PFDS > PFOA > PFBS. There were minimal transcriptional alterations at the lowest concentrations; however, there was some indication for the possibility of non-monotonic responses (e.g., 85 differentially expressed genes for PFOS at 0.1 µM on day-1; and 30 genes for PFDS at 0.1 µM on day-10, Table 1). Note that a detailed pathway analysis of PFOS (presented below) over each time point is depicted within the supplementary information and suggests some differences in responses at lower (i.e., 0.1 µM) compared to higher (i.e., 10 and 20 µM) exposures over specific time points. Interpretation of these observations is beyond the scope of the current study but will be the subject of future investigation. Nevertheless, analysis of the differentially expressed genes for each PFAS at their top non-cytotoxic concentrations (Supplemental File 10 Panel A) and at a uniform concentration of 20 µM (Supplemental File 10B) showed few genes in common. Interestingly, Perilipin-2 (PLIN2), a lipid droplet associated protein involved in the storage of fats, appeared in several of the commonly affected genes.

**Table 1.**
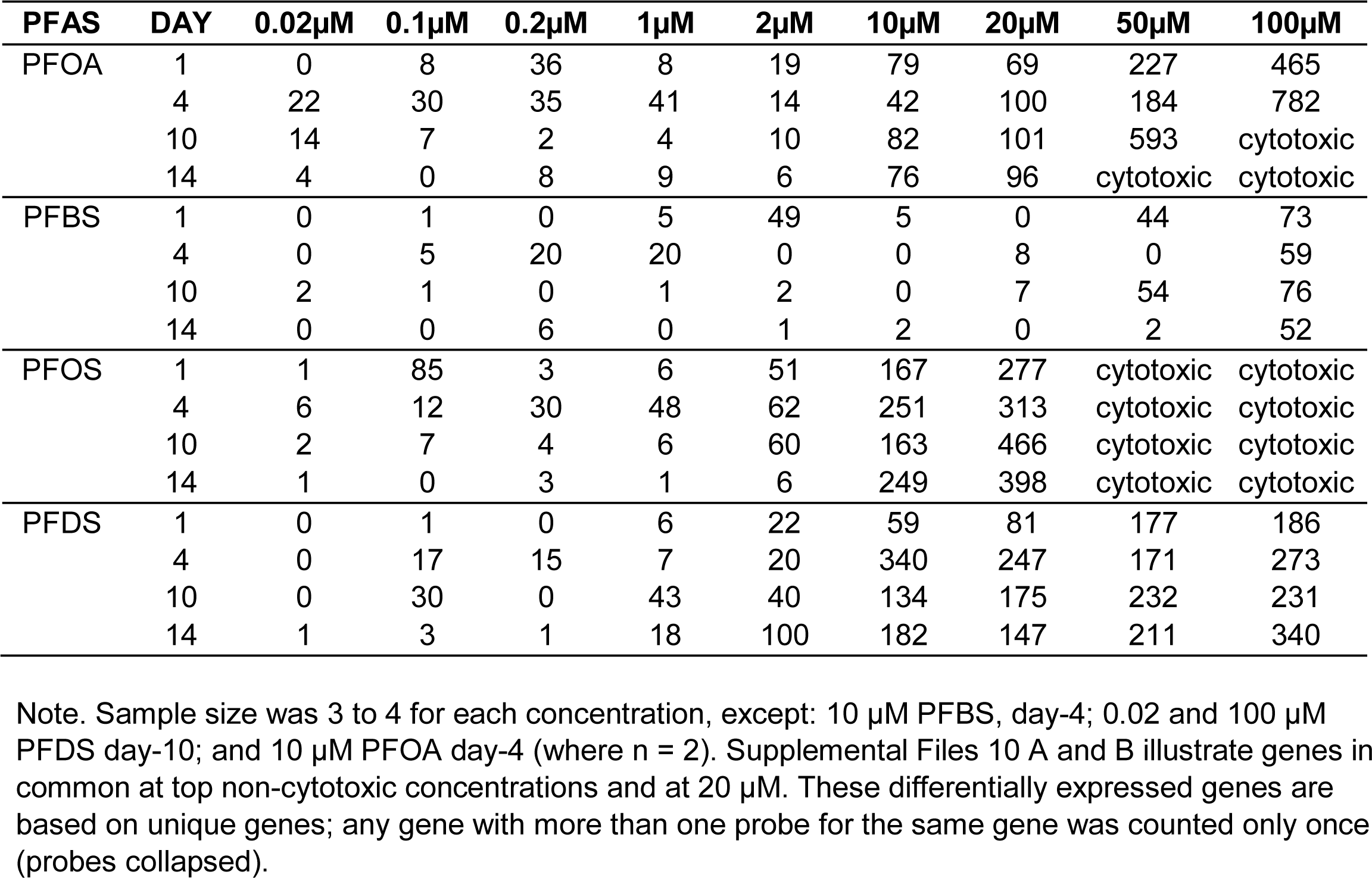
Number of differentially expressed genes in the PFAS-treated spheroids relative to controls for each time point and concentration.

| PFAS | DAY | 0.02µM | 0.1µM | 0.2µM | 1µM | 2µM | 10µM | 20µM | 50µM | 100µM |
| --- | --- | --- | --- | --- | --- | --- | --- | --- | --- | --- |
| PFOA | 1 | 0 | 8 | 36 | 8 | 19 | 79 | 69 | 227 | 465 |
|  | 4 | 22 | 30 | 35 | 41 | 14 | 42 | 100 | 184 | 782 |
|  | 10 | 14 | 7 | 2 | 4 | 10 | 82 | 101 | 593 | cytotoxic |
|  | 14 | 4 | 0 | 8 | 9 | 6 | 76 | 96 | cytotoxic | cytotoxic |
| PFBS | 1 | 0 | 1 | 0 | 5 | 49 | 5 | 0 | 44 | 73 |
|  | 4 | 0 | 5 | 20 | 20 | 0 | 0 | 8 | 0 | 59 |
|  | 10 | 2 | 1 | 0 | 1 | 2 | 0 | 7 | 54 | 76 |
|  | 14 | 0 | 0 | 6 | 0 | 1 | 2 | 0 | 2 | 52 |
| PFOS | 1 | 1 | 85 | 3 | 6 | 51 | 167 | 277 | cytotoxic | cytotoxic |
|  | 4 | 6 | 12 | 30 | 48 | 62 | 251 | 313 | cytotoxic | cytotoxic |
|  | 10 | 2 | 7 | 4 | 6 | 60 | 163 | 466 | cytotoxic | cytotoxic |
|  | 14 | 1 | 0 | 3 | 1 | 6 | 249 | 398 | cytotoxic | cytotoxic |
| PFDS | 1 | 0 | 1 | 0 | 6 | 22 | 59 | 81 | 177 | 186 |
|  | 4 | 0 | 17 | 15 | 7 | 20 | 340 | 247 | 171 | 273 |
|  | 10 | 0 | 30 | 0 | 43 | 40 | 134 | 175 | 232 | 231 |
|  | 14 | 1 | 3 | 1 | 18 | 100 | 182 | 147 | 211 | 340 |

Note. Sample size was 3 to 4 for each concentration, except: 10 µM PFBS, day-4; 0.02 and 100 µM PFDS day-10; and 10 µM PFOA day-4 (where n = 2). Supplemental Files 10 A and B illustrate genes in common at top non-cytotoxic concentrations and at 20 µM. These differentially expressed genes are based on unique genes; any gene with more than one probe for the same gene was counted only once (probes collapsed).

Principal component analysis was conducted on the differentially expressed genes at each time point to explore general relationships in the transcriptional profiles induced by the PFAS (Figure 3; Supplemental File 8B). Generally, principal component 1 (the main driver of variability) appears to be based on PFAS concentration, with higher concentrations to the right on the plot and lower concentrations together on the left. The analysis demonstrates the more marginal impact of PFBS on the liver spheroid transcriptome. The high concentrations of PFBS tend to group more closely with the low concentrations and controls (in particular after day-1). In addition, higher concentrations of PFOS and PFDS were more frequently in close proximity to each other than they were to PFOA, suggesting more similar profiles. However, by day-14, principal component 2 separates PFOS and PFDS. Note that the top concentration here for both PFOS and PFOA is 20 µM.

**Figure 3.**
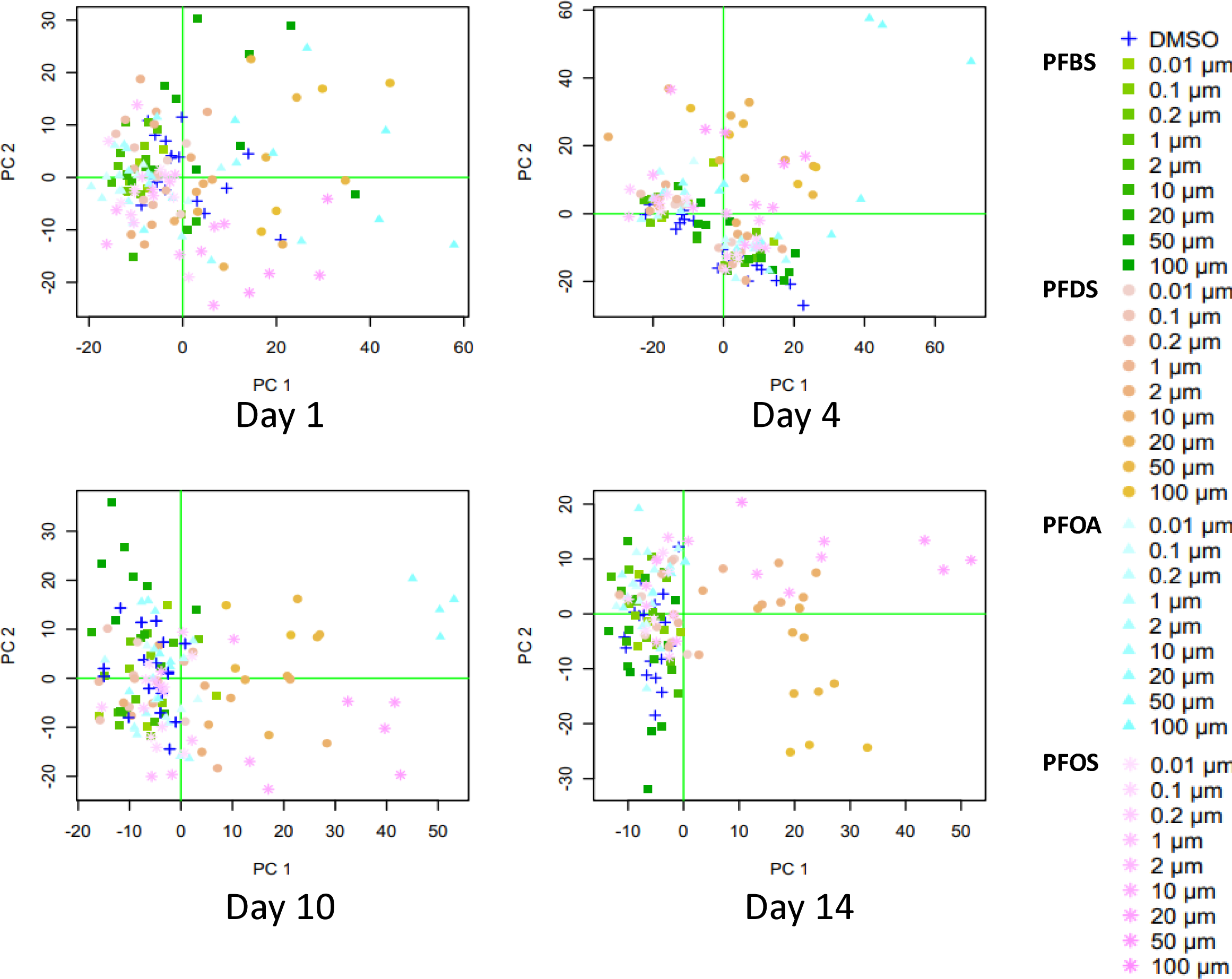
Principal Component Analysis conducted on the probes passing filters (FDR adjusted p-value < 0.05 and an absolute fold change > 1.5 on a linear scale) at each of the four time points. Each point represents the individual replicates. The horizontal and vertical green lines at zero were added to identify the four quadrants in each of the four plots. Sample size was n = 3-4 for all exposure groups, except for n = 2 on day-4 PFBS 10 µM and PFOA 10 µM, and n = 2 on day-10 PFDS 0.02 and 100 µM. Additional principal component analyses explored each chemical individually within a time point (Supplemental File 8A, B). Hierarchical cluster analysis was also performed for each time point and can be found in Supplemental Files 8C, and 8D.

### Functional analysis and upstream regulators

Pathway (i.e., gene set) analyses were conducted in IPA using the differentially expressed genes from the 20 µM concentration (selected because it was the top non-cytotoxic concentration in common across all the PFAS) (Figure 4A), or the top non-cytotoxic concentration (Figure 4B), for each chemical and time points. This analysis revealed that most pathways affected by PFAS were downregulated based on IPA predictions that factor in the directionality of the expression changes. Overall, there was a remarkably similar pattern of pathway enrichment for PFOS and PFDS that was different from PFOA and PFBS.

**Figure 4.**
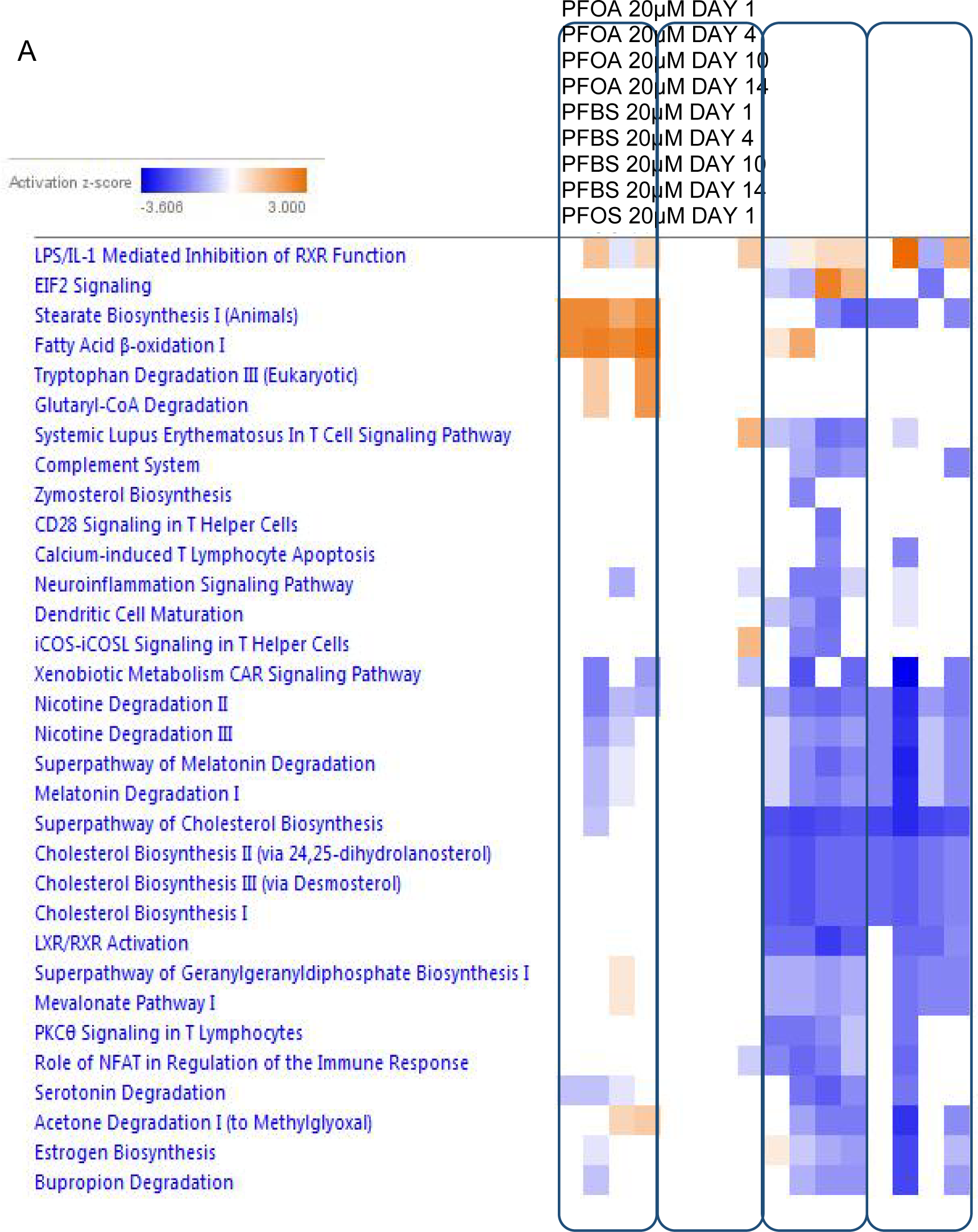
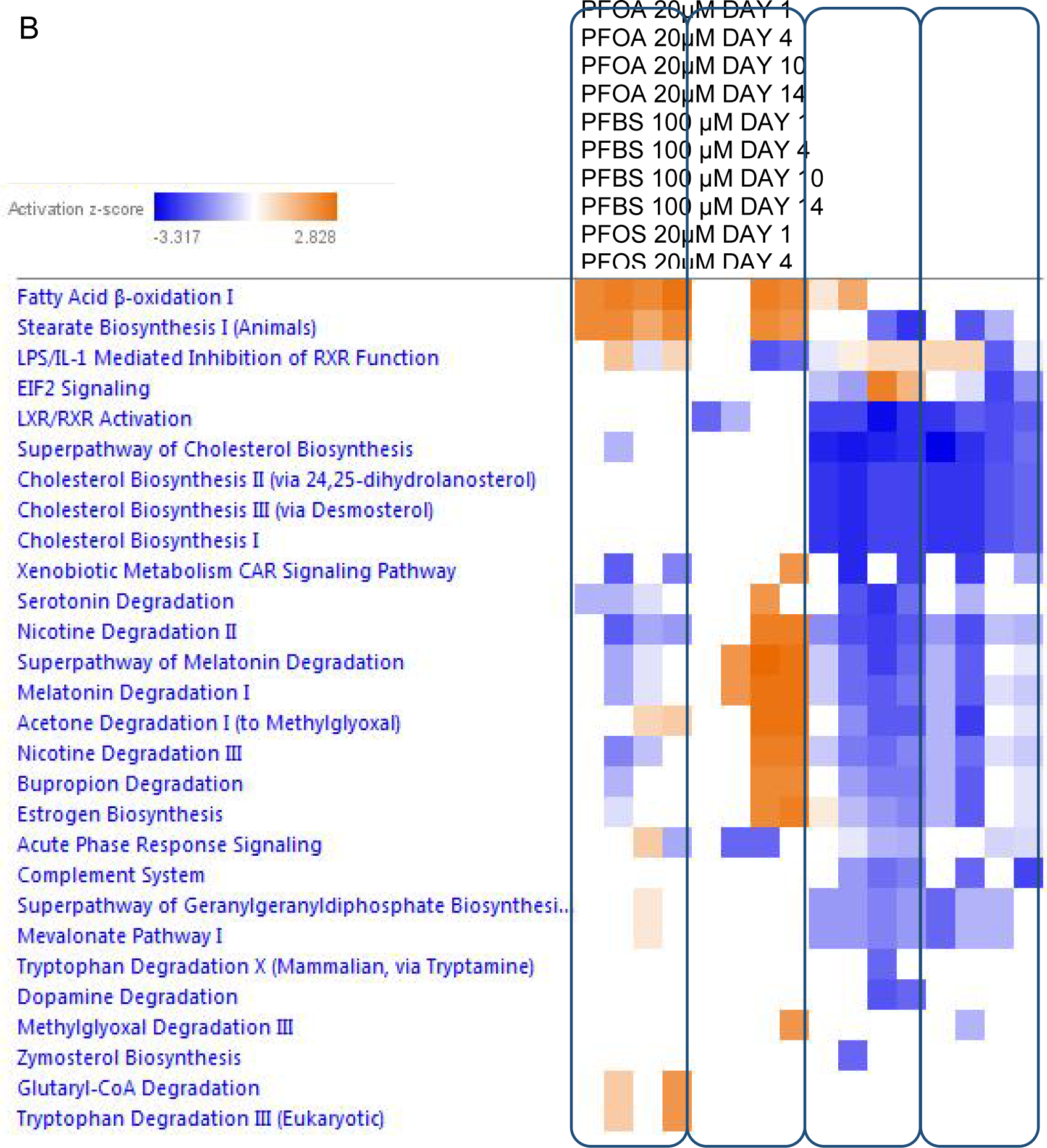
Pathway analysis of the differentially expressed genes for all time points at the 20 µM concentration (A) or for the highest non-cytotoxic concentrations (B). For (B) concentrations were 20 µM for PFOA and PFOS, and 100 µM for PFBS and PFDS. Filters for this analysis were set to z-score ≥ absolute 2.0 and Benjamin–Hochberg adjusted p-value ≤ 0.05. Blue represents inhibited pathways and orange represents activated pathways. n = 3-4 for all exposure groups, except for n = 2 on day-10 PFDS 100 µM. Supplemental File 11 A & B is a less stringent pathway analysis for all PFAS over all time points (no z-score applied). Supplemental File 12 illustrates a more detailed pathway analysis for PFOS over all time points and concentrations.

PFOS and PFDS pathway analyses suggest that the exposures cause inhibition of nicotine and melatonin degradation pathways as well as downregulation of cholesterol biosynthesis pathways and LXL/RXR activity, which all appear to persist across time. In contrast, PFOA treatment appears to strongly activate stearate biosynthesis and fatty acid beta oxidation pathways. Unlike the other PFAS, PFBS displayed such small proportions of gene expression changes that it did not perturb many canonical pathways up to 20 µM. However, the 100 µM concentration of PFBS elicited a strong upregulation of transcript in several pathways at the 10 and 14-day time points (Figure 4B) including fatty acid oxidation, stearate biosynthesis, nicotine degradation, melatonin degradation, acetone degradation, estrogen biosynthesis, and bupropion degradation. Supplemental File 11 A & B illustrates a less stringent pathway analysis for all PFAS over all time points (no z-score applied).

A detailed overview of perturbed pathways over all concentrations and time points for each PFAS is beyond the scope of this study, which is geared toward analysis of general trends/patterns across the data. However, concentration-response for the pathway analysis is shown for PFOS as an illustration (Supplemental File 12). This analysis supports the general trend of increasing pathway perturbations at higher concentrations (with the exception of 0.1 µM on day-1), and consistency in response of specific pathways across concentration and time including inhibition of cholesterol biosynthesis, LXR/RXR activation, melatonin degradation, and nicotine degradation.

The upstream analysis was restricted to more stringent criteria (z-score filter of absolute +/- |3.0| and a Benjamini Hochberg adjusted p-value of 0.05) to focus on the most impacted regulators representing potential key mediators of the differentially expressed genes and to provide additional insight into mode of action. Peroxisome proliferator-activated receptor alpha (PPARα) was predicted to be an activated upstream regulator for all four PFAS at all time points (Figure 5). Furthermore, PPARα was predicted to have increased activity in all PFAS exposures to varying degrees. Of the predicted inhibited upstream regulators, Copper-transporting ATPase 2 (ATP7B), sterol regulatory element-binding transcription factor 1 and 2 (SREBF 1 and SREBF 2), sterol regulatory element-binding protein cleavage-activating protein (SCAP), and

**Figure 5.**
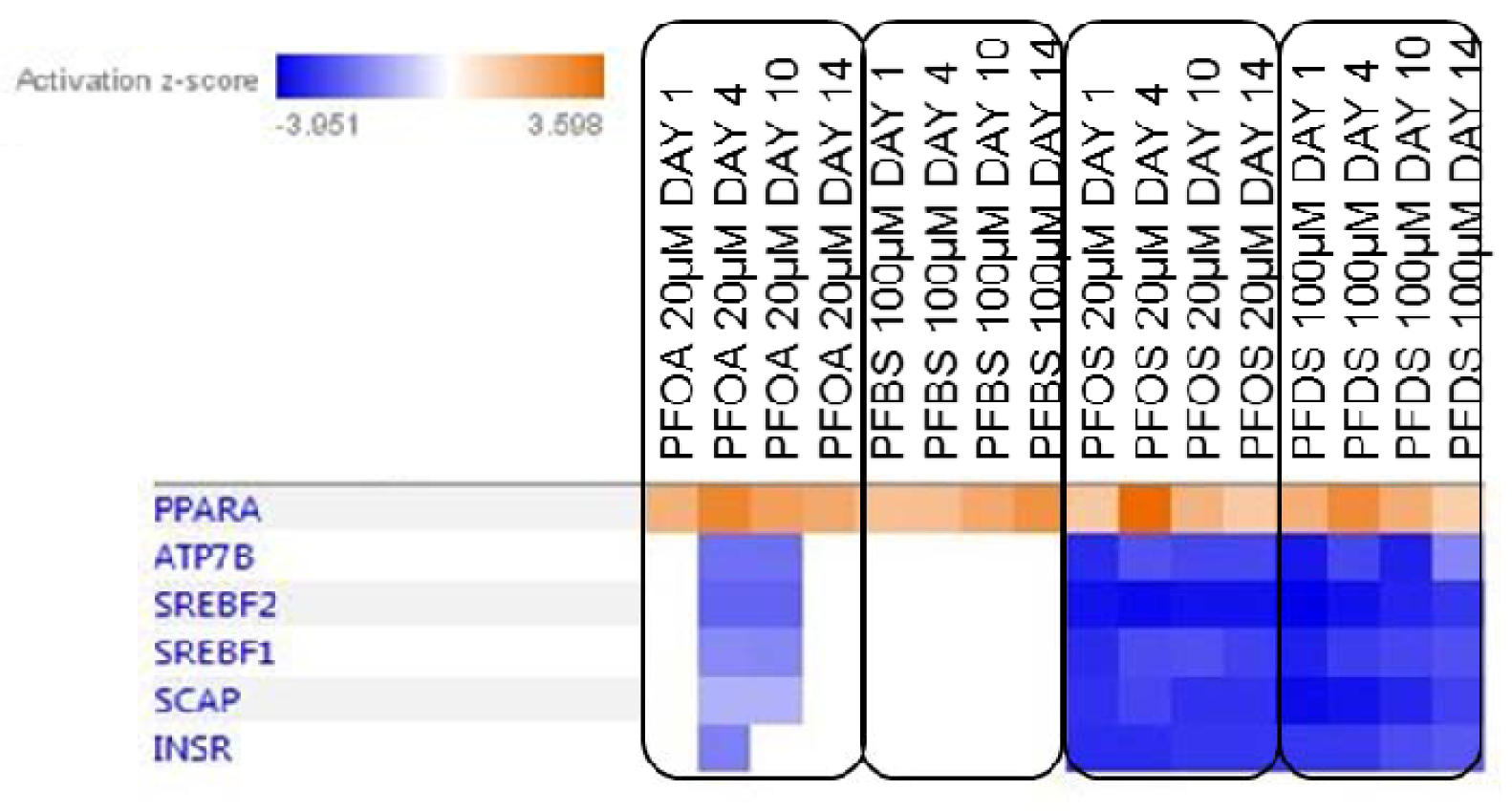
Upstream regulator predictions from the differentially expressed genes for the top non-cytotoxic concentration of each PFAS. The z-score filters were set to 3.0 with a Benjamini Hochberg adj p-value of 0.05 for this analysis. Blue represents inhibited upstream regulators and orange represents activated regulators. n = 3-4 for all exposure groups, except for n = 2 on day-10 PFDS 100 µM.

Insulin Receptor (INSR) were all predicted to be inhibited at all-time points with PFOS and PFDS. Interestingly, these upstream regulators were also predicted to be inhibited (albeit more weakly) by PFOA at the day-4 and day-10 time points. Overall, there was a strikingly consistent pattern of upstream regulator activation/inhibition between PFOS and PFDS that differed from the other two PFAS. Although this stringent approach limited the numbers of regulators analyzed, it was a useful to begin examining the PFAS targets.

### Benchmark concentration analysis

To compare potencies across the PFAS, we conducted a transcriptional benchmark concentration analysis with a benchmark response of one standard deviation. To broadly compare potencies, median benchmark concentrations for all genes that could be obtained from best-fit models within BMDExpress were derived along with a bootstrapped 95% confidence interval (Figure 6). An overview of the proportion of genes that were best fit to each model analyzed is shown in the supplementary information (Supplemental File 13). PFOS consistently had a median across all time points. There was some variability across time in the benchmark concentrations of the other PFAS. The median benchmark concentrations of both PFOA and PFDS on day-1 were significantly higher than PFOS (non-overlapping confidence intervals); however, by day-4 (for PFDS), and day-14 (for PFOA), both of these PFAS were within the same range as PFOS, suggesting similar potencies. In contrast, PFBS was significantly less potent with benchmark concentrations ranging from 46.3 to 69.7 µM (Supplemental File 14).

**Figure 6.**
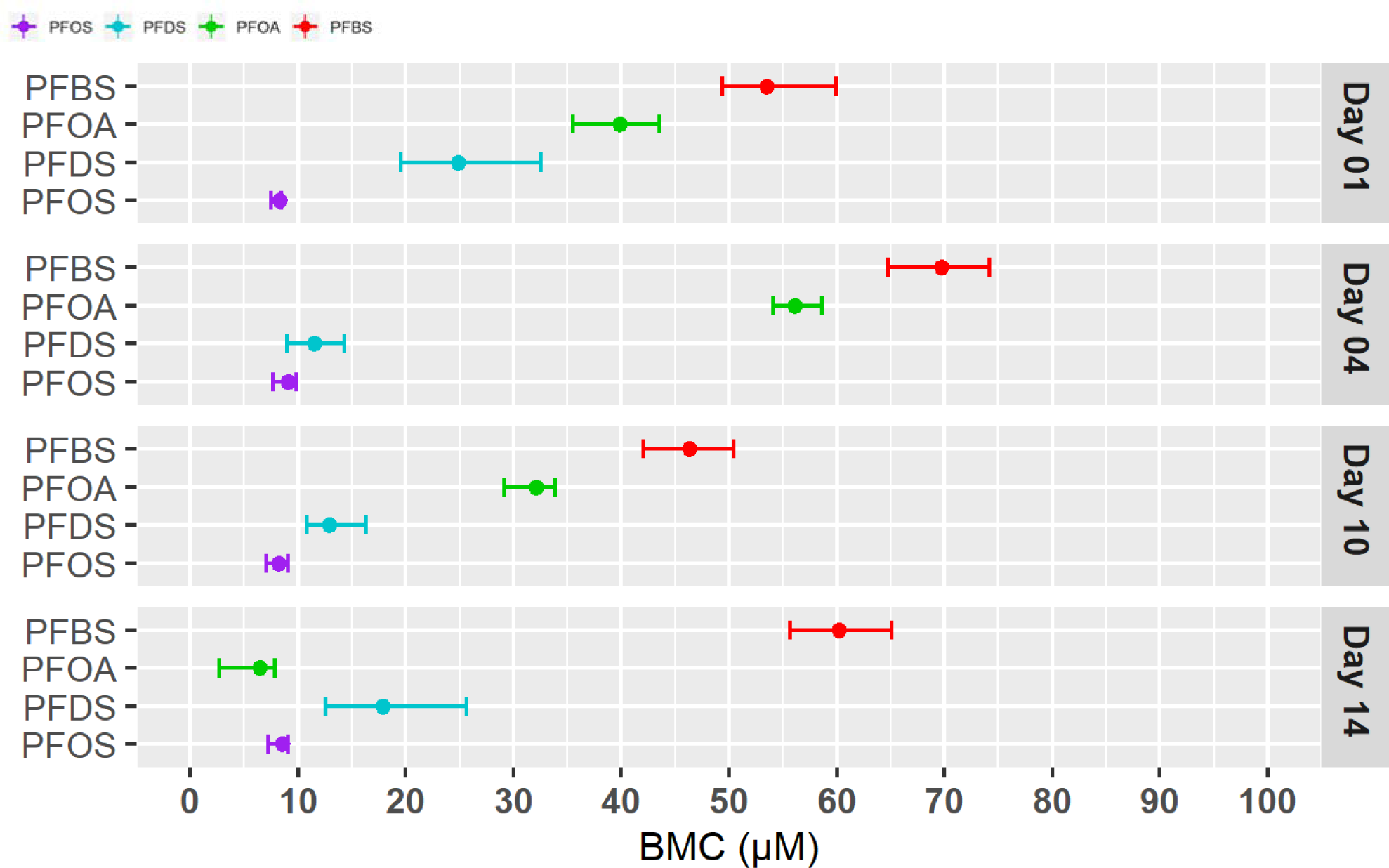
Median overall gene benchmark concentration (BMC) (colored dot) for all the genes that could be modeled and 95% confidence intervals for each of the PFAS. n = 3-4 for all treatment groups, except for n = 2 on day- 4 for PFBS 10 µM and PFOA 10 µM, and day-10 PFDS 0.02 and 100 µM. Supplemental File 14 shows overall benchmark concentrations (95 % CI) by day. A summary of the distribution for all of the modeled genes exposed to PFAS over 1, 4, 10, and 14 days is in Supplemental File 13, and a summary of sample replicates is in Supplemental File 6.

Gene accumulation plots show the benchmark concentrations for each of the genes that could be modeled (Figure 7). Note that overtly cytotoxic concentrations were not modeled; thus, some curves terminate at lower concentrations than others (i.e., 20 µM for all PFOS time points, and 20 µM for PFOA for day- 14). The plot demonstrates that of those genes that could be modeled, the transcriptional benchmark concentrations for PFOS were generally lower overall than the other PFAS (i.e., shifted furthest to the left on the x-axis). Similar to the overall BMC median gene comparisons above, PFDS and PFOS accumulation curves in the low concentration ranges are more overlapping by day-4 and later, suggesting similar potencies. By comparison, PFBS transcripts had higher benchmark concentrations relative to the other three PFAS, indicating that it was the least potent in inducing transcriptional changes (most right shifted). Notably, transcriptional activity for PFOS, PFOA, and PFDS converge at the origin, indicating that the ‘most sensitive’ gene transcripts are induced in a similar concentration range.

**Figure 7.**
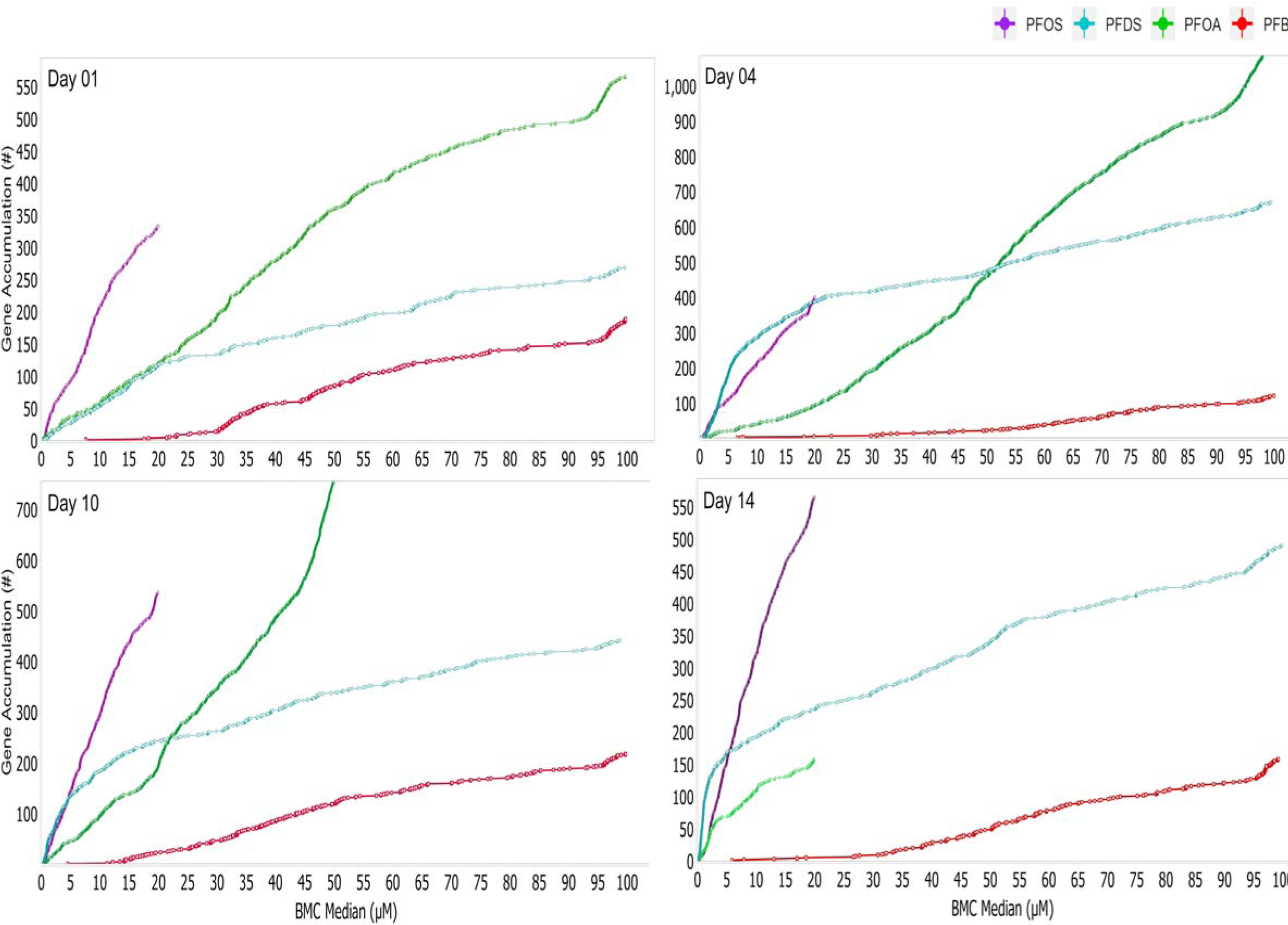
Gene accumulation plots showing the benchmark concentrations (BMC) for each gene that could be modeled for each of the PFAS across time points. n = 3-4 for all exposure groups, except for n = 2 for day-4 PFBS 10 µM and PFOA 10 µM and day-10 PFDS, 0.02 and 100 µM. Supplemental File 14 summarizes the benchmark concentrations (and 95 % CI) by day. A summary of the distribution for all of the modeled genes exposed to PFAS over 1, 4, 10, and 14 days is in Supplemental File 13, and a summary of sample replicates is in Supplemental File 6.

### Bioactivity Exposure Ratio

The bioactivity exposure ratio (BER) is the ratio of the administered equivalent dose for a new approach methodology (NAM) point of departure to the daily population exposure for a given chemical. A large bioactivity exposure ratio value is an indication of lower risk, whereas a lower value indicates potentially increased risk to human health (Wetmore *et al*., 2015b; Paul Friedman *et al*., 2020). The use of transcriptomics in risk assessment is a relatively new concept and there is lack of consistency in how to derive the transcriptomic point of departure. Recognizing this lack of standardization, Farmahin *et al*., (2017) demonstrated general consistency (within 10x) across a variety of methods used for transcriptomic point of departure determination. Herein we compare two of the most conservative approaches (applying precautionary practices) for deriving a transcriptomic point of departure. For **Approach 1**, the applied point of departure is comparable to those approaches used previously for ToxCast data: the 5^th^ percentile gene benchmark concentration (Paul Friedman *et al*., 2020). **Approach 2** was based on data demonstrating a strong correlation between the dose at which apical effects *in vivo* occur and the lowest (most sensitive) *in vivo* pathway benchmark concentration (Thomas *et al*., 2013). Bioactivity exposure ratios were derived by dividing the administered equivalent dose (for each transcriptomic point of departure approach) by exposure estimates taken from the US population (Figure 8). The bioactivity exposure ratios for each approach (approach 1, blue diamond; approach 2, red diamond) were then compared to a range of endpoints derived from animal exposure data used in risk assessment guidelines (green crossbar) at each time point for PFOA and PFOS (Table 2). As previously noted, PFBS and PFDS were excluded from bioactivity exposure ratio calculation due to lack of available data.

**Figure 8.**
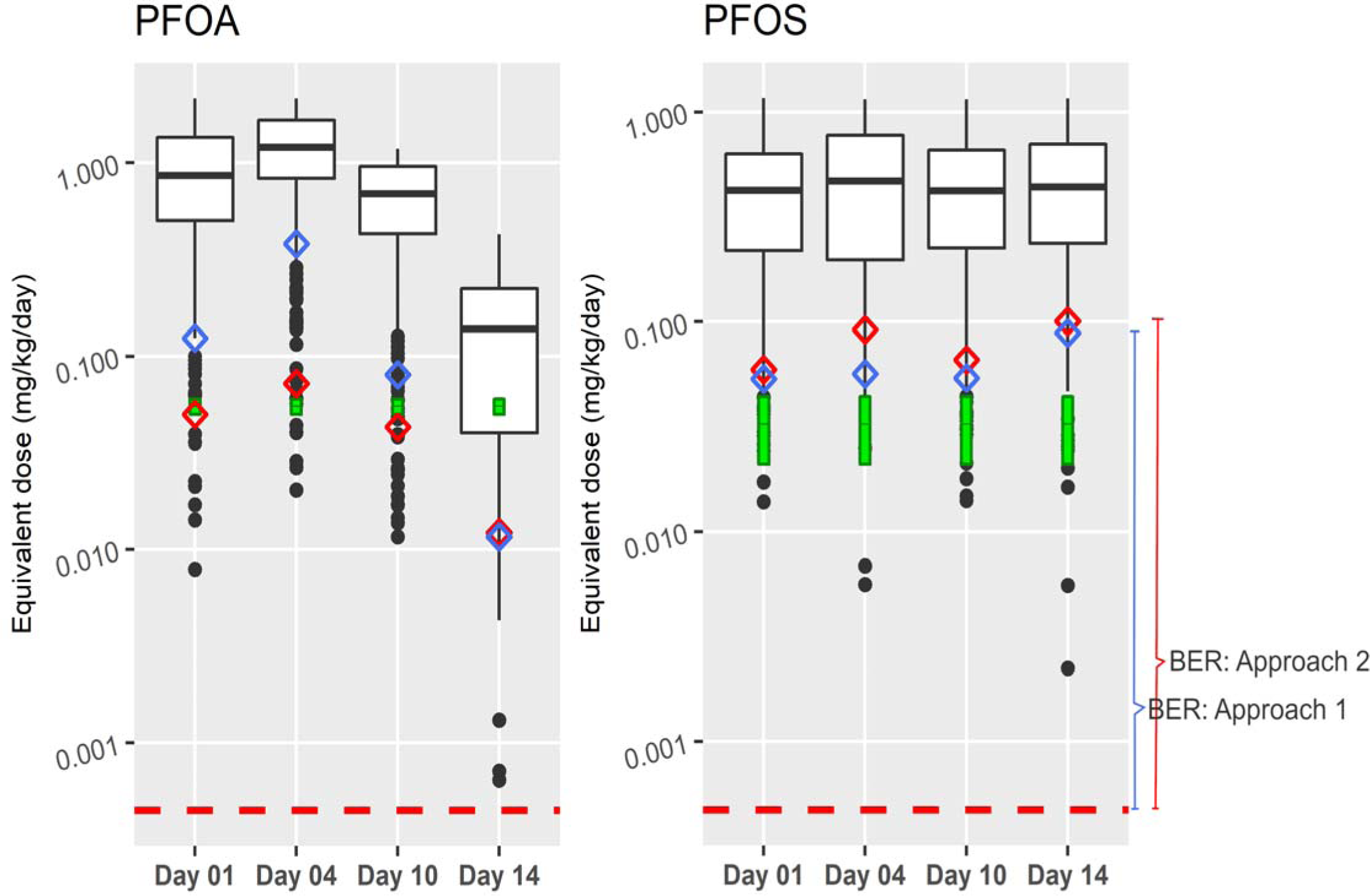
Boxplots of administered equivalent doses (mg/kg-BW/day) of all transcripts modeled for PFOA and PFOS are shown. These were used to determine the Bioactivity Exposure Ratio (BER) based on either the lowest 5th percentile administered equivalent dose (blue diamond) or the median administered equivalent dose of the gene-set from the most sensitive pathway (red diamond). These were compared with the estimated daily exposure of the upper limit of the daily US population exposure (95th percentile) obtained from the US EPA CompTox dashboard (red dash). A range of administered equivalent doses was included as a representation of apical points of departure that were derived from non-cancer animal in vivo data (green crossbar) from Health Canada drinking water guidelines (Health Canada 2018a, and 2018b).

**Table 2.**
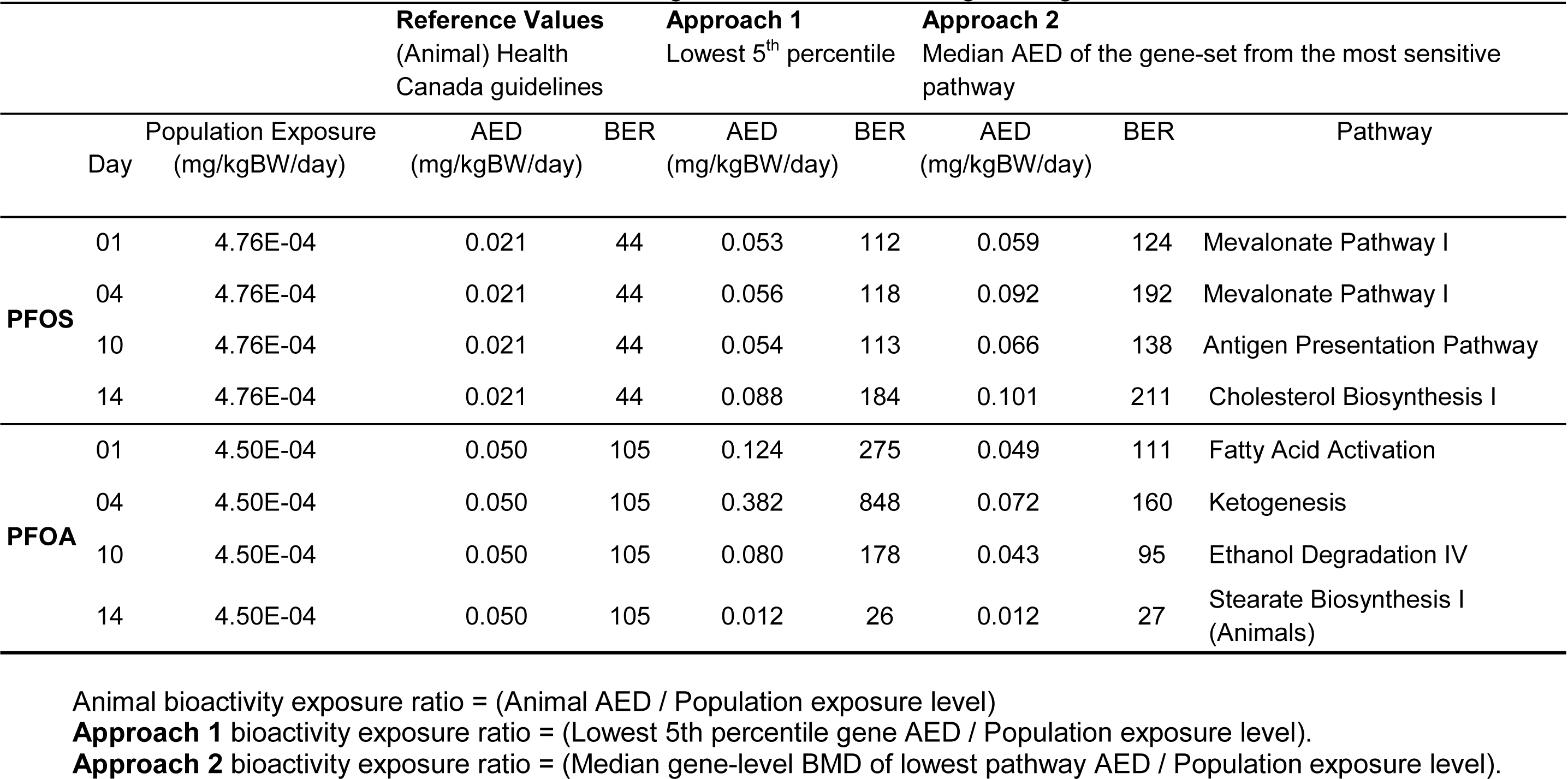
Administered Equivalent Dose (AEDs) and bioactivity exposure ratio (BER) derived from Approaches 1 and 2 as well as reference values from animal models using Health Canada drinking water guidelines for PFOA and PFOS.

For PFOA, administered equivalent dose values slightly increased from day-1 to day-4 and were lowest by day-14 for both point of departure approaches (Figure 8). The calculated bioactivity exposure ratios for approach 1 were within 2-orders of magnitude of the population exposure, with a range 178 to 848, except for day-14 with a bioactivity exposure ratio of 26 (Table 2). The second approach generally decreased the bioactivity exposure ratios for longer treatment times (day-10 and day-14 bioactivity exposure ratios of 95 and 27, respectively) compared to shorter treatment times (day-1 and day-4 bioactivity exposure ratios of 111 and 160, respectively) (Table 2). The reference bioactivity exposure ratio for PFOA (105) was derived using the lowest endpoint in Health Canada’s drinking water guidelines: the BMD_10_ for induction of hepatocellular hypertrophy in rats (Health Canada, 2018a). This animal *in vivo* bioactivity exposure ratio was within the same order of magnitude for days 1, 4, and 10, but was about 4x higher than the bioactivity exposure ratios for both approaches 1 and 2 on day-14 (Table 2).

For PFOS, administered equivalent dose values remained relatively constant over treatment time for both approaches (Figure 8). In addition, the bioactivity exposure ratios (Table 2) for approach 1 (range of 112 to 184) were highly similar to approach 2 (range 124 to 211), with approach 1 being marginally more conservative. The reference bioactivity exposure ratio for PFOS (44) was derived using the lowest endpoint from Health Canada’s drinking water guidelines: the no-observed-adverse-effect-level (NOAEL) for induction of hepatocellular hypertrophy in rats (Health Canada, 2018b). This animal *in vivo* data-derived bioactivity exposure ratio was within an order of magnitude of both transcriptomic points of departure, but was consistently lower than the NAM-based ratio.

## Discussion

This study investigates the utility of a primary human liver cell spheroid model paired with transcriptional profiling as an alternative approach to toxicological assessment of PFAS. Four prototype PFAS were examined using a detailed concentration-response analysis across four time points. The extent of transcriptional changes induced by PFOS was larger than any of the other PFAS, with transcriptional perturbations at matched concentrations in the order PFOS > PFDS > PFOA > PFBS. Although PFOS had the lowest benchmark concentrations at the early time points, the transcriptomic benchmark concentrations of PFOS, PFDS, and PFOA were overlapping at the later time points, indicating the importance of transcriptional time-series analyses in potency comparisons. In addition, the lowest transcriptomic benchmark concentrations were highly similar across these three PFAS, suggesting fairly equal potencies. Pathway and upstream regulator analysis revealed that although PFOS and PFDS transcriptional profiles were most similar to each other, all of the PFAS appeared to activate PPARα. Indeed, PFOS and PFDS induced remarkably similar transcriptional α alterations at the pathway level. In contrast, the short-chain PFAS (PFBS) induced marginal changes in transcriptional activity and was much less potent. These data serve as a foundational data set for comparison of additional data-poor PFAS and for developing approaches in this human liver cell model in future studies. Functional analysis of the differentially expressed genes suggests that PPARα plays a role in mediating the toxic effects of PFAS in human liver spheroids. PPARα was predicted to be an ‘activated’ upstream across all of the PFAS and time points. PPARs are nuclear receptors involved in the regulation of genes involved in energy homeostasis and lipid metabolism (Janssen *et al*., 2015; Liss and Finck, 2017). Dose-dependent increases in PPARα-regulated genes with PFOS exposure have been observed in rodents (Ye et al., 2012). Likewise, increased PPARα activity in rodents is associated with exposure to both PFOA and PFOS (Rosen *et al*., 2008, 2010). However, these studies also reveal that PPARα knock-out mice have altered lipid metabolism pathways following PFOS and PFOA exposure, similar to the wild type animals, indicating PPARα -independent mechanisms as well (Rosen et al., 2008, 2010; Abbott et al., 2009). Our data support that PPARα activity is increased in response to PFAS exposure in human liver spheroids, but that there are other upstream regulators that are likewise affected.

PFOA, PFOS, and PFDS induced changes in gene expression in our study that are suggestive of inhibition of several upstream regulators: ATP7B, SREBF1 and 2, SCAP, and INSR. Inhibition of ATP7B, an ATPase copper protein transporter required for the transport of copper to the bile, is associated with copper accumulation in hepatocytes (Bartee and Lutsenko, 2007). Inhibition of sterol regulatory binding factors 1 and 2 (SREBF 1 and 2), and sterol regulatory element-binding protein cleavage – activating protein (SCAP), which are both a part of the cholesterol, fatty acids, and triglycerides synthesis pathways (Moslehi and Hamidi-zad, 2018), suggests downregulation of lipogenesis. Perturbation of these pathways is potentially adaptive; i.e., as PPARα is activated, SREBP1, 2 and SCAP may be downregulated in an attempt to reach homeostasis of lipid accumulation. Finally, suppression of the insulin receptor (INSR), an activator of the SREBF 1 transcription factor, suggests that PFAS exposure affects the insulin signalling pathway and thus impairs normal lipogenesis further. Over time, continued disruption of cholesterol efflux into the intestine may ultimately lead to liver disease (Vrins *et al*., 2012; Utzschneider and Kahn, 2006).

PFOA exhibited a broadly different pattern of transcriptional pathway alterations at the concentrations examined than the other PFAS. PFOS and PFDS exerted major effects on the transcription of genes involved in cholesterol biosynthesis. In contrast, PFOA upregulated transcripts involved in the stearate biosynthesis I pathway, in parallel with the fatty acid-ß oxidation pathway. Das *et al*. (2017) found that exposure of wild type and PPARα-null mice to PFOA resulted in increased fatty acid accumulation in the liver, which is aligned with our findings. Upregulation of the fatty acid-ß oxidation pathway after PFOA exposure has also been reported in Balb/c mice (Yu *et al*., 2016) and in Wistar rats (Haughom and Spydevold, 1992). Although upregulation of the stearate biosynthesis pathway, at the same time as upregulation of the oxidation or breakdown of fatty acids, seems counterproductive, we speculate that PFOA is causing dysregulation of pathways involved in lipid homeostasis in the liver. Overall, although fairly equal in transcriptional potency, PFOA appears to operate through different mechanisms to PFOS and PFDS, suggesting that it may be advisable to consider this carboxylate separately from PFAS with sulfonic acid functional groups.

PFBS is clearly less potent at inducing transcriptional changes than the other PFAS. Indeed, only the highest concentrations of PFBS and later time points had significant transcriptional activity. Our results are consistent with an analysis by Naile *et al*. (2011) of 10 PFAS in rat hepatocytes that revealed that shorter chain PFAS cause less transcriptional perturbation.

Our transcriptomic findings are also consistent with demonstrated effects of PFAS on cholesterol, lipid synthesis, and metabolism in other model organisms. Although data for most PFAS are limited, rodent exposure to PFOS and PFOA is known to induce changes in fatty acid production, cholesterol synthesis, and lipid homeostasis leading to cholestasis/steatosis (Wang *et al*., 2014; Bijland *et al*., 2011; Du *et al*., 2009; Peden-Adams *et al*., 2009; Wan *et al*., 2012). In keeping with these effects, obese participants from the National Health and Nutrition Examination Survey (2011-2014) exhibit a positive association of PFOA, PFHxS, and PFNA exposure with major liver biomarkers of steatosis (Jain and Ducatman, 2019). These findings support the relevance of our results for exposed humans. Overall, our short-term *in vitro* transcriptomic study points to the same genes/pathways/disease outcomes as predicted by *in vivo* studies and associated human data (Guruge *et al.,* 2006; Jin *et al.,* 2020). Our results support an effect of PFAS on lipogenesis in human liver cells that warrants further research.

Benchmark concentration analysis was conducted to compare potencies of the four PFAS. Based on median gene benchmark concentration as a central estimate of the transcriptional activity, PFOS initially appeared to be the most potent; however, at the later timepoints, PFDS and PFOA had similar potencies to PFOS. The same pattern is observed in the gene accumulation plots, which show the initial PFOS curve to be left shifted relative to the other PFAS, but with PFDS and PFOA curves more super- imposed over PFOS at the later time points. It is important to bear in mind that our potency analysis is based on nominal concentrations of the PFAS exposures with these *in vitro* models. Cell permeability and uptake of PFOS has been found to be much faster than PFBS in human lung epithelial and murine preadipocyte embryonic fibroblasts (Sanchez Garcia *et al*., 2018), and PFOS uptake efficiency was 20 times higher than PFBS in an *in vitro* HEK293 cell model (Zhao *et al*., 2017). Furthermore, both active and passive transport mechanisms for PFOA have been identified in isolated rat hepatocytes (Han *et al*., 2008) and in the human colorectal adenocarcinoma cell line Caco-2 (Kimura *et al*., 2017). In humans, PFOS accumulates within the liver to a greater extent than PFBS, PFOA, and PFDS (Pérez *et al*., 2013). Based on the premise of selective uptake and concentrations of different PFAS, it is reasonable to speculate that differences in PFAS potencies over time may be influenced in part by the kinetics of cellular uptake and cell permeability. Zhao et al. (2015; 2017) found that several PFAS are transported into hepatocytes via Na+ / taurocholate co-transporting polypeptide (NTCP) and that human organic anion transporting polypeptides (OATPs), which are expressed in the spheroids in our study (data not shown), contributed to the enterohepatic circulation of certain PFAS and extend their half-lives. Thus, one important limitation of our study was that we did not measure the concentration of PFAS within the cells in the spheroids or the free fraction of the chemicals. Future studies will examine internal concentrations of PFAS within the spheroid cells over time.

Pharmacokinetic and exposure information were only available for PFOS and PFOA to derive bioactivity exposure ratios. The two approaches used (Approach 1: 5^th^percentile gene benchmark concentration and Approach 2: lowest pathway benchmark concentration) yielded results that were within 10-fold of the apical endpoint bioactivity exposure ratio, lending confidence to these approaches. Several of the lowest pathway benchmark concentrations across the PFAS are pathophysiologically relevant, including mevalonate pathway I, fatty acid activation, and stearate biosynthesis. These pathways are also consistent with the literature indicating that both PFOS (Reif *et al*., 2010) and PFOA (Das *et al*., 2017) cause hepatic steatosis (i.e., fatty liver disease), providing biological plausibility that these transcriptomic points of departure are meaningful. Steatosis was highlighted as an important endpoint within the US EPA’s Integrated Risk Information System (IRIS) on PFOS/PFOA and is recognized as a key step in liver disease progression (Kaiser *et al*., 2012). Wetmore *et al*. (2015b) proposed that a bioactivity exposure ratio cut-off > 100 suggests that the administered equivalent dose of biological activity is significantly different from daily population exposure estimates. Our study yielded bioactivity exposure ratios that were close to or less than <100, indicating that there is a relatively narrow margin between highly exposed members of the US population and activity *in vitro* for PFOS and PFOA. We note that the assumptions used within our approach to derive a value for the bioactivity exposure ratio from the administered equivalent dose do not account for the bioaccumulation from repeat low-dose exposures, which may be a limitation; this was a consideration when interpreting the values of the bioactivity exposure ratio within the investigation.

Liver concentration values were estimated from median values in Perez *et al*., 2013, and were equivalent to 0.002, 0.08, and 0.01 µM for PFBS, PFOS, and PFOA, respectively; PFDS could not be compared as observations were below detection limits within this study (Perez *et a*l., 2013). Since serum concentration measured from the same area were 10 to 5 times lower for the population (Ericson *et al*., 2007), the liver concentration can be used as an adequate marker of exposure against the benchmark concentrations values. The lowest median gene the benchmark concentrations within our observations were 46.3, 8.2, and 6.4 µM for PFBS, PFOS, and PFOA respectively. The PFBS estimates from the literature are several orders of magnitude different from the lowest median gene benchmark concentration in our findings but there was a < 1000-fold difference for PFOA, and approximately 100-fold difference in ratio for PFOS. Considering a more conservative estimate of the margin of potential exposure, the median benchmark concentration of the gene-set from the lowest, most sensitive pathway, yields ratios of 0.74 and 1.2 for PFOS and PFOA respectively, suggesting that changes in gene expression at current exposure levels within this population are likely.

High-throughput transcriptomics is of considerable value because it broadly informs global biological perturbations induced by chemical exposures and can be readily used to compare similarities/differences in biological activities and potencies across chemicals. This study produced a baseline transcriptomic data set and pipeline to be applied in future studies to study data-poor PFAS. We demonstrate several approaches that can be used to inform hazard, mode of action, and potency.

## Acknowledgements

We thank Dr. Francesco Marchetti, Dr. Marc Beal, and Dr. Scott Auerbach for their helpful commentary and review of this research, the US EPA for their valuable consultations, as well as BioSpyder for their contributions on this project. We would like to thank Dr. Iain Lambert, Carleton University, for logistical support for this project.

## Funding

We acknowledge the Water and Air Quality Bureau, HECSB, Health Canada, the Chemicals and Environmental Health Management Bureau, HECSB, Health Canada, and the Environmental Health Sciences & Research Bureau, HECSB, Health Canada who jointly funded this research.

## Notes

https://www.ncbi.nlm.nih.gov/geo/query/acc.cgi?acc=GSE144775

### Competing Interest Statement

The authors have declared no competing interest.

