## Supplementary material for "High-throughput transcriptomic analysis of human primary hepatocyte spheroids exposed to per- and polyfluoroalkyl substances (PFAS) as a platform for relative potency characterization": Supplemental Files-March 10.docx

**Please note Supplemental Files 1-5 are also available as pdfs for increased resolution.**

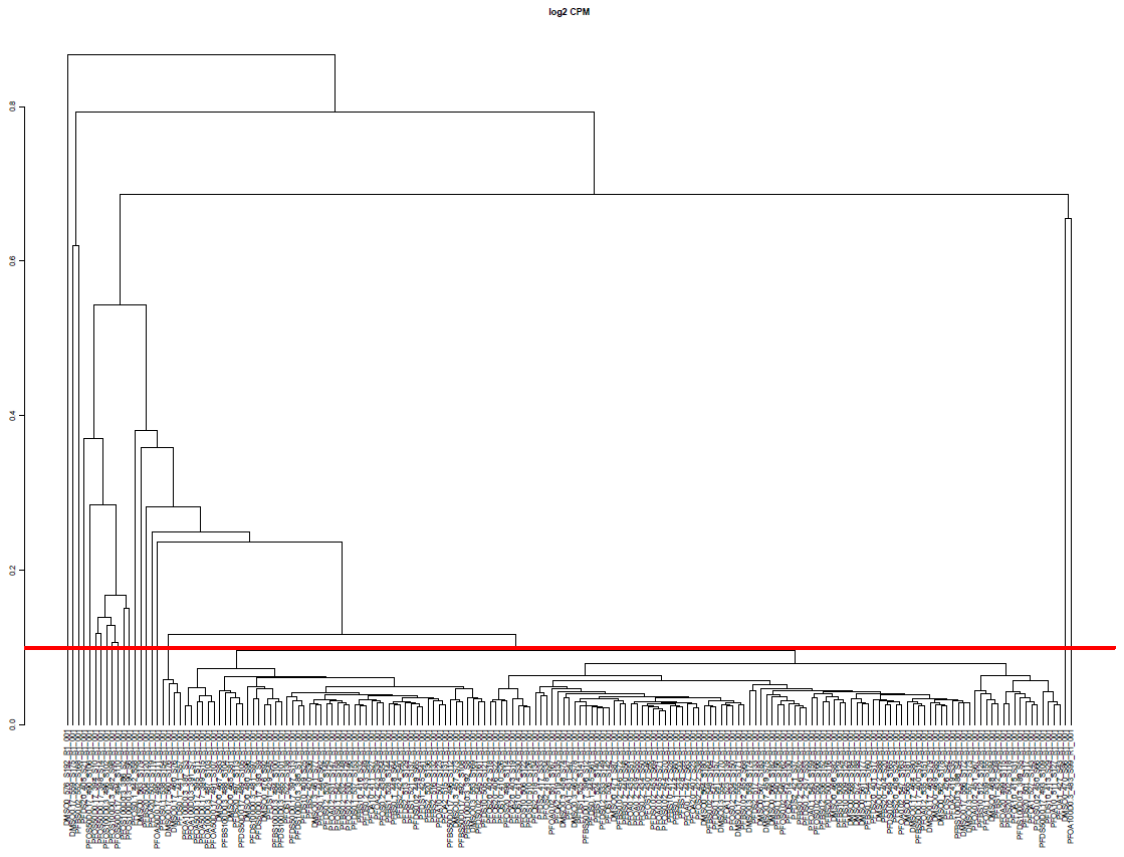

Supplemental File 1-1: Dendrogram of day-1 samples with outliers. Dendrograms were generated in R for each timepoint with and without outlier samples. Dendrograms were based on the one minus spearman correlation using complete linkage. A red line is drawn at the 0.1 dissimilarity to identify samples that clustered as singletons. These samples were removed in the final analyses.

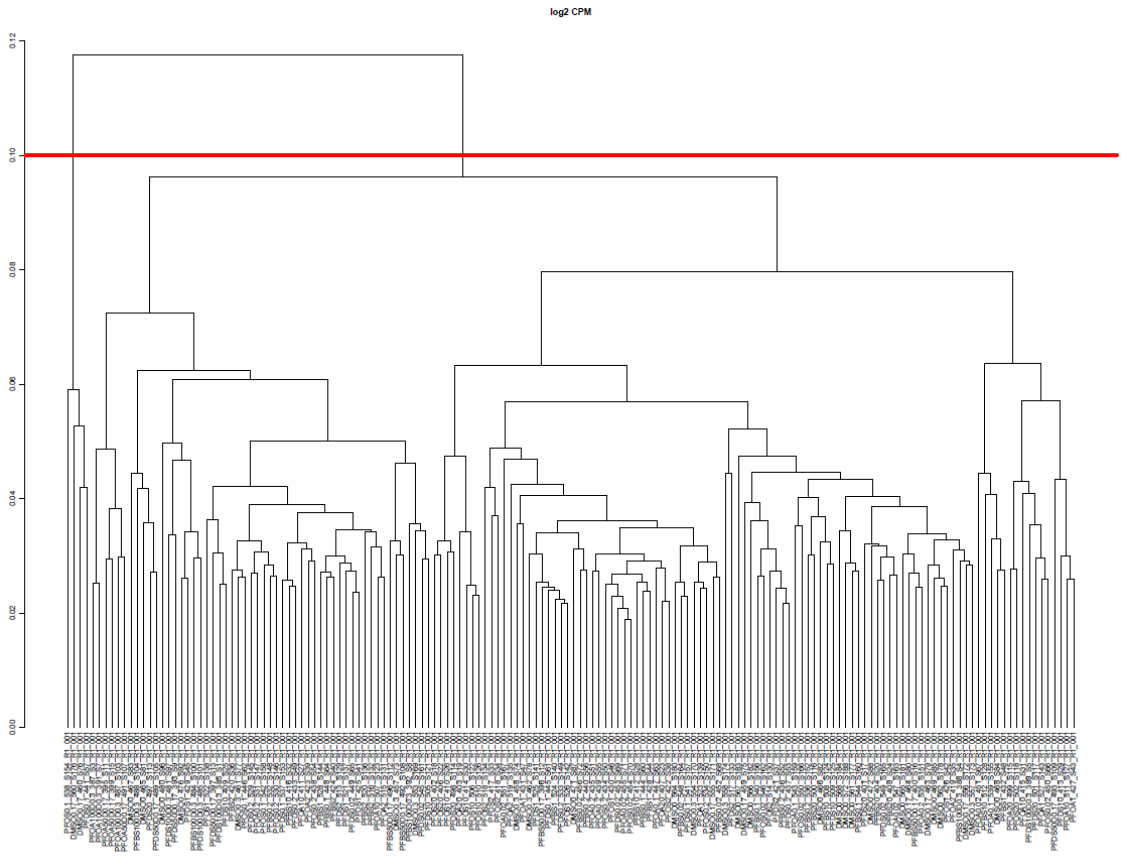

Supplemental File 1-2: Dendrogram of day-1 samples without outliers. Dendrograms were generated in R for each timepoint with and without outlier samples. Dendrograms were based on the one minus spearman correlation using complete linkage. A red line is drawn at the 0.1 dissimilarity to identify samples that clustered as singletons. These samples were removed in the final analyses.

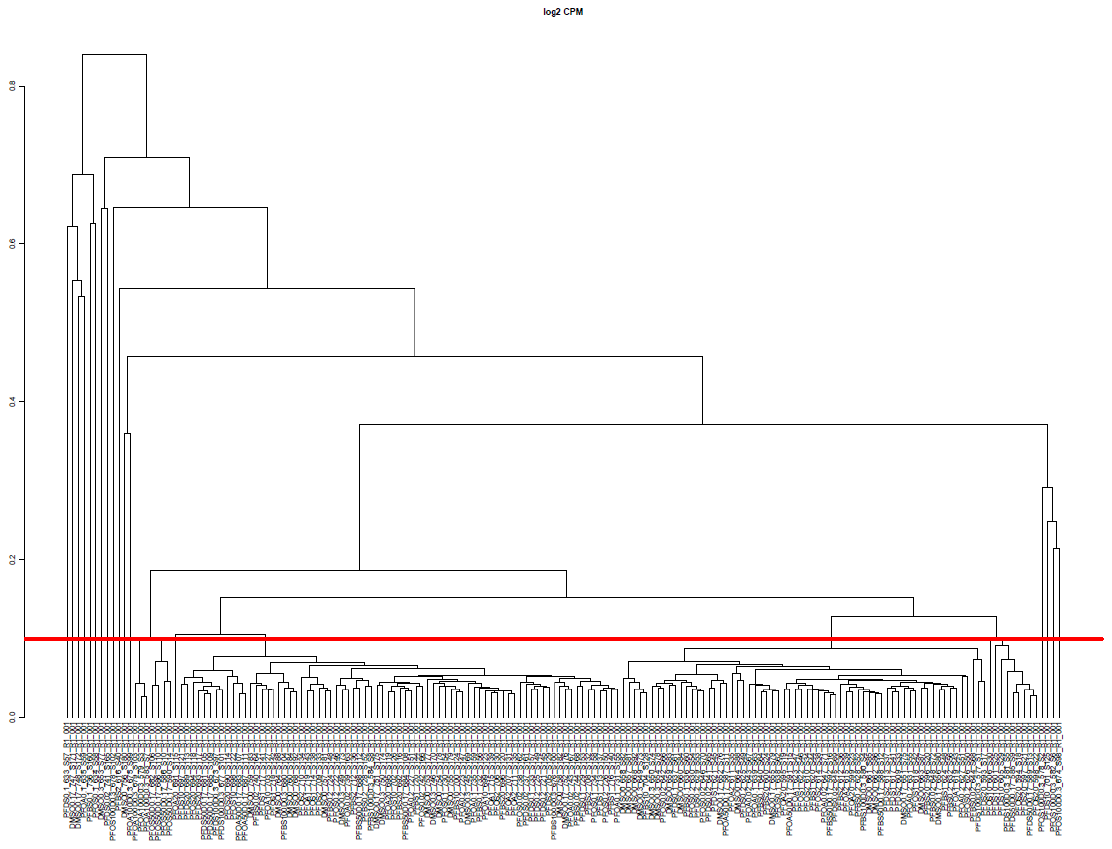

Supplemental File 1-3: Dendrogram of day-4 samples with outliers. Dendrograms were generated in R for each timepoint with and without outlier samples. Dendrograms were based on the one minus spearman correlation using complete linkage. A red line is drawn at the 0.1 dissimilarity to identify samples that clustered as singletons. These samples were removed in the final analyses.

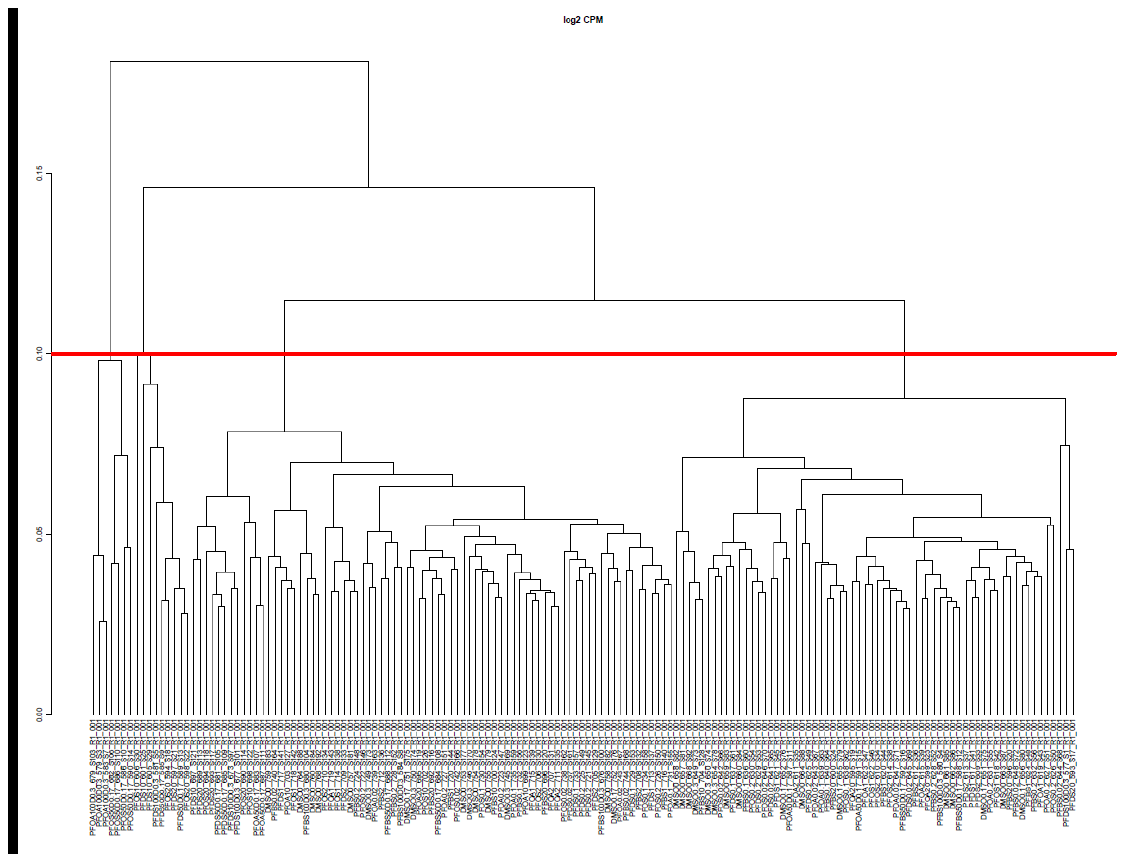

Supplemental File 1-4: Dendrogram of day-4 samples without outliers. Dendrograms were generated in R for each timepoint with and without outlier samples. Dendrograms were based on the one minus spearman correlation using complete linkage. A red line is drawn at the 0.1 dissimilarity to identify samples that clustered as singletons. These samples were removed in the final analyses.

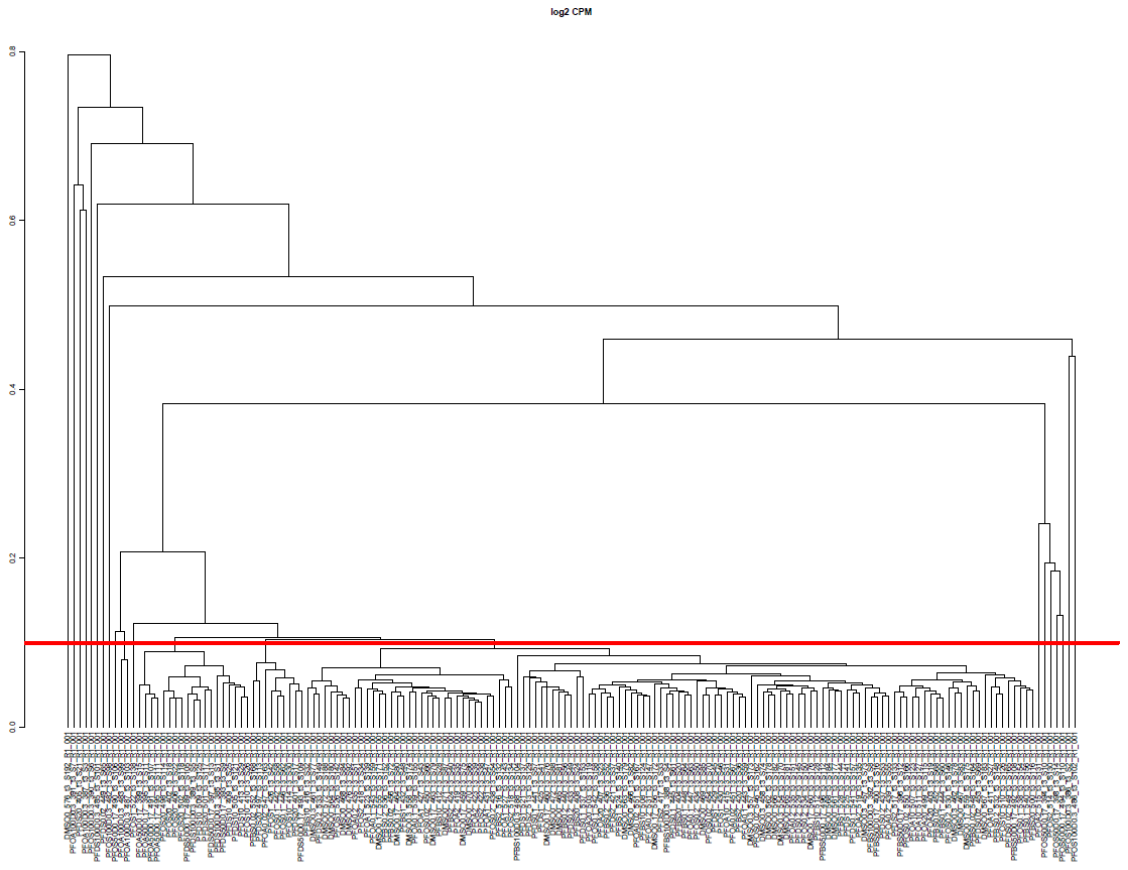

Supplemental File 1-5: Dendrogram of day-10 samples with outliers. Dendrograms were generated in R for each timepoint with and without outlier samples. Dendrograms were based on the one minus spearman correlation using complete linkage. A red line is drawn at the 0.1 dissimilarity to identify samples that clustered as singletons. These samples were removed in the final analyses.

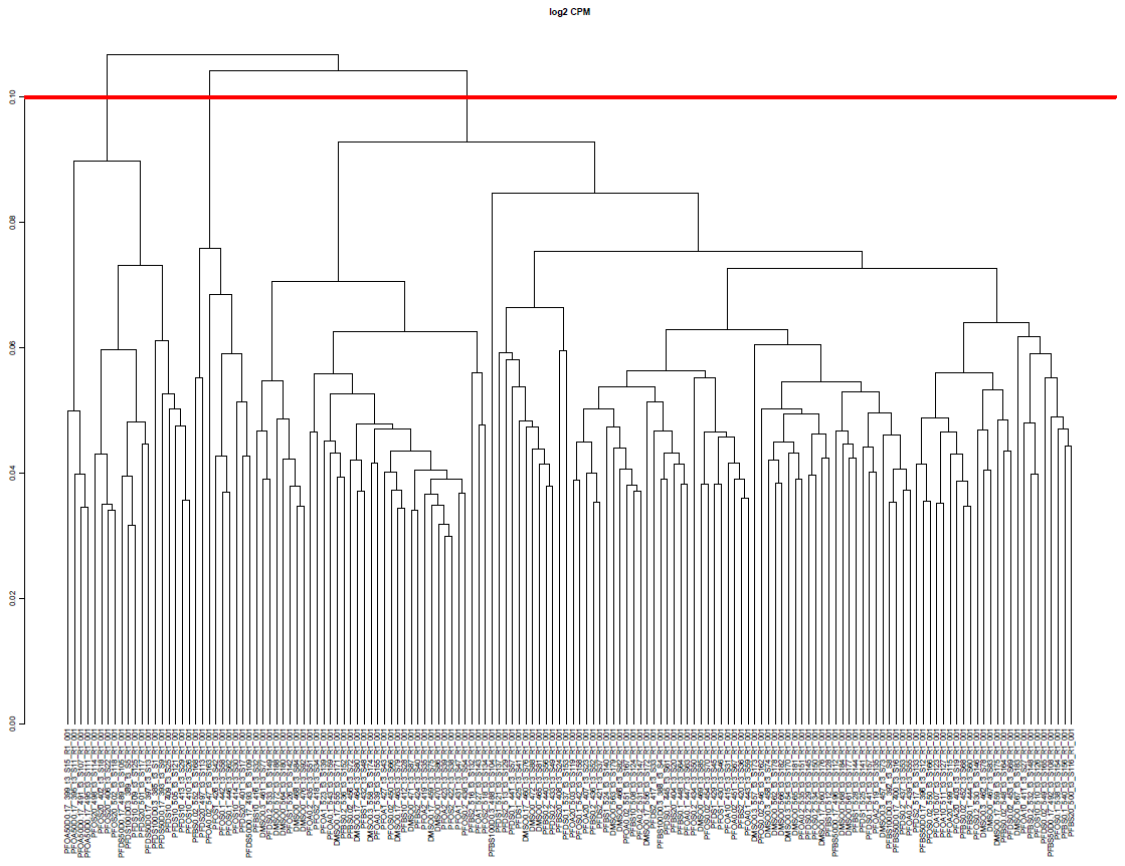

Supplemental File 1-6: Dendrogram of day-10 samples without outliers. Dendrograms were generated in R for each timepoint with and without outlier samples. Dendrograms were based on the one minus spearman correlation using complete linkage. A red line is drawn at the 0.1 dissimilarity to identify samples that clustered as singletons. These samples were removed in the final analyses.

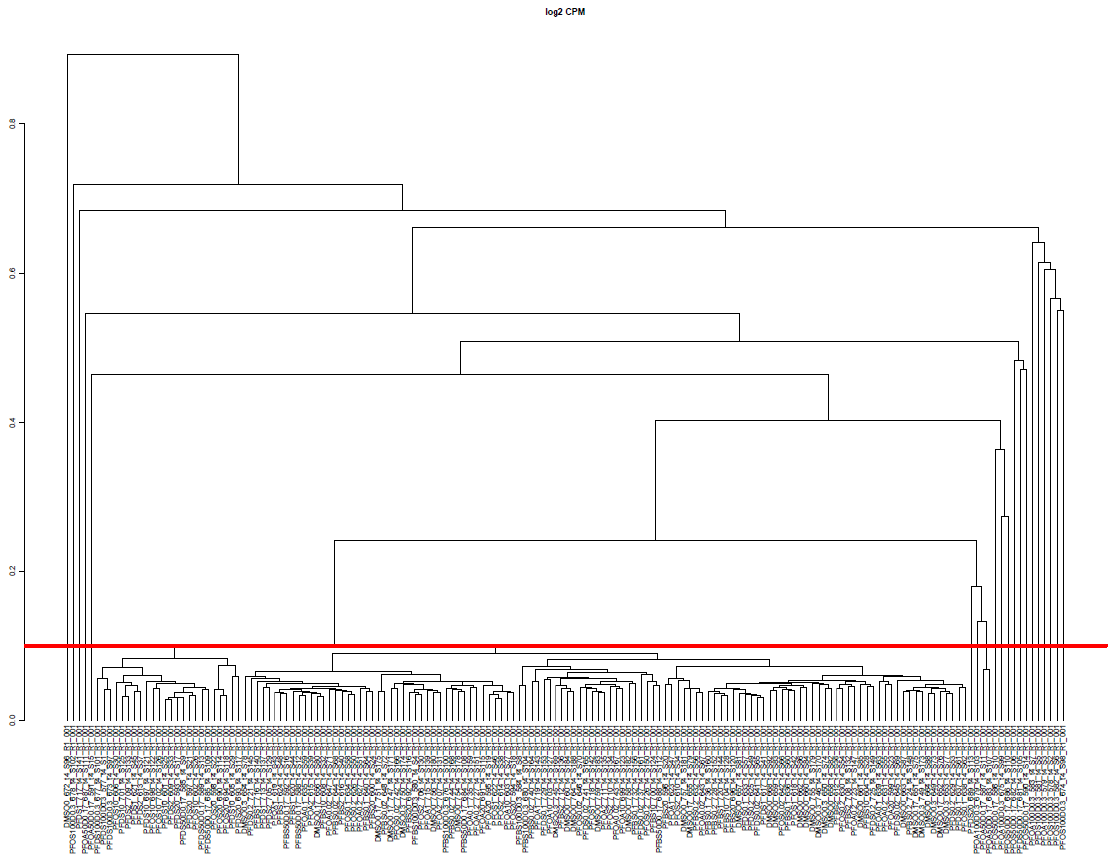

Supplemental File 1-7: Dendrogram of day-14 samples with outliers. Dendrograms were generated in R for each timepoint with and without outlier samples. Dendrograms were based on the one minus spearman correlation using complete linkage. A red line is drawn at the 0.1 dissimilarity to identify samples that clustered as singletons. These samples were removed in the final analyses.

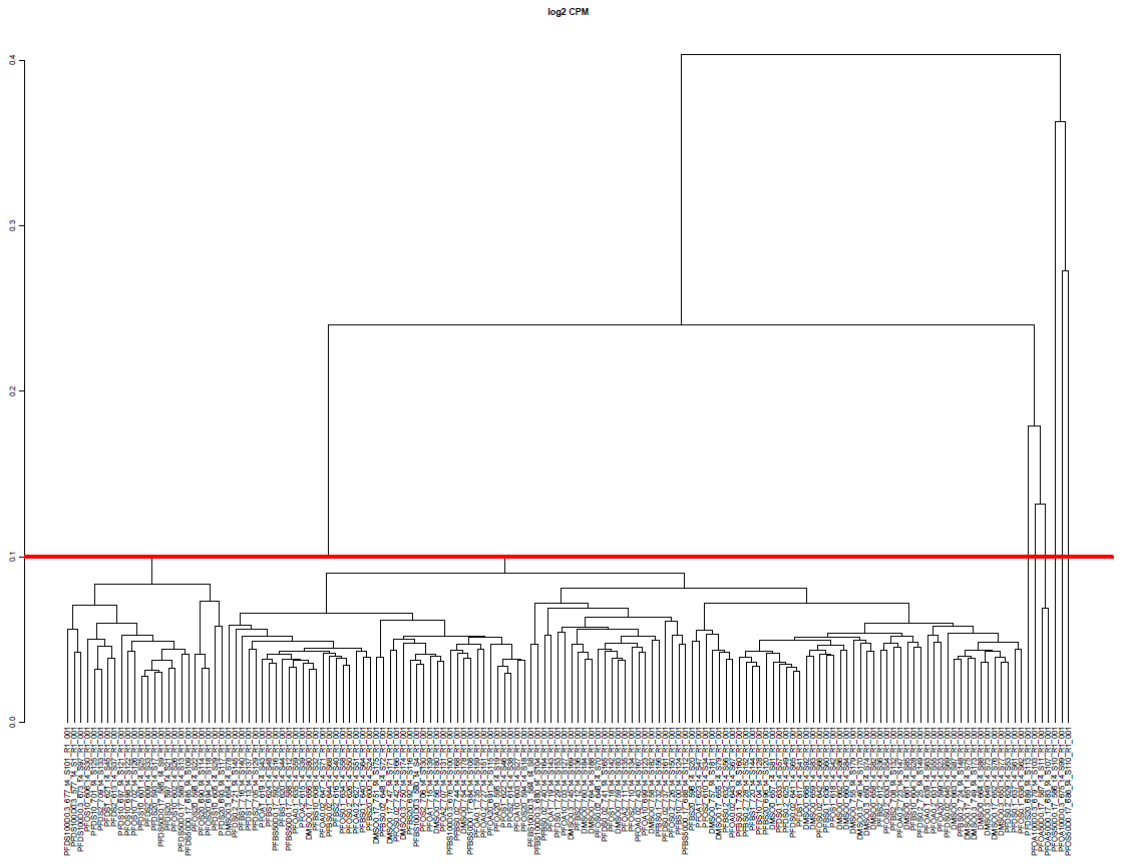
Supplemental File 1-8: Dendrogram of day-14 samples without outliers. Dendrograms were generated in R for each timepoint with and without outlier samples. Dendrograms were based on the one minus spearman correlation using complete linkage. A red line is drawn at the 0.1 dissimilarity to identify samples that clustered as singletons. These samples were removed in the final analyses.

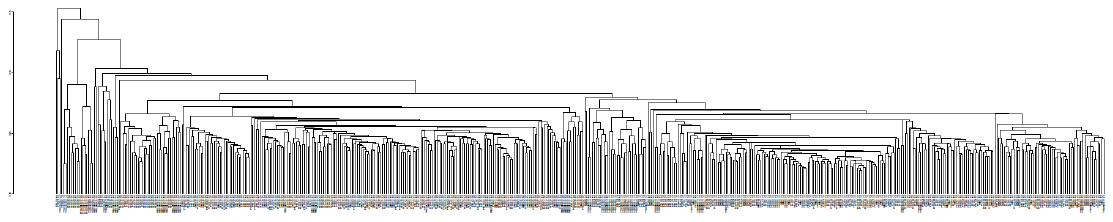

Supplemental File 1-9: Dendrogram of all samples with outliers removed. Dendrograms were generated in R using the one minus Spearman correlation metric with complete linkage

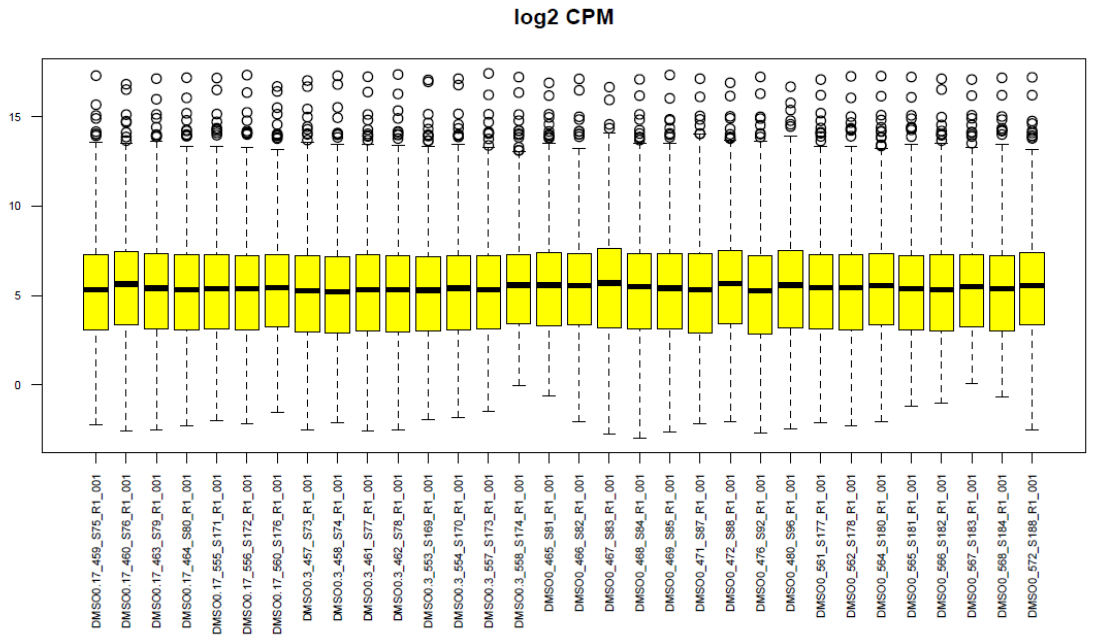
Supplemental File 2-1: DMSO boxplots of mapped reads day-1. Boxplots were generated in the R statistical environment for each sample using the filtered counts per million. Sample distributions of probes that had at least a median of 5 counts in at least one treatment group are displayed in each boxplot.

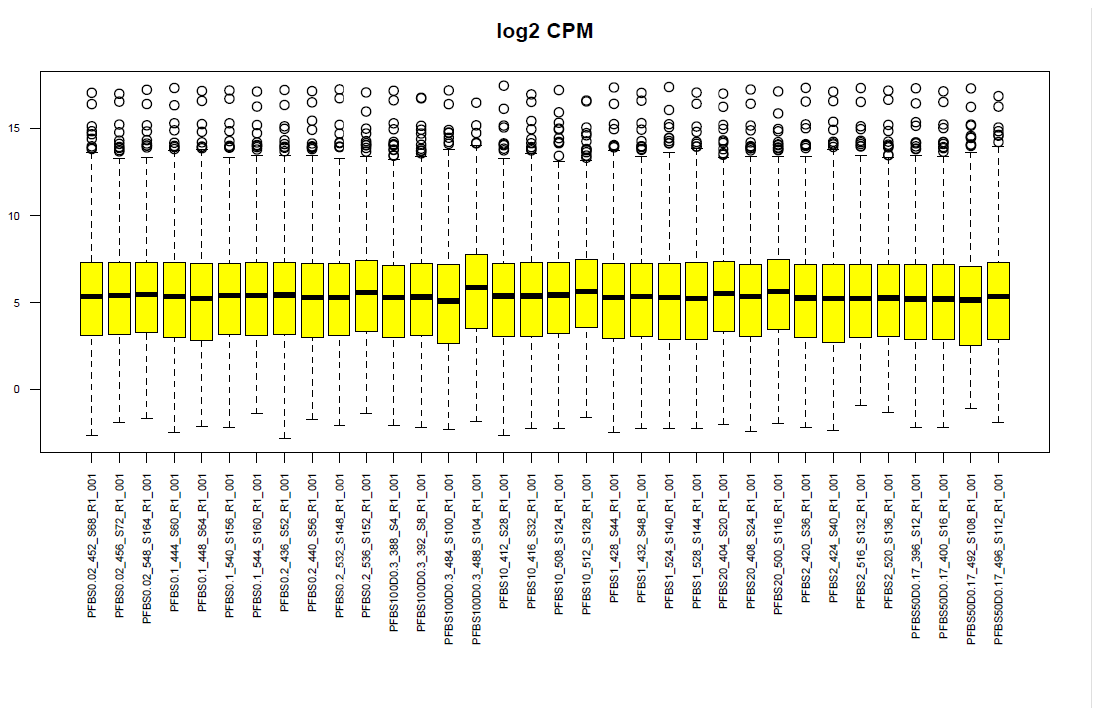
Supplemental File 2-2: PFBS boxplots of mapped reads day-1.Boxplots were generated in the R statistical environment for each sample using the filtered counts per million. Sample distributions of probes that had at least a median of 5 counts in at least one treatment group are displayed in each boxplot.

Supplemental figure XX:Upstream regulator predictions from the differentially expressed genes for the top non-cytotoxic concentration of each PFAS. The z-score filters were relaxed to 2.0 with a Benjamini Hochberg adj p-value of 0.05 for this analysis. Blue represents inhibited upstream regulators, and orange represents activated regulators.

PFOA 20µM DAY 1

PFOA 20µM DAY 4

PFOA 20µM DAY 10

PFOA 20µM DAY 14

PFBS 100µM DAY 1

PFBS 100µM DAY 4

PFBS 100µM DAY 10

PFBS 100µM DAY 14

PFOS 20µM DAY 1

PFOS 20µM DAY 4

PFOS 20µM DAY 10

PFOS 20µM DAY 14

PFDS 100µM DAY 1

PFDS 100µM DAY 4

PFDS 100µM DAY 10

PFDS 100µM DAY 14

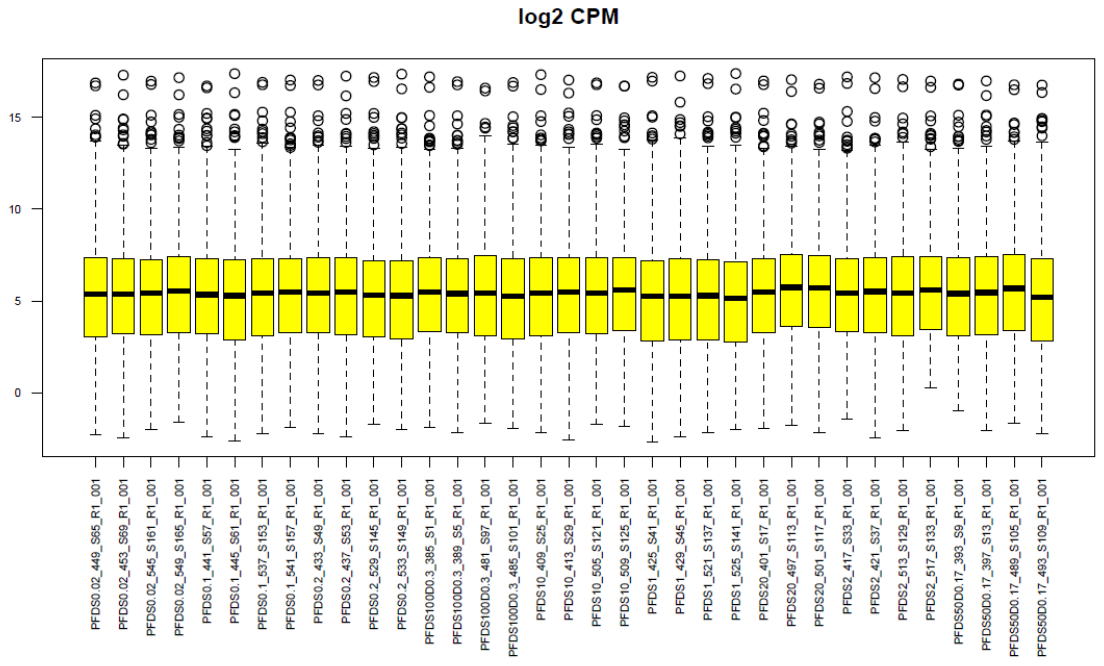

Supplemental File 2-3: PFDS boxplots of mapped reads day-1. Boxplots were generated in the R statistical environment for each sample using the filtered counts per million. Sample distributions of probes that had at least a median of 5 counts in at least one treatment group are displayed in each boxplot.

Supplemental figure XX:Upstream regulator predictions from the differentially expressed genes for the top non-cytotoxic concentration of each PFAS. The z-score filters were relaxed to 2.0 with a Benjamini Hochberg adj p-value of 0.05 for this analysis. Blue represents inhibited upstream regulators, and orange represents activated regulators.

PFOA 20µM DAY 1

PFOA 20µM DAY 4

PFOA 20µM DAY 10

PFOA 20µM DAY 14

PFBS 100µM DAY 1

PFBS 100µM DAY 4

PFBS 100µM DAY 10

PFBS 100µM DAY 14

PFOS 20µM DAY 1

PFOS 20µM DAY 4

PFOS 20µM DAY 10

PFOS 20µM DAY 14

PFDS 100µM DAY 1

PFDS 100µM DAY 4

PFDS 100µM DAY 10

PFDS 100µM DAY 14

Supplemental figure XX:Upstream regulator predictions from the differentially expressed genes for the top non-cytotoxic concentration of each PFAS. The z-score filters were relaxed to 2.0 with a Benjamini Hochberg adj p-value of 0.05 for this analysis. Blue represents inhibited upstream regulators, and orange represents activated regulators.

PFOA 20µM DAY 1

PFOA 20µM DAY 4

PFOA 20µM DAY 10

PFOA 20µM DAY 14

PFBS 100µM DAY 1

PFBS 100µM DAY 4

PFBS 100µM DAY 10

PFBS 100µM DAY 14

PFOS 20µM DAY 1

PFOS 20µM DAY 4

PFOS 20µM DAY 10

PFOS 20µM DAY 14

PFDS 100µM DAY 1

PFDS 100µM DAY 4

PFDS 100µM DAY 10

PFDS 100µM DAY 14

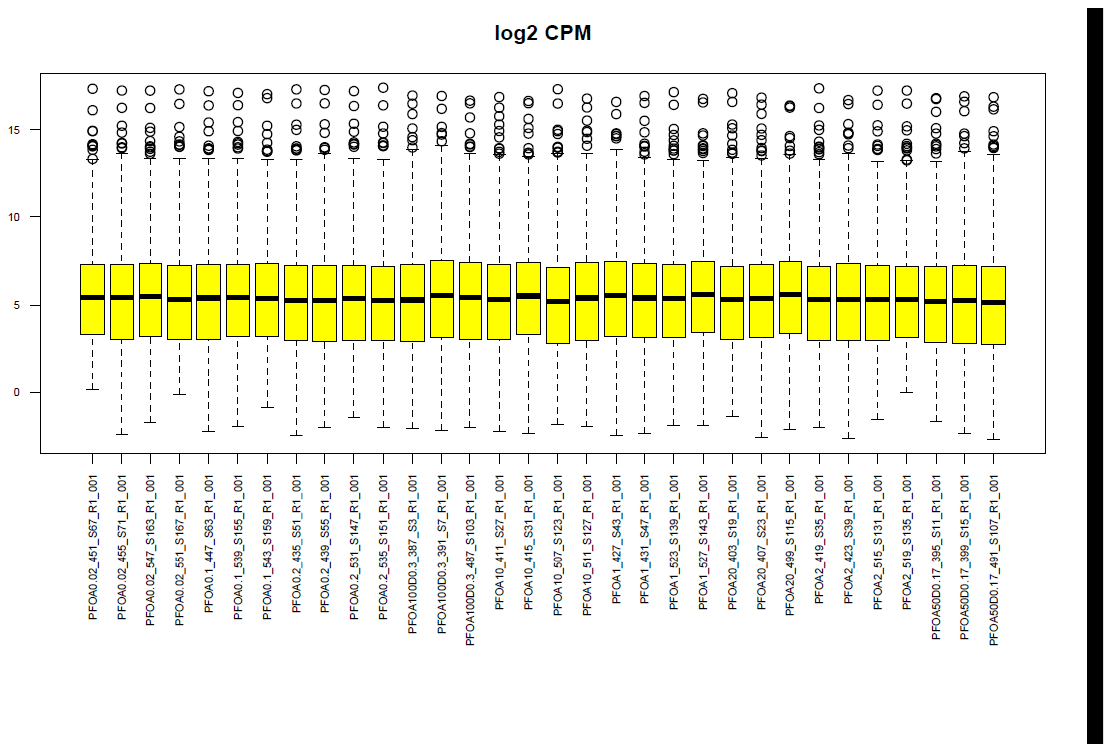

Supplemental File 2-4: PFOA boxplots of mapped reads day-1. Boxplots were generated in the R statistical environment for each sample using the filtered counts per million. Sample distributions of probes that had at least a median of 5 counts in at least one treatment group are displayed in each boxplot.

Supplemental figure XX:Upstream regulator predictions from the differentially expressed genes for the top non-cytotoxic concentration of each PFAS. The z-score filters were relaxed to 2.0 with a Benjamini Hochberg adj p-value of 0.05 for this analysis. Blue represents inhibited upstream regulators, and orange represents activated regulators.

PFOA 20µM DAY 1

PFOA 20µM DAY 4

PFOA 20µM DAY 10

PFOA 20µM DAY 14

PFBS 100µM DAY 1

PFBS 100µM DAY 4

PFBS 100µM DAY 10

PFBS 100µM DAY 14

PFOS 20µM DAY 1

PFOS 20µM DAY 4

PFOS 20µM DAY 10

PFOS 20µM DAY 14

PFDS 100µM DAY 1

PFDS 100µM DAY 4

PFDS 100µM DAY 10

PFDS 100µM DAY 14

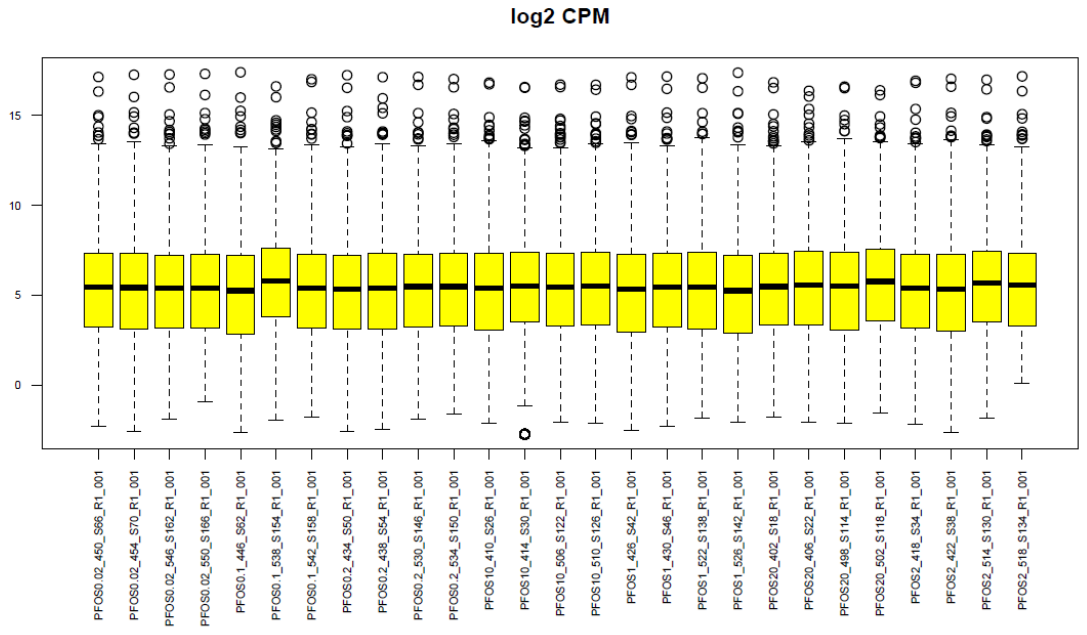
Supplemental File 2-5: PFOS boxplots of mapped reads day-1. Boxplots were generated in the R statistical environment for each sample using the filtered counts per million. Sample distributions of probes that had at least a median of 5 counts in at least one treatment group are displayed in each boxplot.

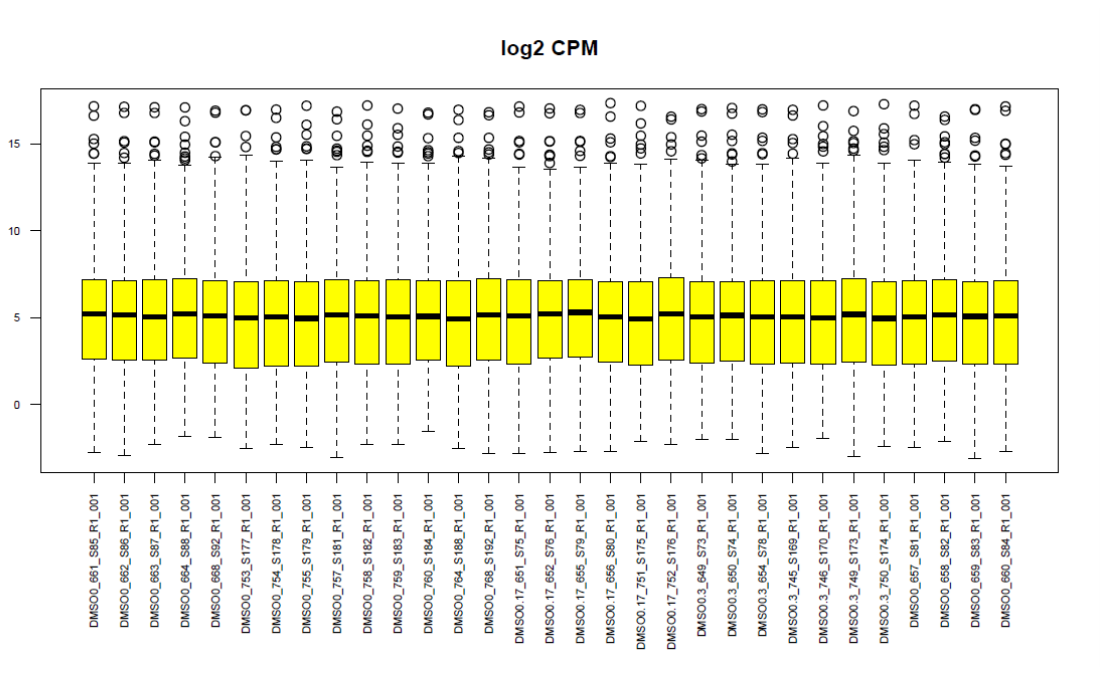
Supplemental File 3-1: DMSO boxplots of mapped reads day-4. Boxplots were generated in the R statistical environment for each sample using the filtered counts per million. Sample distributions of probes that had at least a median of 5 counts in at least one treatment group are displayed in each boxplot.

Supplemental figure XX:Upstream regulator predictions from the differentially expressed genes for the top non-cytotoxic concentration of each PFAS. The z-score filters were relaxed to 2.0 with a Benjamini Hochberg adj p-value of 0.05 for this analysis. Blue represents inhibited upstream regulators, and orange represents activated regulators.

PFOA 20µM DAY 1

PFOA 20µM DAY 4

PFOA 20µM DAY 10

PFOA 20µM DAY 14

PFBS 100µM DAY 1

PFBS 100µM DAY 4

PFBS 100µM DAY 10

PFBS 100µM DAY 14

PFOS 20µM DAY 1

PFOS 20µM DAY 4

PFOS 20µM DAY 10

PFOS 20µM DAY 14

PFDS 100µM DAY 1

PFDS 100µM DAY 4

PFDS 100µM DAY 10

PFDS 100µM DAY 14

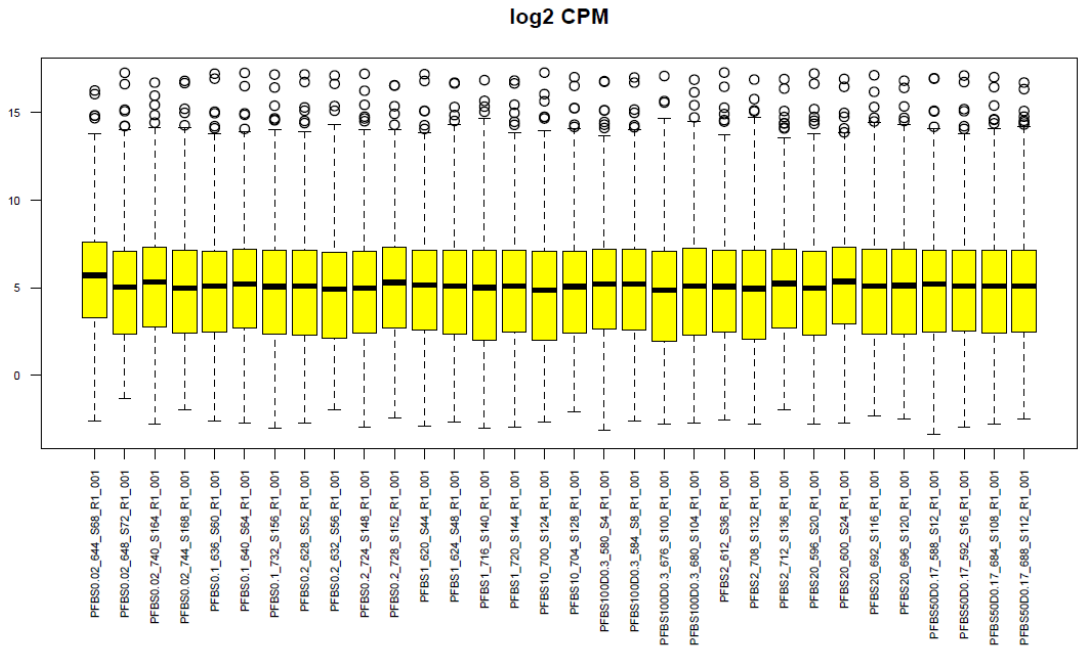

Supplemental File 3-2: PFBS boxplots of mapped reads day-4. Boxplots were generated in the R statistical environment for each sample using the filtered counts per million. Sample distributions of probes that had at least a median of 5 counts in at least one treatment group are displayed in each boxplot.

Supplemental figure XX:Upstream regulator predictions from the differentially expressed genes for the top non-cytotoxic concentration of each PFAS. The z-score filters were relaxed to 2.0 with a Benjamini Hochberg adj p-value of 0.05 for this analysis. Blue represents inhibited upstream regulators, and orange represents activated regulators.

PFOA 20µM DAY 1

PFOA 20µM DAY 4

PFOA 20µM DAY 10

PFOA 20µM DAY 14

PFBS 100µM DAY 1

PFBS 100µM DAY 4

PFBS 100µM DAY 10

PFBS 100µM DAY 14

PFOS 20µM DAY 1

PFOS 20µM DAY 4

PFOS 20µM DAY 10

PFOS 20µM DAY 14

PFDS 100µM DAY 1

PFDS 100µM DAY 4

PFDS 100µM DAY 10

PFDS 100µM DAY 14

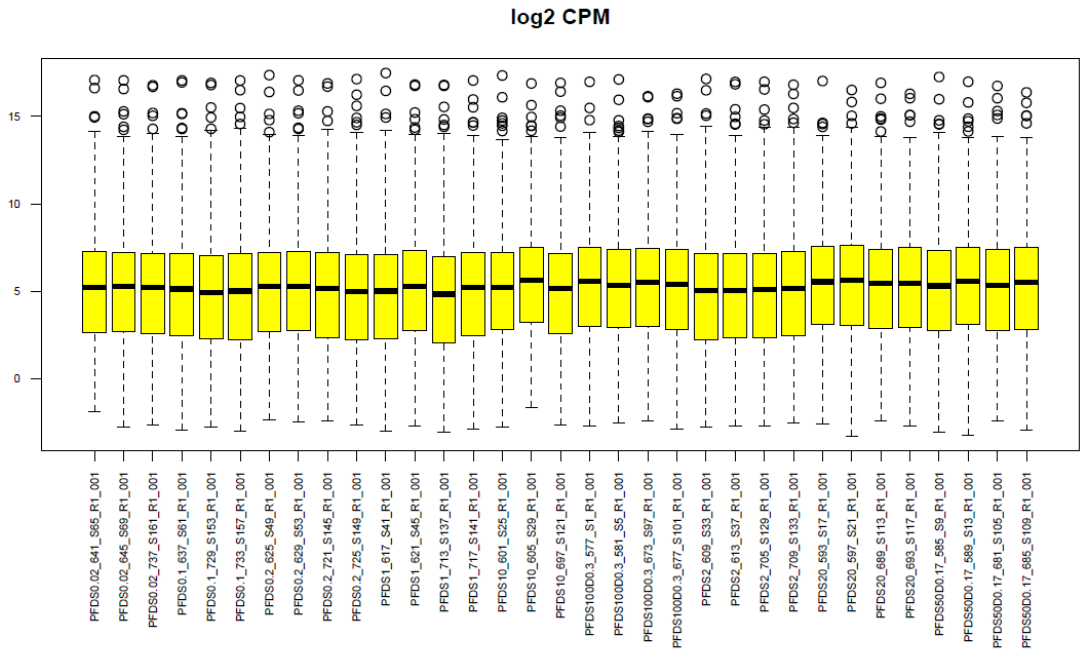

Supplemental File 3-3: PFDS boxplots of mapped reads day-4. Boxplots were generated in the R statistical environment for each sample using the filtered counts per million. Sample distributions of probes that had at least a median of 5 counts in at least one treatment group are displayed in each boxplot.

Supplemental figure XX:Upstream regulator predictions from the differentially expressed genes for the top non-cytotoxic concentration of each PFAS. The z-score filters were relaxed to 2.0 with a Benjamini Hochberg adj p-value of 0.05 for this analysis. Blue represents inhibited upstream regulators, and orange represents activated regulators.

PFOA 20µM DAY 1

PFOA 20µM DAY 4

PFOA 20µM DAY 10

PFOA 20µM DAY 14

PFBS 100µM DAY 1

PFBS 100µM DAY 4

PFBS 100µM DAY 10

PFBS 100µM DAY 14

PFOS 20µM DAY 1

PFOS 20µM DAY 4

PFOS 20µM DAY 10

PFOS 20µM DAY 14

PFDS 100µM DAY 1

PFDS 100µM DAY 4

PFDS 100µM DAY 10

PFDS 100µM DAY 14

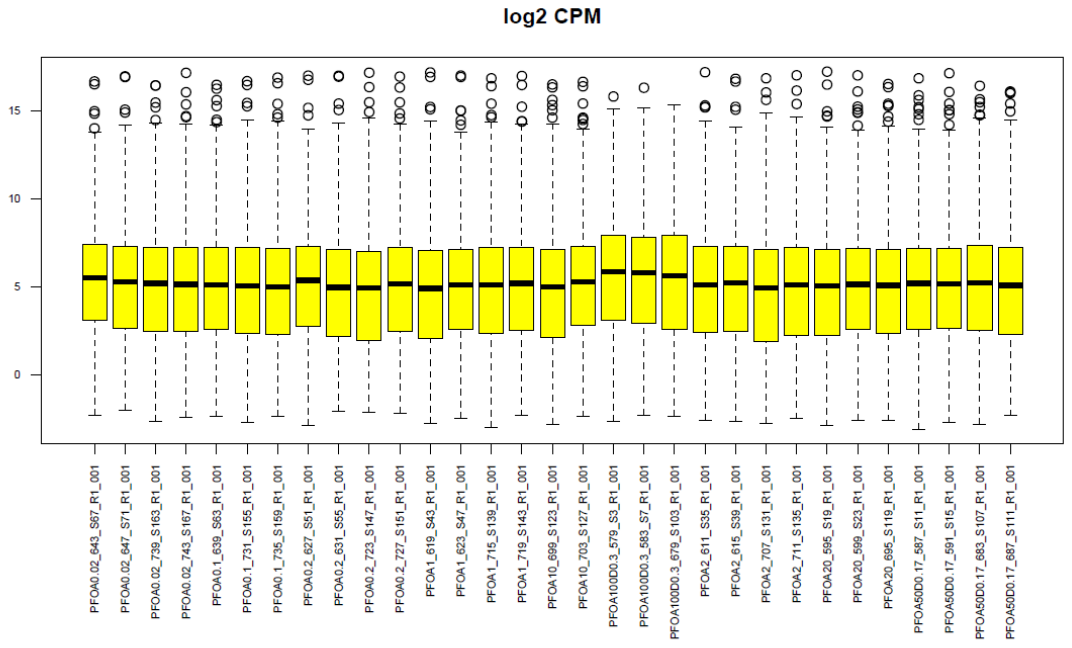

Supplemental File 3-4: PFOA boxplots of mapped reads day-4. Boxplots were generated in the R statistical environment for each sample using the filtered counts per million. Sample distributions of probes that had at least a median of 5 counts in at least one treatment group are displayed in each boxplot.

Supplemental figure XX:Upstream regulator predictions from the differentially expressed genes for the top non-cytotoxic concentration of each PFAS. The z-score filters were relaxed to 2.0 with a Benjamini Hochberg adj p-value of 0.05 for this analysis. Blue represents inhibited upstream regulators, and orange represents activated regulators.

PFOA 20µM DAY 1

PFOA 20µM DAY 4

PFOA 20µM DAY 10

PFOA 20µM DAY 14

PFBS 100µM DAY 1

PFBS 100µM DAY 4

PFBS 100µM DAY 10

PFBS 100µM DAY 14

PFOS 20µM DAY 1

PFOS 20µM DAY 4

PFOS 20µM DAY 10

PFOS 20µM DAY 14

PFDS 100µM DAY 1

PFDS 100µM DAY 4

PFDS 100µM DAY 10

PFDS 100µM DAY 14

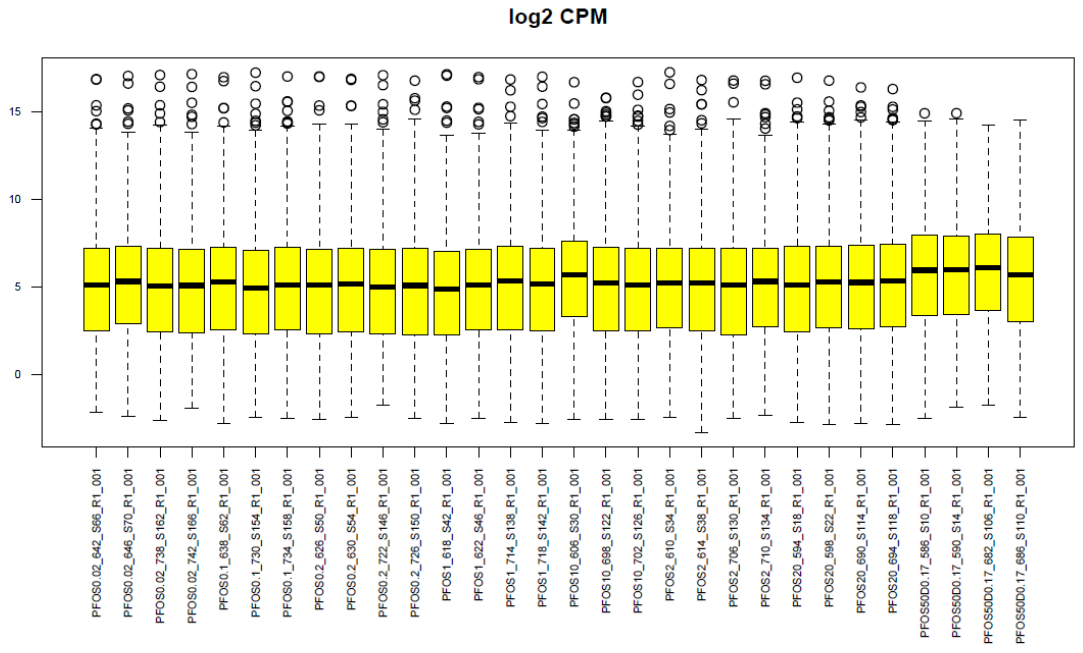

Supplemental File 3-5: PFOS boxplots of mapped reads day-4. Boxplots were generated in the R statistical environment for each sample using the filtered counts per million. Sample distributions of probes that had at least a median of 5 counts in at least one treatment group are displayed in each boxplot.

Supplemental figure XX:Upstream regulator predictions from the differentially expressed genes for the top non-cytotoxic concentration of each PFAS. The z-score filters were relaxed to 2.0 with a Benjamini Hochberg adj p-value of 0.05 for this analysis. Blue represents inhibited upstream regulators, and orange represents activated regulators.

PFOA 20µM DAY 1

PFOA 20µM DAY 4

PFOA 20µM DAY 10

PFOA 20µM DAY 14

PFBS 100µM DAY 1

PFBS 100µM DAY 4

PFBS 100µM DAY 10

PFBS 100µM DAY 14

PFOS 20µM DAY 1

PFOS 20µM DAY 4

PFOS 20µM DAY 10

PFOS 20µM DAY 14

PFDS 100µM DAY 1

PFDS 100µM DAY 4

PFDS 100µM DAY 10

PFDS 100µM DAY 14

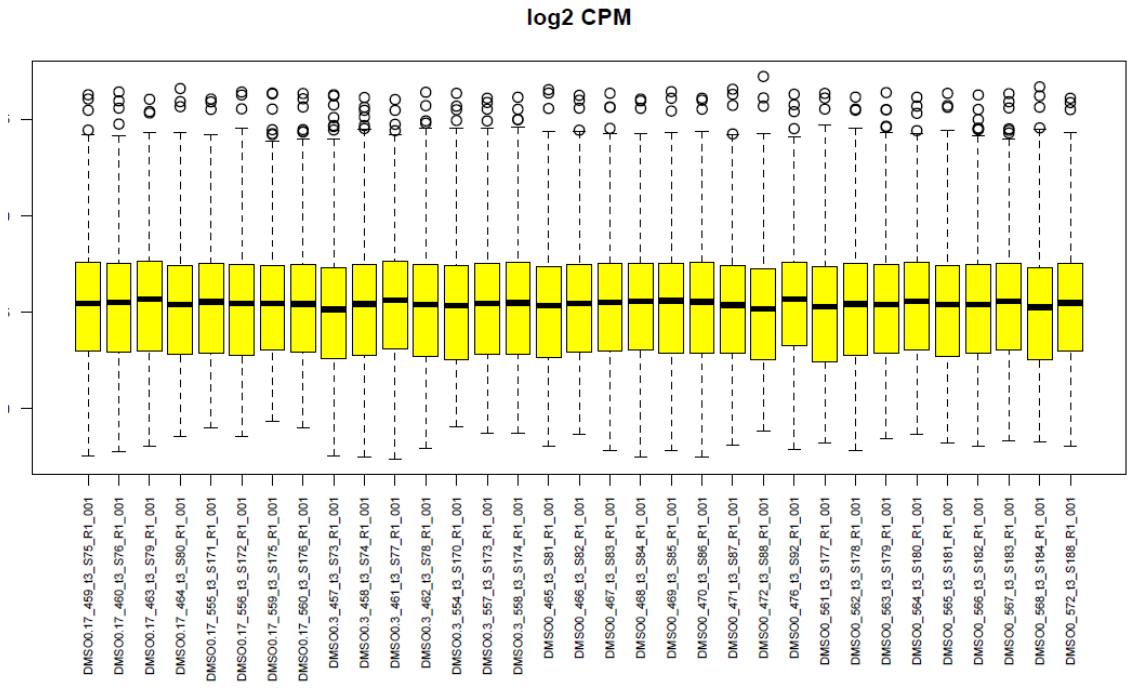

Supplemental File 4-1: DMSO boxplots of mapped reads day-10. Boxplots were generated in the R statistical environment for each sample using the filtered counts per million. Sample distributions of probes that had at least a median of 5 counts in at least one treatment group are displayed in each boxplot.

Supplemental figure XX:Upstream regulator predictions from the differentially expressed genes for the top non-cytotoxic concentration of each PFAS. The z-score filters were relaxed to 2.0 with a Benjamini Hochberg adj p-value of 0.05 for this analysis. Blue represents inhibited upstream regulators, and orange represents activated regulators.

PFOA 20µM DAY 1

PFOA 20µM DAY 4

PFOA 20µM DAY 10

PFOA 20µM DAY 14

PFBS 100µM DAY 1

PFBS 100µM DAY 4

PFBS 100µM DAY 10

PFBS 100µM DAY 14

PFOS 20µM DAY 1

PFOS 20µM DAY 4

PFOS 20µM DAY 10

PFOS 20µM DAY 14

PFDS 100µM DAY 1

PFDS 100µM DAY 4

PFDS 100µM DAY 10

PFDS 100µM DAY 14

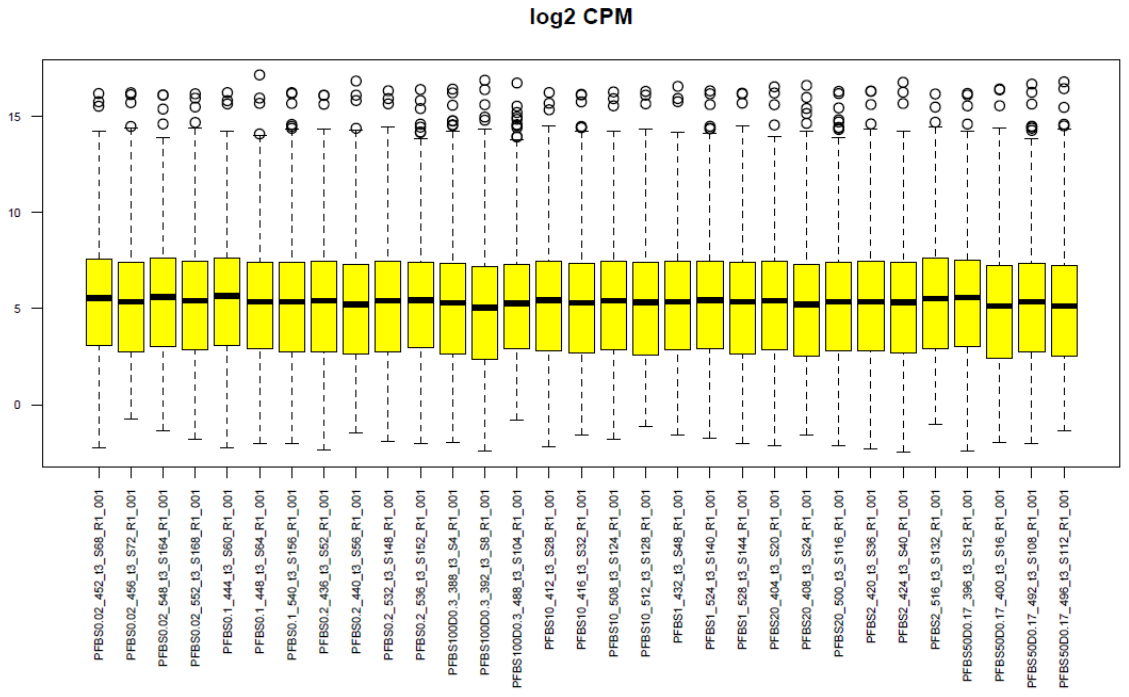
Supplemental File 4-2: PFBS boxplots of mapped reads day-10. Boxplots were generated in the R statistical environment for each sample using the filtered counts per million. Sample distributions of probes that had at least a median of 5 counts in at least one treatment group are displayed in each boxplot.

Supplemental figure XX:Upstream regulator predictions from the differentially expressed genes for the top non-cytotoxic concentration of each PFAS. The z-score filters were relaxed to 2.0 with a Benjamini Hochberg adj p-value of 0.05 for this analysis. Blue represents inhibited upstream regulators, and orange represents activated regulators.

PFOA 20µM DAY 1

PFOA 20µM DAY 4

PFOA 20µM DAY 10

PFOA 20µM DAY 14

PFBS 100µM DAY 1

PFBS 100µM DAY 4

PFBS 100µM DAY 10

PFBS 100µM DAY 14

PFOS 20µM DAY 1

PFOS 20µM DAY 4

PFOS 20µM DAY 10

PFOS 20µM DAY 14

PFDS 100µM DAY 1

PFDS 100µM DAY 4

PFDS 100µM DAY 10

PFDS 100µM DAY 14

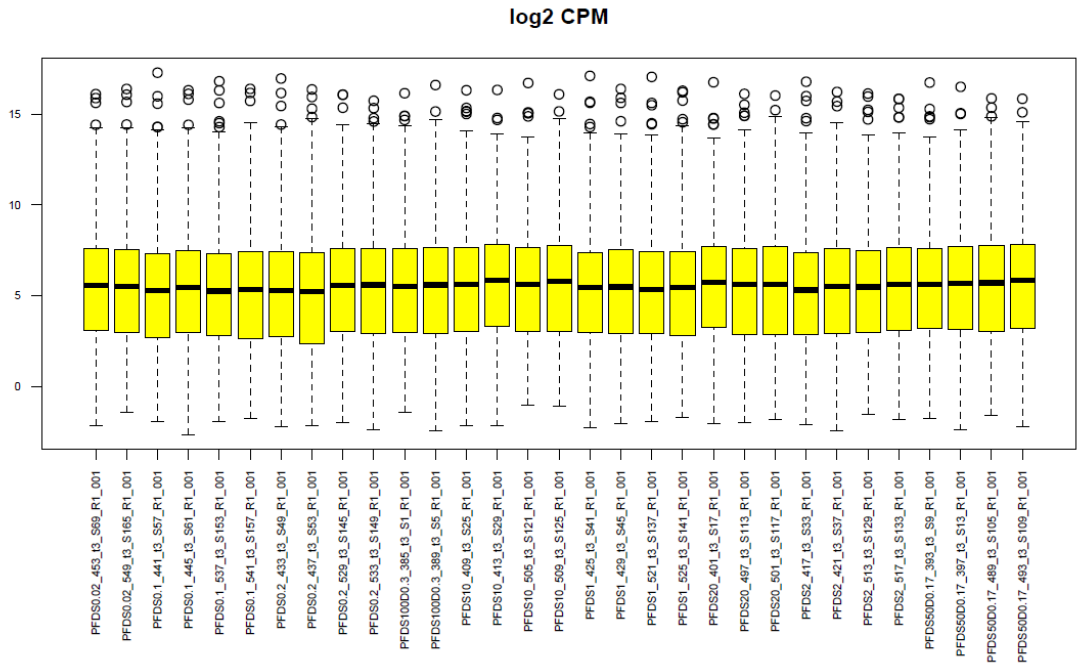

Supplemental File 4-3: PFDS boxplots day-10 of mapped reads. Boxplots were generated in the R statistical environment for each sample using the filtered counts per million. Sample distributions of probes that had at least a median of 5 counts in at least one treatment group are displayed in each boxplot.

Supplemental figure XX:Upstream regulator predictions from the differentially expressed genes for the top non-cytotoxic concentration of each PFAS. The z-score filters were relaxed to 2.0 with a Benjamini Hochberg adj p-value of 0.05 for this analysis. Blue represents inhibited upstream regulators, and orange represents activated regulators.

PFOA 20µM DAY 1

PFOA 20µM DAY 4

PFOA 20µM DAY 10

PFOA 20µM DAY 14

PFBS 100µM DAY 1

PFBS 100µM DAY 4

PFBS 100µM DAY 10

PFBS 100µM DAY 14

PFOS 20µM DAY 1

PFOS 20µM DAY 4

PFOS 20µM DAY 10

PFOS 20µM DAY 14

PFDS 100µM DAY 1

PFDS 100µM DAY 4

PFDS 100µM DAY 10

PFDS 100µM DAY 14

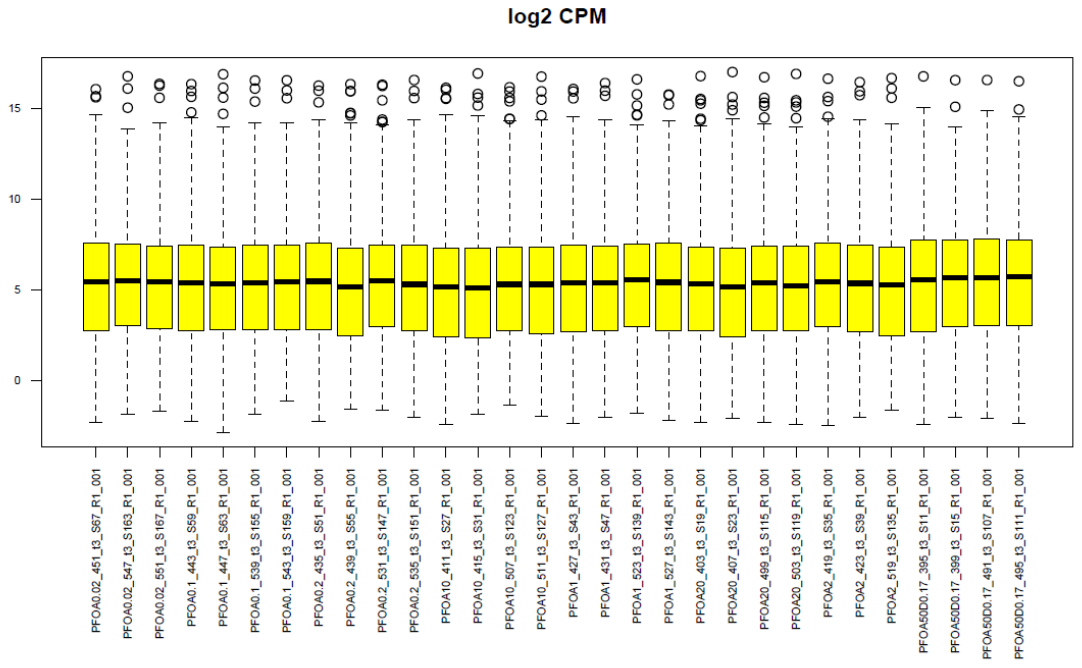

Supplemental File 4-4: PFOA boxplots of mapped reads day-10. Boxplots were generated in the R statistical environment for each sample using the filtered counts per million. Sample distributions of probes that had at least a median of 5 counts in at least one treatment group are displayed in each boxplot.

Supplemental figure XX:Upstream regulator predictions from the differentially expressed genes for the top non-cytotoxic concentration of each PFAS. The z-score filters were relaxed to 2.0 with a Benjamini Hochberg adj p-value of 0.05 for this analysis. Blue represents inhibited upstream regulators, and orange represents activated regulators.

PFOA 20µM DAY 1

PFOA 20µM DAY 4

PFOA 20µM DAY 10

PFOA 20µM DAY 14

PFBS 100µM DAY 1

PFBS 100µM DAY 4

PFBS 100µM DAY 10

PFBS 100µM DAY 14

PFOS 20µM DAY 1

PFOS 20µM DAY 4

PFOS 20µM DAY 10

PFOS 20µM DAY 14

PFDS 100µM DAY 1

PFDS 100µM DAY 4

PFDS 100µM DAY 10

PFDS 100µM DAY 14

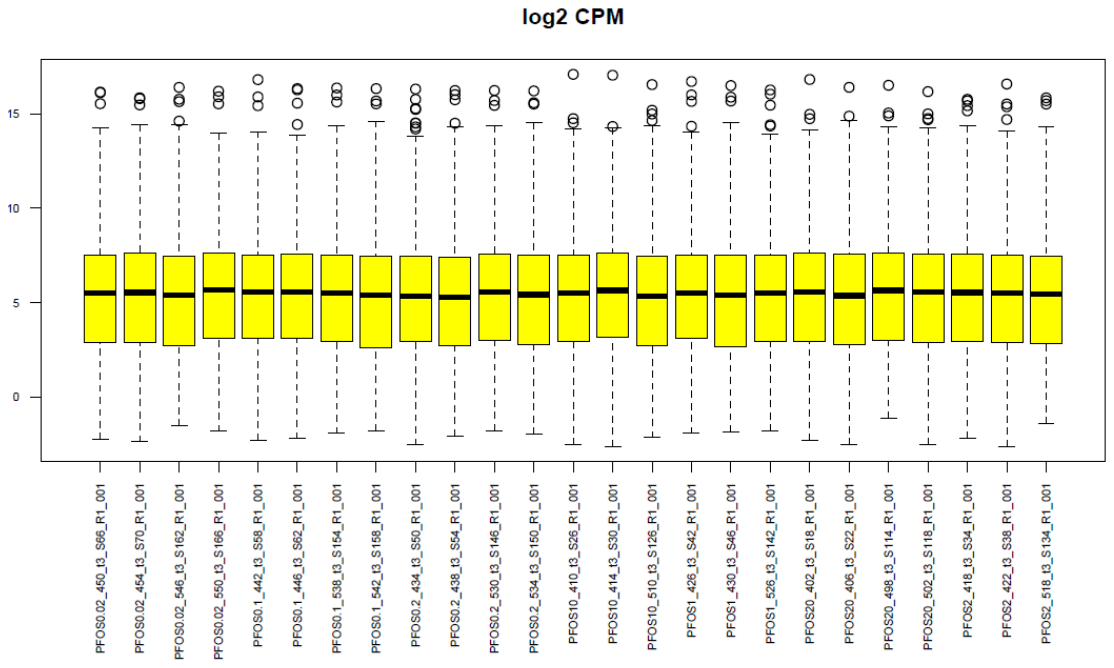
Supplemental File 4-5: PFOS boxplots of mapped reads day-10. Boxplots were generated in the R statistical environment for each sample using the filtered counts per million. Sample distributions of probes that had at least a median of 5 counts in at least one treatment group are displayed in each boxplot.

Supplemental figure XX:Upstream regulator predictions from the differentially expressed genes for the top non-cytotoxic concentration of each PFAS. The z-score filters were relaxed to 2.0 with a Benjamini Hochberg adj p-value of 0.05 for this analysis. Blue represents inhibited upstream regulators, and orange represents activated regulators.

PFOA 20µM DAY 1

PFOA 20µM DAY 4

PFOA 20µM DAY 10

PFOA 20µM DAY 14

PFBS 100µM DAY 1

PFBS 100µM DAY 4

PFBS 100µM DAY 10

PFBS 100µM DAY 14

PFOS 20µM DAY 1

PFOS 20µM DAY 4

PFOS 20µM DAY 10

PFOS 20µM DAY 14

PFDS 100µM DAY 1

PFDS 100µM DAY 4

PFDS 100µM DAY 10

PFDS 100µM DAY 14

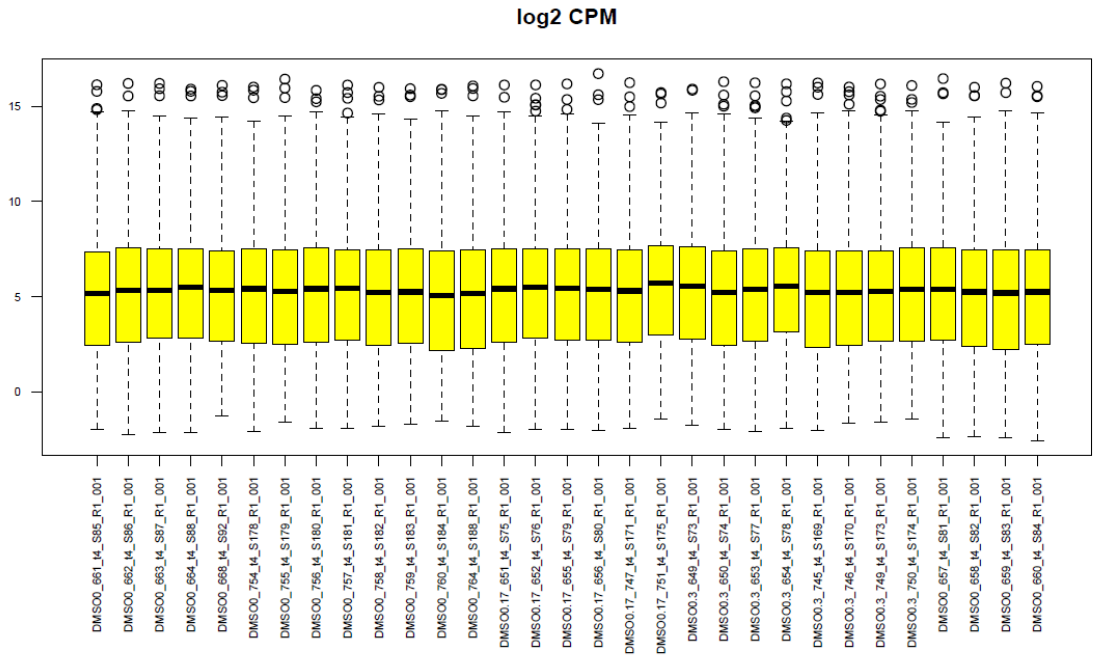

Supplemental File 5-1: DMSO boxplots of mapped reads day-14. Boxplots were generated in the R statistical environment for each sample using the filtered counts per million. Sample distributions of probes that had at least a median of 5 counts in at least one treatment group are displayed in each boxplot.

Supplemental figure XX:Upstream regulator predictions from the differentially expressed genes for the top non-cytotoxic concentration of each PFAS. The z-score filters were relaxed to 2.0 with a Benjamini Hochberg adj p-value of 0.05 for this analysis. Blue represents inhibited upstream regulators, and orange represents activated regulators.

PFOA 20µM DAY 1

PFOA 20µM DAY 4

PFOA 20µM DAY 10

PFOA 20µM DAY 14

PFBS 100µM DAY 1

PFBS 100µM DAY 4

PFBS 100µM DAY 10

PFBS 100µM DAY 14

PFOS 20µM DAY 1

PFOS 20µM DAY 4

PFOS 20µM DAY 10

PFOS 20µM DAY 14

PFDS 100µM DAY 1

PFDS 100µM DAY 4

PFDS 100µM DAY 10

PFDS 100µM DAY 14

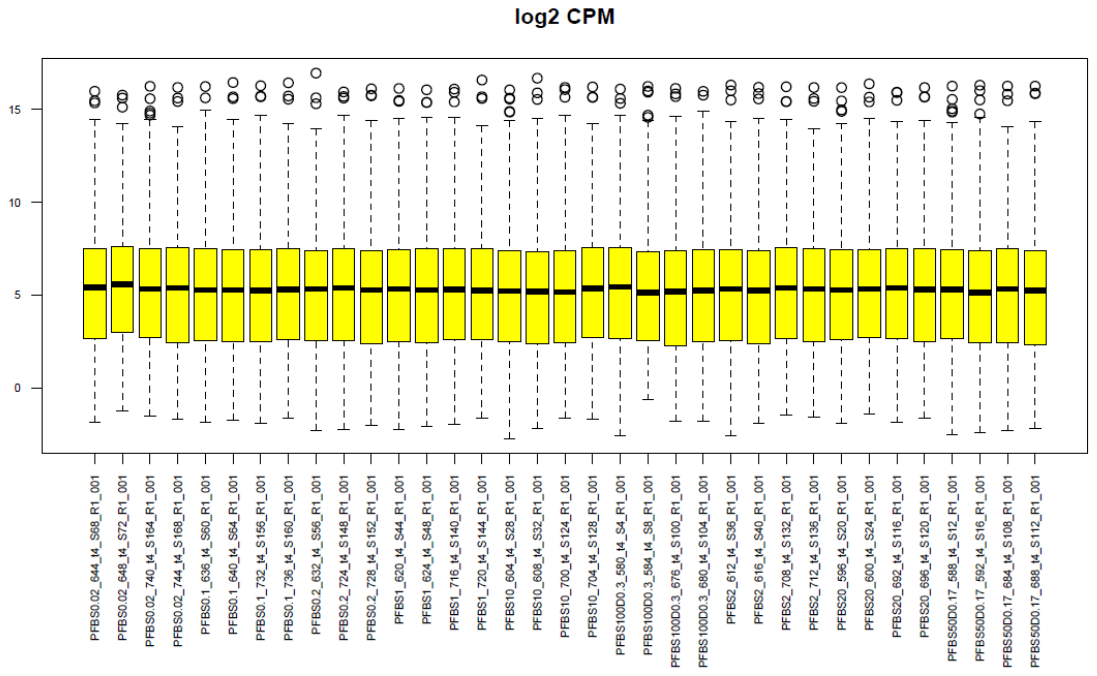
Supplemental File 5-2: PFBS boxplots of mapped reads day-14. Boxplots were generated in the R statistical environment for each sample using the filtered counts per million. Sample distributions of probes that had at least a median of 5 counts in at least one treatment group are displayed in each boxplot.

Supplemental figure XX:Upstream regulator predictions from the differentially expressed genes for the top non-cytotoxic concentration of each PFAS. The z-score filters were relaxed to 2.0 with a Benjamini Hochberg adj p-value of 0.05 for this analysis. Blue represents inhibited upstream regulators, and orange represents activated regulators.

PFOA 20µM DAY 1

PFOA 20µM DAY 4

PFOA 20µM DAY 10

PFOA 20µM DAY 14

PFBS 100µM DAY 1

PFBS 100µM DAY 4

PFBS 100µM DAY 10

PFBS 100µM DAY 14

PFOS 20µM DAY 1

PFOS 20µM DAY 4

PFOS 20µM DAY 10

PFOS 20µM DAY 14

PFDS 100µM DAY 1

PFDS 100µM DAY 4

PFDS 100µM DAY 10

PFDS 100µM DAY 14

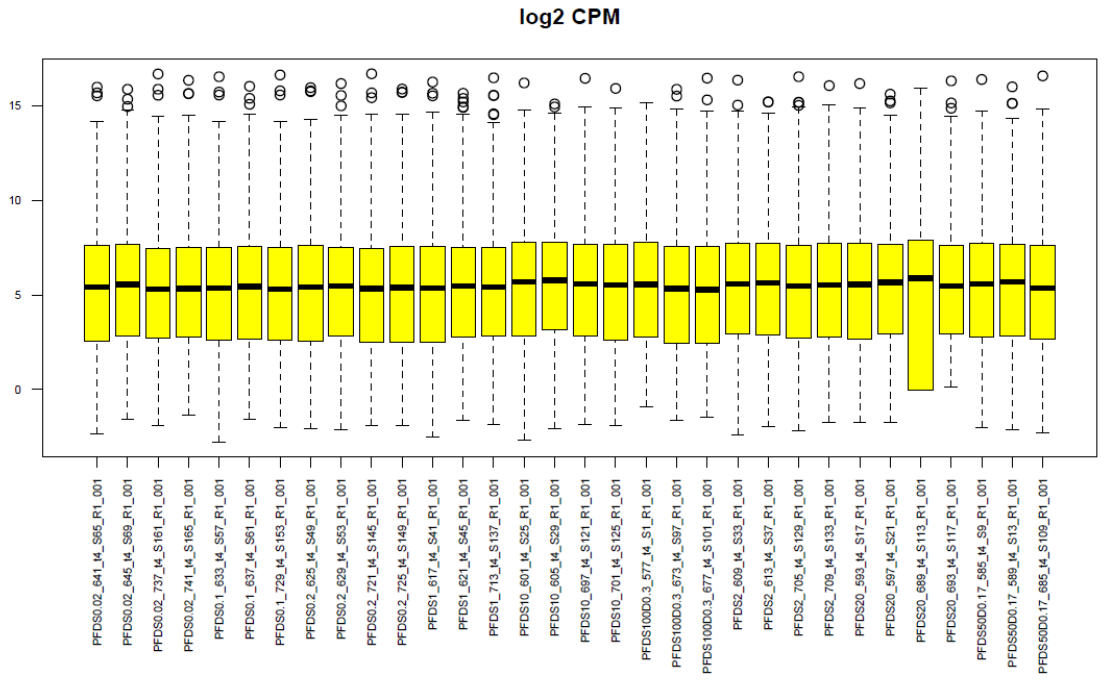
Supplemental File 5-3: PFDS boxplots of mapped reads day-14. Boxplots were generated in the R statistical environment for each sample using the filtered counts per million. Sample distributions of probes that had at least a median of 5 counts in at least one treatment group are displayed in each boxplot.

Supplemental figure XX:Upstream regulator predictions from the differentially expressed genes for the top non-cytotoxic concentration of each PFAS. The z-score filters were relaxed to 2.0 with a Benjamini Hochberg adj p-value of 0.05 for this analysis. Blue represents inhibited upstream regulators, and orange represents activated regulators.

PFOA 20µM DAY 1

PFOA 20µM DAY 4

PFOA 20µM DAY 10

PFOA 20µM DAY 14

PFBS 100µM DAY 1

PFBS 100µM DAY 4

PFBS 100µM DAY 10

PFBS 100µM DAY 14

PFOS 20µM DAY 1

PFOS 20µM DAY 4

PFOS 20µM DAY 10

PFOS 20µM DAY 14

PFDS 100µM DAY 1

PFDS 100µM DAY 4

PFDS 100µM DAY 10

PFDS 100µM DAY 14

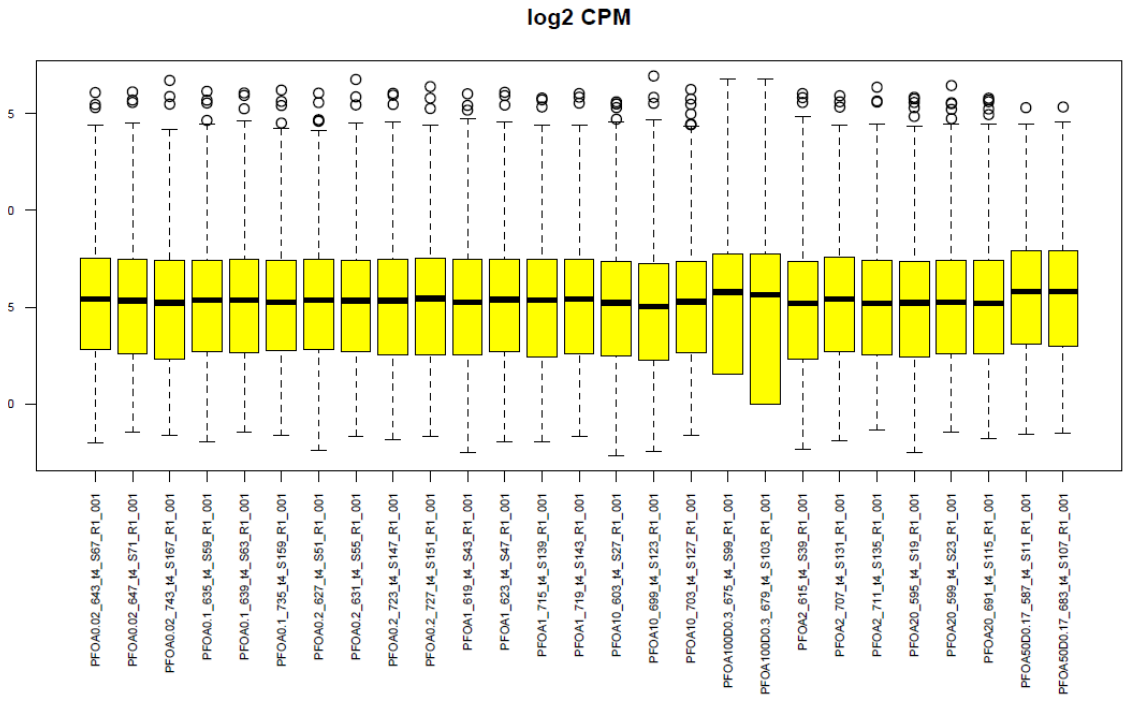
Supplemental File 5-4: PFOA boxplots of mapped reads day-14. Boxplots were generated in the R statistical environment for each sample using the filtered counts per million. Sample distributions of probes that had at least a median of 5 counts in at least one treatment group are displayed in each boxplot.

Supplemental figure XX:Upstream regulator predictions from the differentially expressed genes for the top non-cytotoxic concentration of each PFAS. The z-score filters were relaxed to 2.0 with a Benjamini Hochberg adj p-value of 0.05 for this analysis. Blue represents inhibited upstream regulators, and orange represents activated regulators.

PFOA 20µM DAY 1

PFOA 20µM DAY 4

PFOA 20µM DAY 10

PFOA 20µM DAY 14

PFBS 100µM DAY 1

PFBS 100µM DAY 4

PFBS 100µM DAY 10

PFBS 100µM DAY 14

PFOS 20µM DAY 1

PFOS 20µM DAY 4

PFOS 20µM DAY 10

PFOS 20µM DAY 14

PFDS 100µM DAY 1

PFDS 100µM DAY 4

PFDS 100µM DAY 10

PFDS 100µM DAY 14

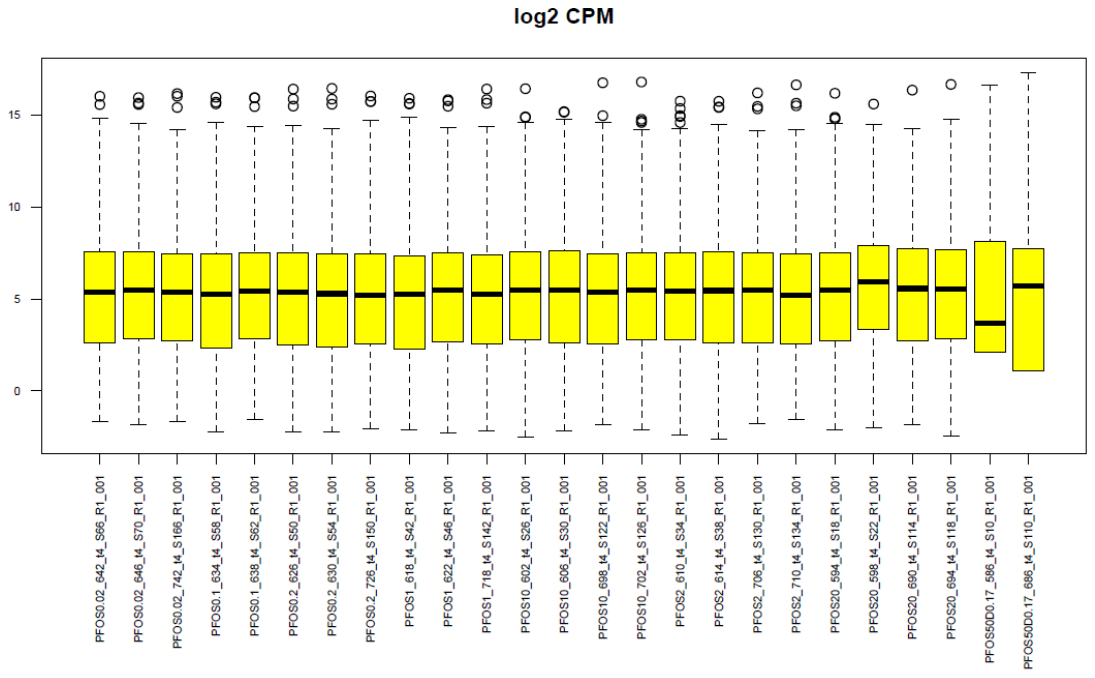
Supplemental File 5-5: PFOS boxplots of mapped reads day-14. Boxplots were generated in the R statistical environment for each sample using the filtered counts per million. Sample distributions of probes that had at least a median of 5 counts in at least one treatment group are displayed in each boxplot.

Supplemental figure XX:Upstream regulator predictions from the differentially expressed genes for the top non-cytotoxic concentration of each PFAS. The z-score filters were relaxed to 2.0 with a Benjamini Hochberg adj p-value of 0.05 for this analysis. Blue represents inhibited upstream regulators, and orange represents activated regulators.

PFOA 20µM DAY 1

PFOA 20µM DAY 4

PFOA 20µM DAY 10

PFOA 20µM DAY 14

PFBS 100µM DAY 1

PFBS 100µM DAY 4

PFBS 100µM DAY 10

PFBS 100µM DAY 14

PFOS 20µM DAY 1

PFOS 20µM DAY 4

PFOS 20µM DAY 10

PFOS 20µM DAY 14

PFDS 100µM DAY 1

PFDS 100µM DAY 4

PFDS 100µM DAY 10

PFDS 100µM DAY 14

|  | **(µM)** | **0.02** | **0.1** | **1** | **2** | **10** | **20** | **50** | **100** |
| --- | --- | --- | --- | --- | --- | --- | --- | --- | --- |
| **PFBS** | 1-Day | 3 | 4 | 4 | 4 | 4 | 3 | 4 | 4 |
| **PFDS** | 1-Day | 4 | 4 | 4 | 4 | 4 | 3 | 4 | 4 |
| **PFOA** | 1-Day | 4 | 3 | 4 | 4 | 4 | 3 | 3 | 4 |
| **PFOS** | 1-Day | 4 | 3 | 4 | 4 | 4 | 4 | cytotoxic | cytotoxic |
| **PFBS** | 4-Day | 4 | 3 | 4 | 3 | 2 | 4 | 4 | 4 |
| **PFDS** | 4-Day | 3 | 3 | 4 | 4 | 3 | 4 | 4 | 4 |
| **PFOA** | 4-Day | 4 | 3 | 4 | 4 | 2 | 3 | 4 | 3 |
| **PFOS** | 4-Day | 4 | 3 | 4 | 4 | 3 | 4 | cytotoxic | cytotoxic |
| **PFBS** | 10-Day | 4 | 3 | 3 | 3 | 4 | 3 | 3 | 4 |
| **PFDS** | 10-Day | 2 | 4 | 4 | 4 | 4 | 3 | 4 | 2 |
| **PFOA** | 10-Day | 3 | 4 | 4 | 3 | 4 | 4 | 4 | cytotoxic |
| **PFOS** | 10-Day | 4 | 4 | 3 | 3 | 3 | 4 | cytotoxic | cytotoxic |
| **PFBS** | 14-Day | 3 | 4 | 4 | 4 | 4 | 4 | 4 | 4 |
| **PFDS** | 14-Day | 4 | 3 | 3 | 4 | 4 | 3 | 3 | 3 |
| **PFOA** | 14-Day | 3 | 3 | 4 | 3 | 3 | 3 | cytotoxic | cytotoxic |
| **PFOS** | 14-Day | 4 | 3 | 4 | 3 | 3 | 3 | cytotoxic | cytotoxic |

Supplemental File 6A: The number of replicates (n) included for each sample concentration.

| DMSO | total number of replicates pre-filtering (n= ) | total number of replicates post-filtering (n= ) |
| --- | --- | --- |
| 0.30% | 32 | 29 |
| 0.17% | 32 | 30 |
| 0.10% | 80 | 69 |

Supplemental File 6B: The number of control replicates (n) pre and post filtering

For DMSO 0.3%

newProbeLog2CPMvalue = log2(probeCPMvalue) - [log2(mean(DMSO0.3)) – log2(mean (DMSO0.1)]

For DMSO 0.17%

newProbeLog2CPMvalue = log2(probeCPMvalue) - [log2(mean(DMSO0.17)) – log2(mean (DMSO0.1)]

If a plate correction was made:

Data were 2^x^, then

newProbeLog2CPMvalue = log2(probeCPMvalue) - [log2(mean(DMSO0.1ForPlateX)) – log2(mean(DMSO0.1ForAllPlates))]

in order to correct for the plate effect.

Supplemental File 7. Pipeline code

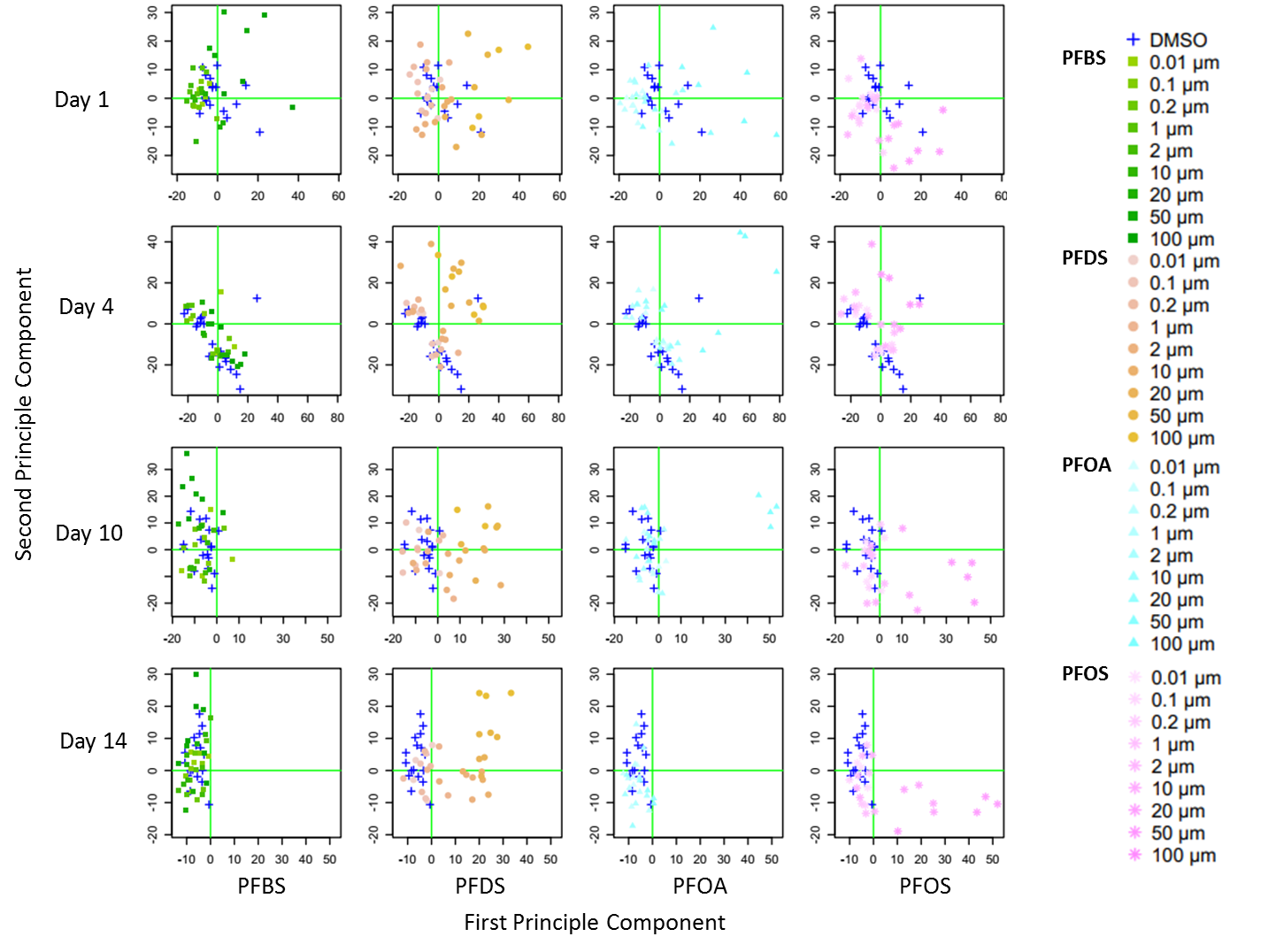

B)
C)

PFBS

PFOA

PFDS

PFOS

Day 1 1

Day 4

Day 10

Day 14

First Principle Component

Second Principle Component

**PFBS**

**PFDS**

**PFOA**

**PFOS**

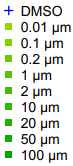

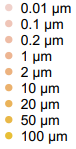

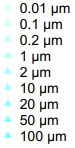

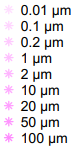

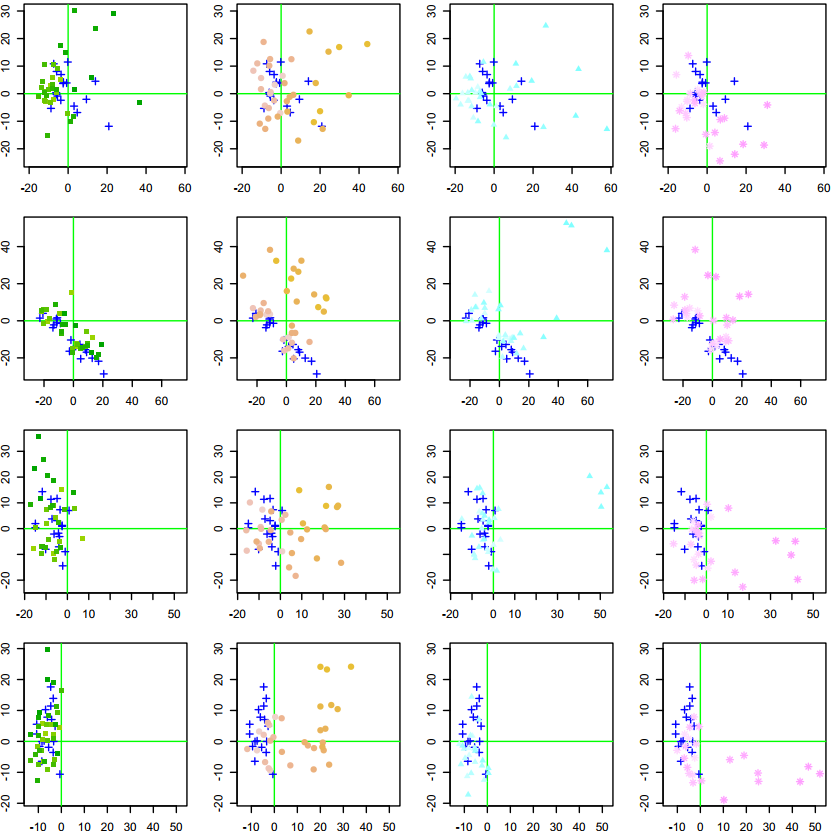

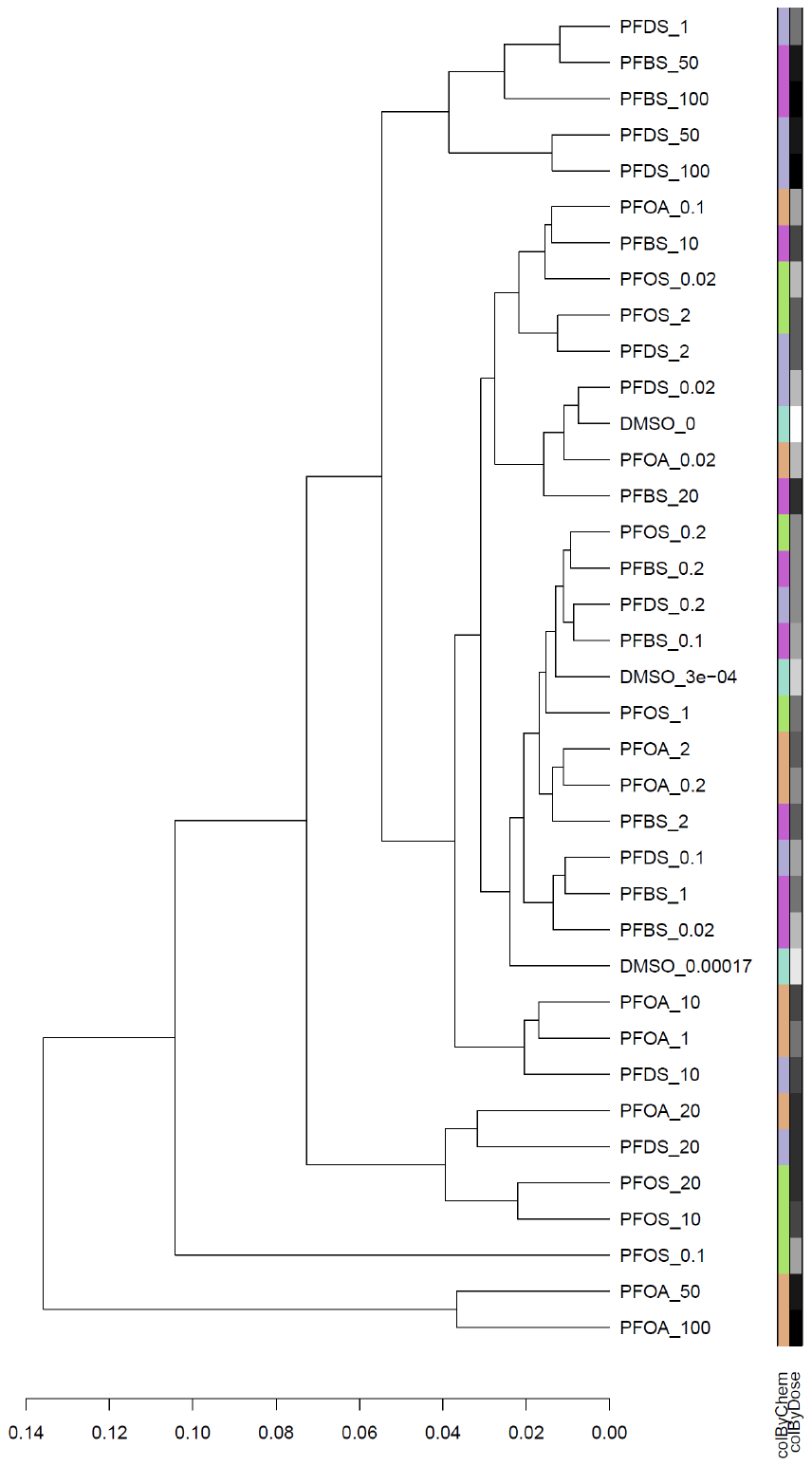

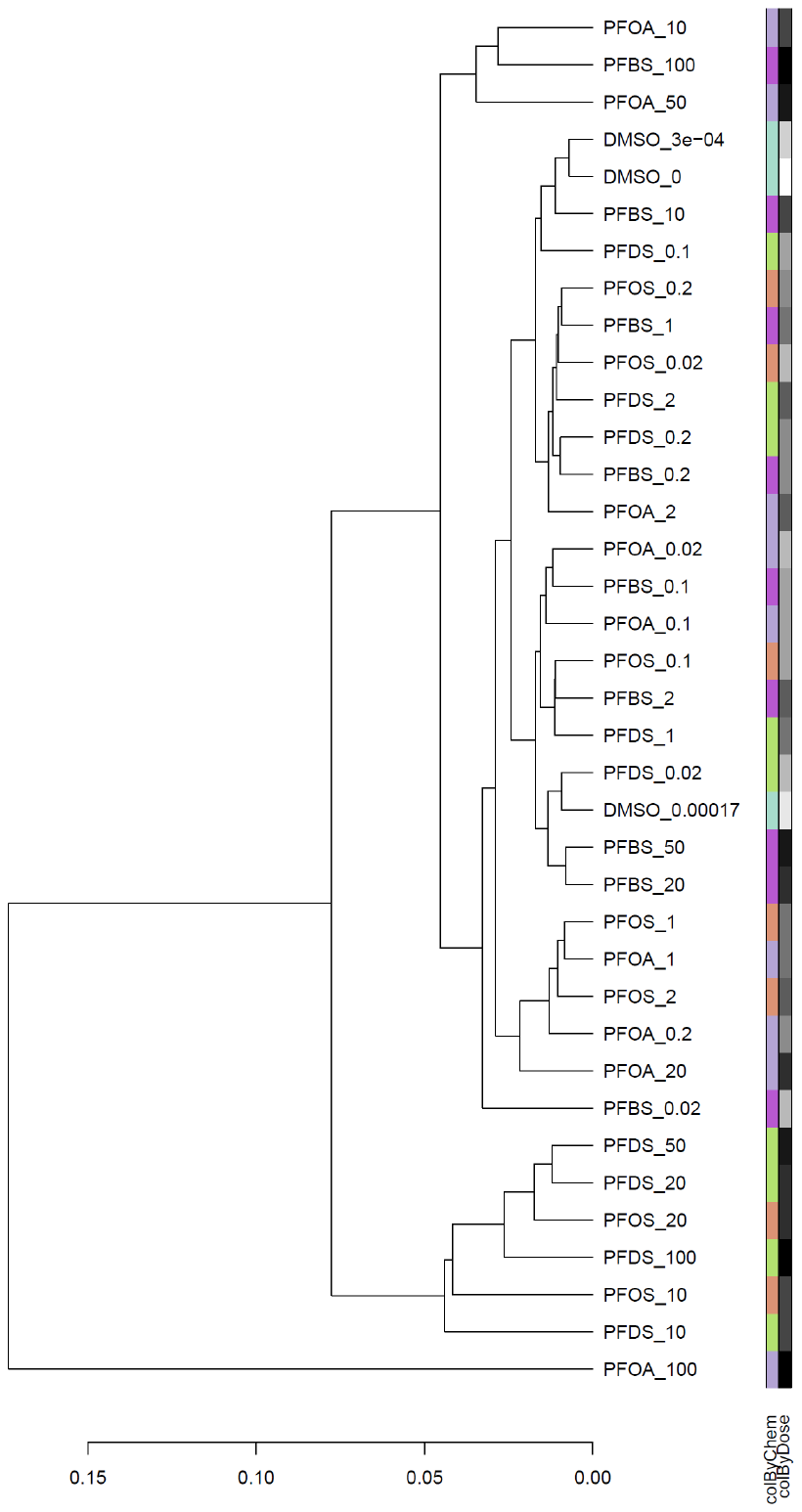

Day 1

Day 4

chemical

concentration

chemical

concentration

D)

Day 14

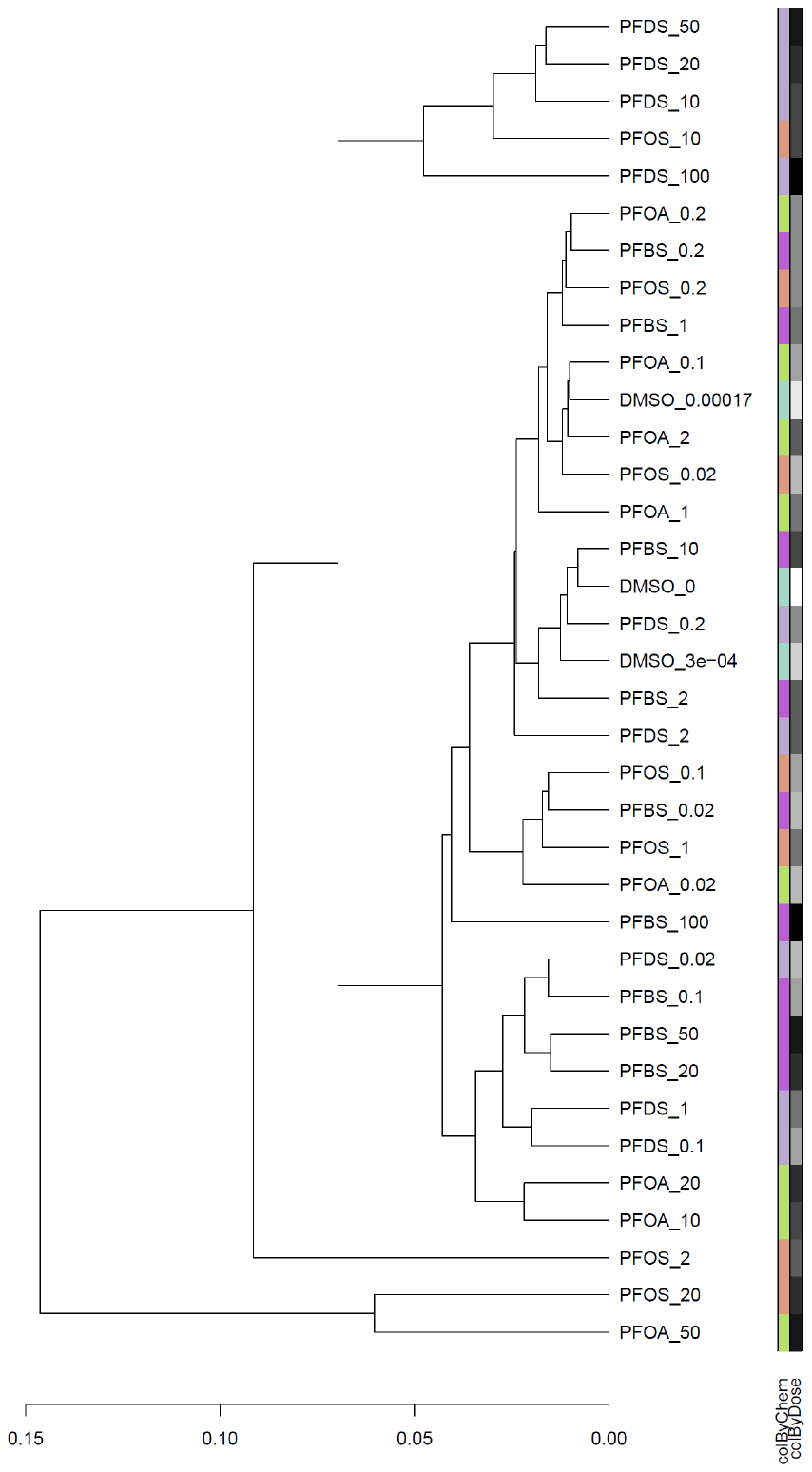

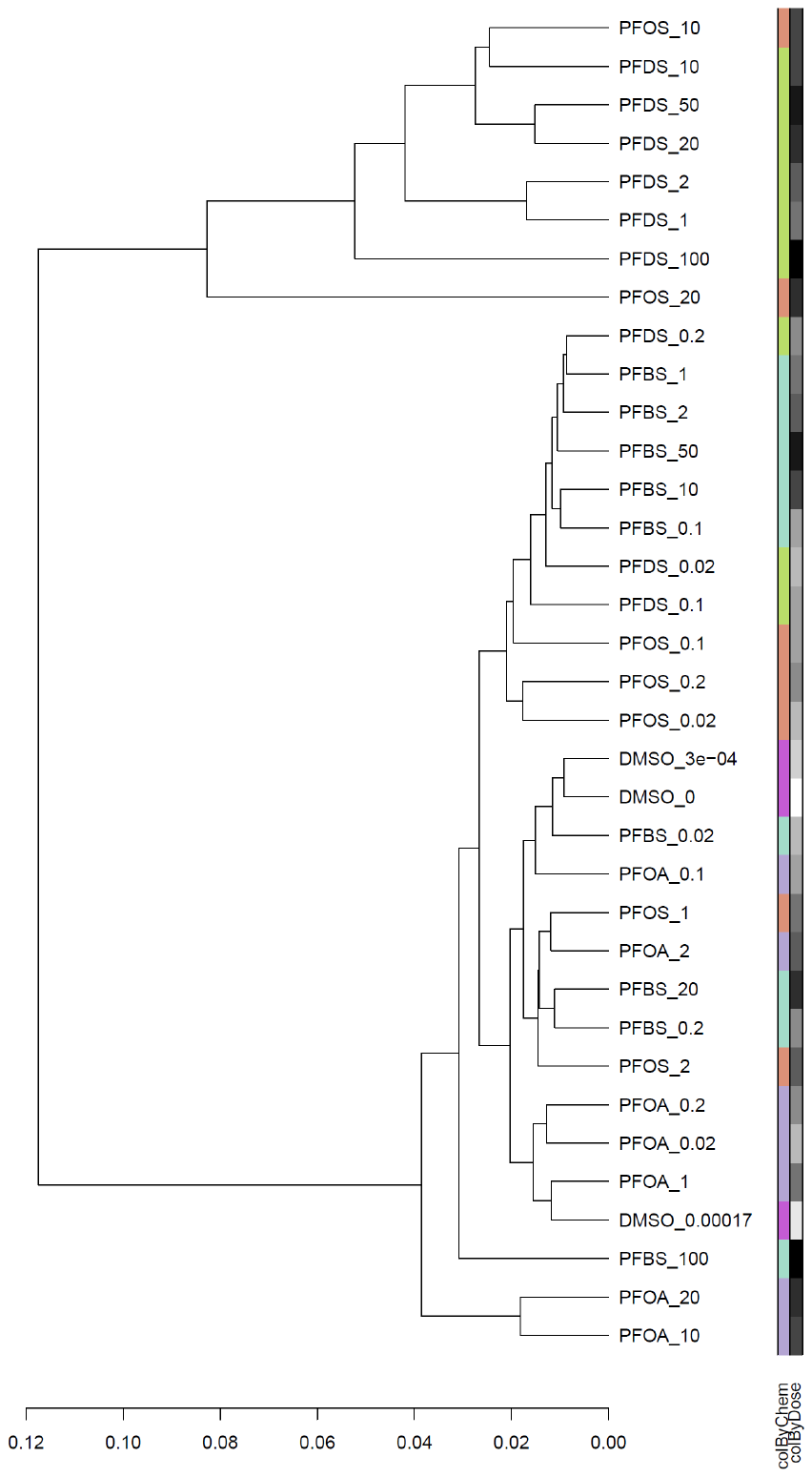

Day 10

chemical

concentration

chemical

concentration

D)

Supplemental File 8: (A) Principal Component Analysis conducted on the probes passing filters (FDR adjusted p-value < 0.05 and an absolute fold change > 1.5 on a linear scale) at each of the four time points of the four individual PFAS. Each point represents the individual replicates. The horizontal and vertical green lines at zero were added to identify the four quadrants in each of the four plots. Sample size was n = 3-4 for all exposure groups, except for n = 2 on day-4 PFBS 10 µM and PFOA 10 µM, and n = 2 on day-10 PFDS 0.02 and 100 µM. Note that an outlier within the solvent controls is apparent in the upper right quadrant of Day 4. This was thus removed from subsequent analysis, the DEGs recalculated, and the PCA replotted in (B). Hierarchical clustering was conducted on differentially expressed genes (post filtering and removal of outliers) from the four PFAS for all concentrations at (C) days 1 and 4, and (D) days 10 and 14, using complete linkage and one minus spearman correlation (n=2 to 4).

<https://github.com/mattjmeier/2020_PFAS_DEGs/tree/main/BMDExpress2>

Supplemental File 9:Link to github bm2 files

Supplemental figure XX:Upstream regulator predictions from the differentially expressed genes for the top non-cytotoxic concentration of each PFAS. The z-score filters were relaxed to 2.0 with a Benjamini Hochberg adj p-value of 0.05 for this analysis. Blue represents inhibited upstream regulators, and orange represents activated regulators.

PFOA 20µM DAY 1

PFOA 20µM DAY 4

PFOA 20µM DAY 10

PFOA 20µM DAY 14

PFBS 100µM DAY 1

PFBS 100µM DAY 4

PFBS 100µM DAY 10

PFBS 100µM DAY 14

PFOS 20µM DAY 1

PFOS 20µM DAY 4

PFOS 20µM DAY 10

PFOS 20µM DAY 14

PFDS 100µM DAY 1

PFDS 100µM DAY 4

PFDS 100µM DAY 10

PFDS 100µM DAY 14

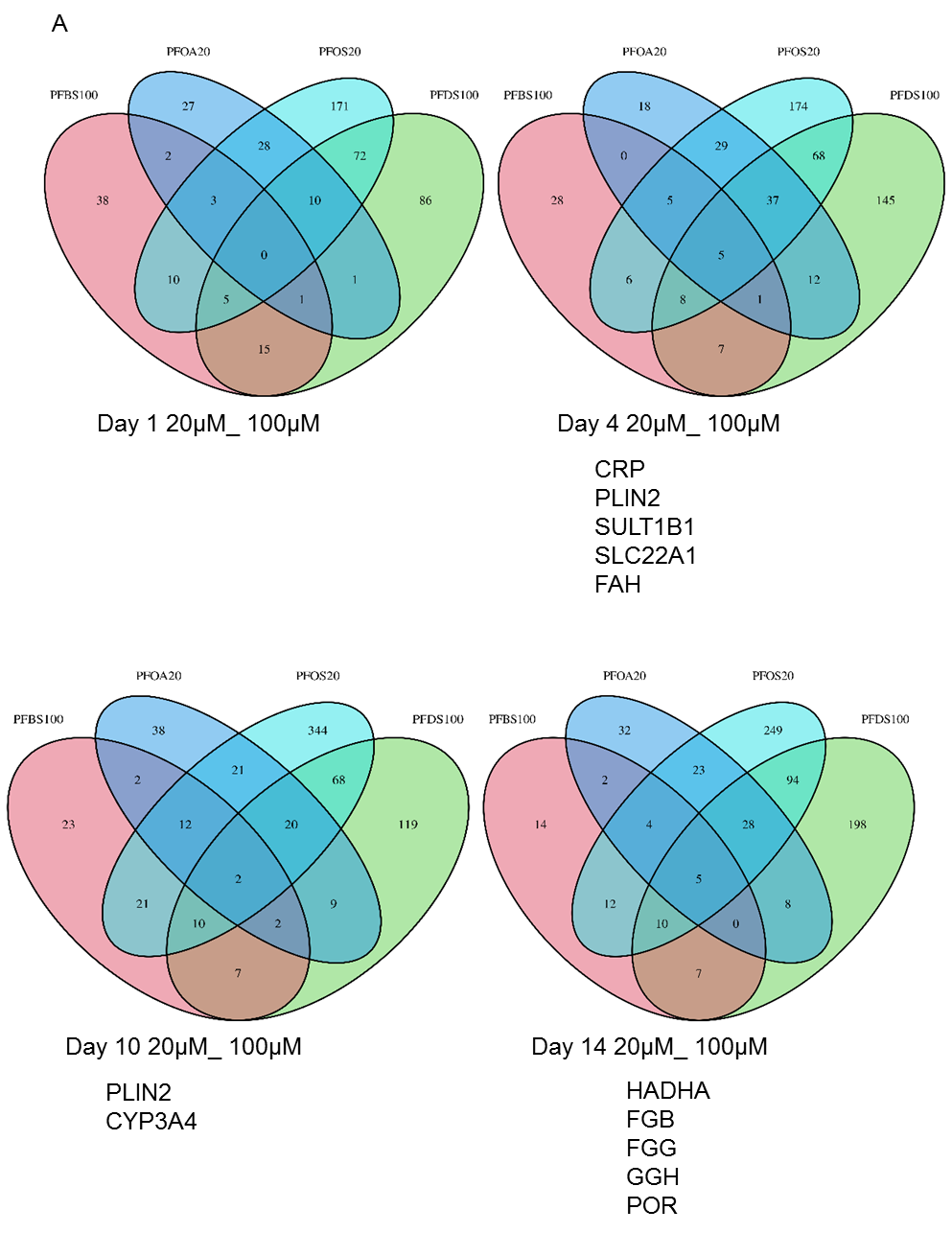

B

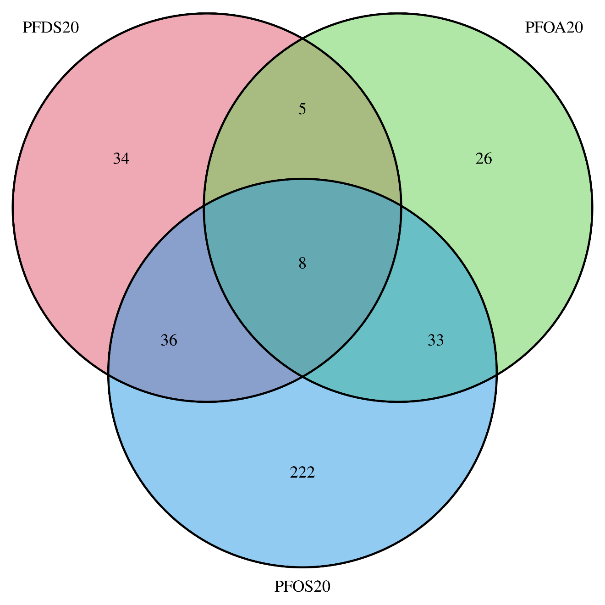

Day 1 20µM

Day 10 20µM

Day 14 20µM

TNFSF10

FGFR2

SLC1A2

GSTA2

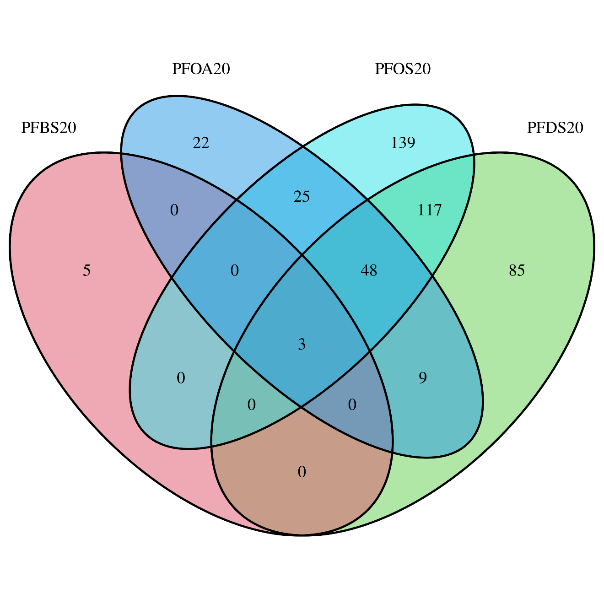

FAM96A

UGT2B7

APP

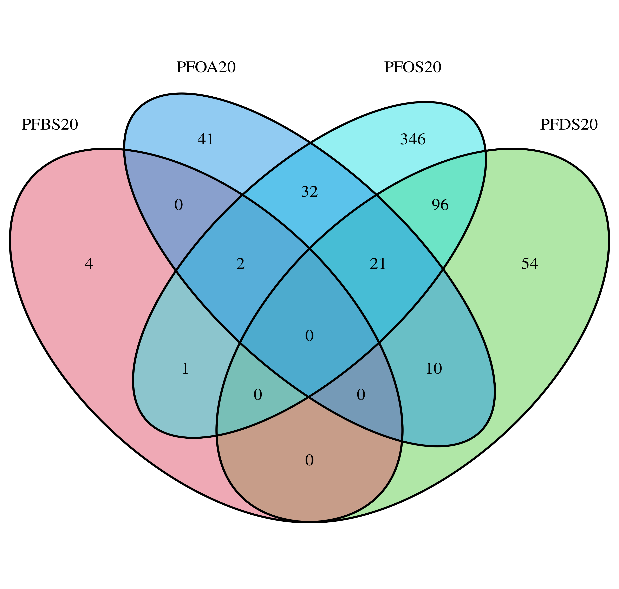

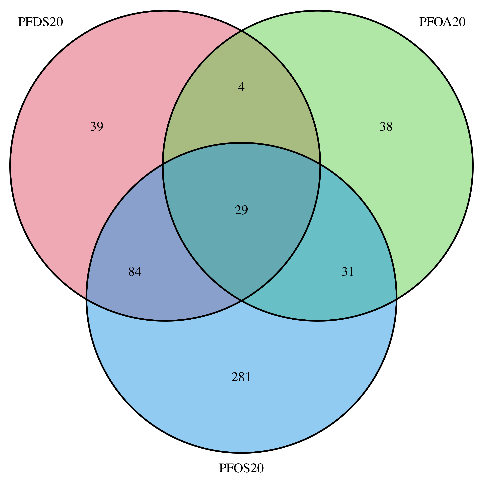

AGXT2

UGT2B7

SLC27A2

DSG2

Day 4 20µM

FN1_27231

FN1_2459

FN1_13666

GGH

UGT2B7

SLC22A1

IGFBP1

GSTA5

PLIN2

HMGCS2

MLLT11

IGFBP1

HP

COMT

GCLM

FMO5

CYP2A6

CYP4A11

SAA1

CYP7A1

TRIB3

GCLM

SLC10A1

SERPINE1

FAM96A

CIDEC

CXCL1

EEF1G

Supplemental File 10. Venn diagrams of differentially expressed genes illustrating genes that were in common over treatments and time. Panel A) Analysis of the differentially expressed genes for each PFAS at their top non-cytotoxic concentration (PFBS 100 µM, PFDS 100 µM, PFOA 20 µM, and PFOS 20 µM).

Panel B) Analysis of the differentially expressed genes for each PFAS at the 20 µM concentration. Days 1 and14 for PFBS did have any genes meeting the filtering criteria (FDR p<0.05 and FC±1.5).

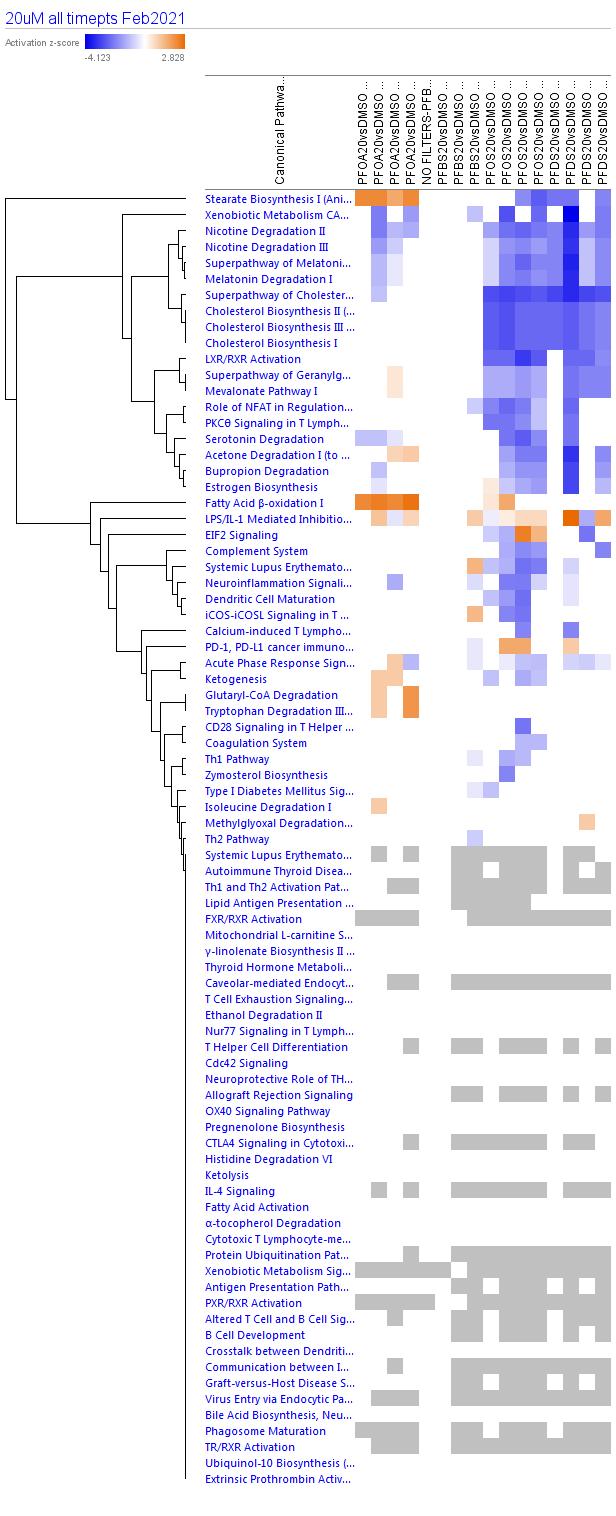

PFOA 20µM DAY 1

PFOA 20µM DAY 4

PFOA 20µM DAY 10

PFOA 20µM DAY 14

PFBS 20µM DAY 1

PFBS 20µM DAY 4

PFBS 20µM DAY10

PFBS 20µM DAY 14

PFOS 20µM DAY 1

PFOS 20µM DAY 4

PFOS 20µM DAY 10

PFOS 20µM DAY 14

PFDS 20µM DAY 1

PFDS 20µM DAY 4

PFDS 20µM DAY 10

PFDS 20µM DAY 14

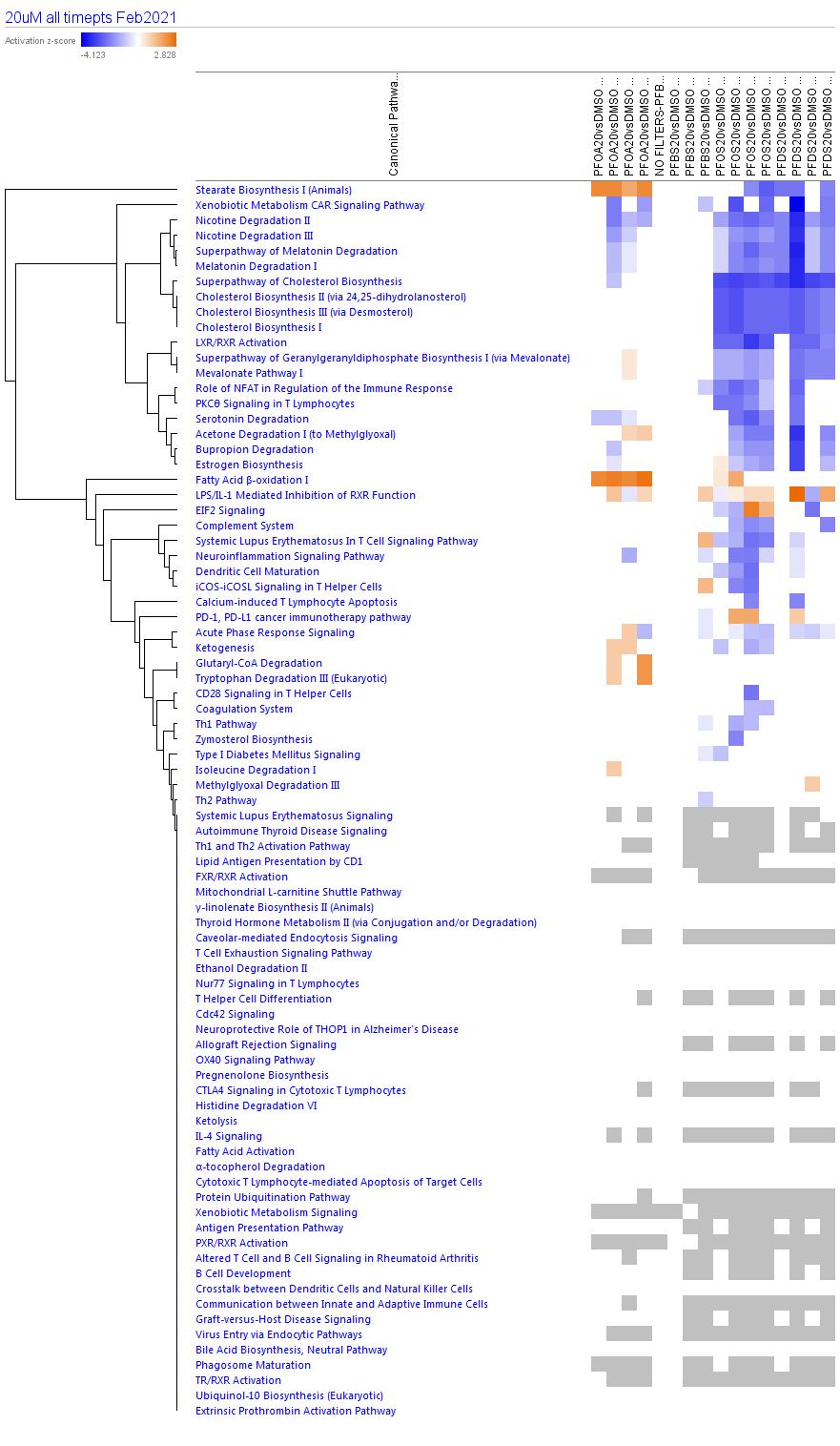

A

PFOA 20µM DAY 1

PFOA 20µM DAY 4

PFOA 20µM DAY 10

PFOA 20µM DAY 14

PFBS 100 µM DAY 1

PFBS 100 µM DAY 4

PFBS 100 µM DAY 10

PFBS 100 µM DAY 14

PFOS 20µM DAY 1

PFOS 20µM DAY 4

PFOS 20µM DAY 10

PFOS 20µM DAY 14

PFDS 100 µM DAY 1

PFDS 100 µM DAY 4

PFDS 100 µM DAY 10

PFDS 100 µM DAY 14

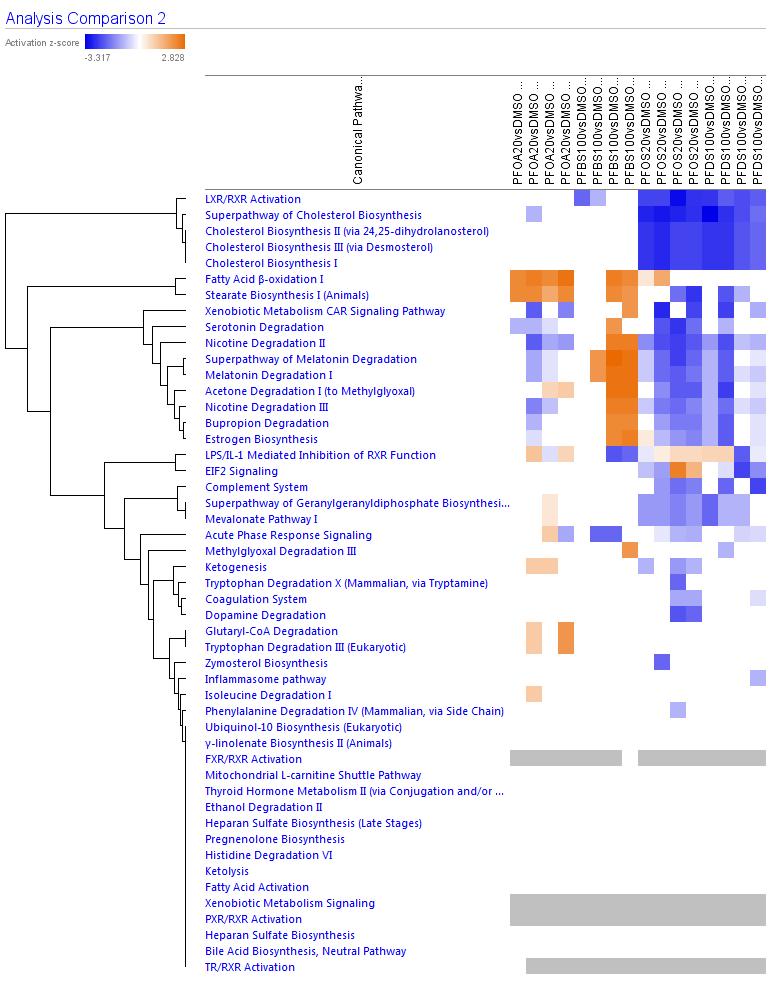

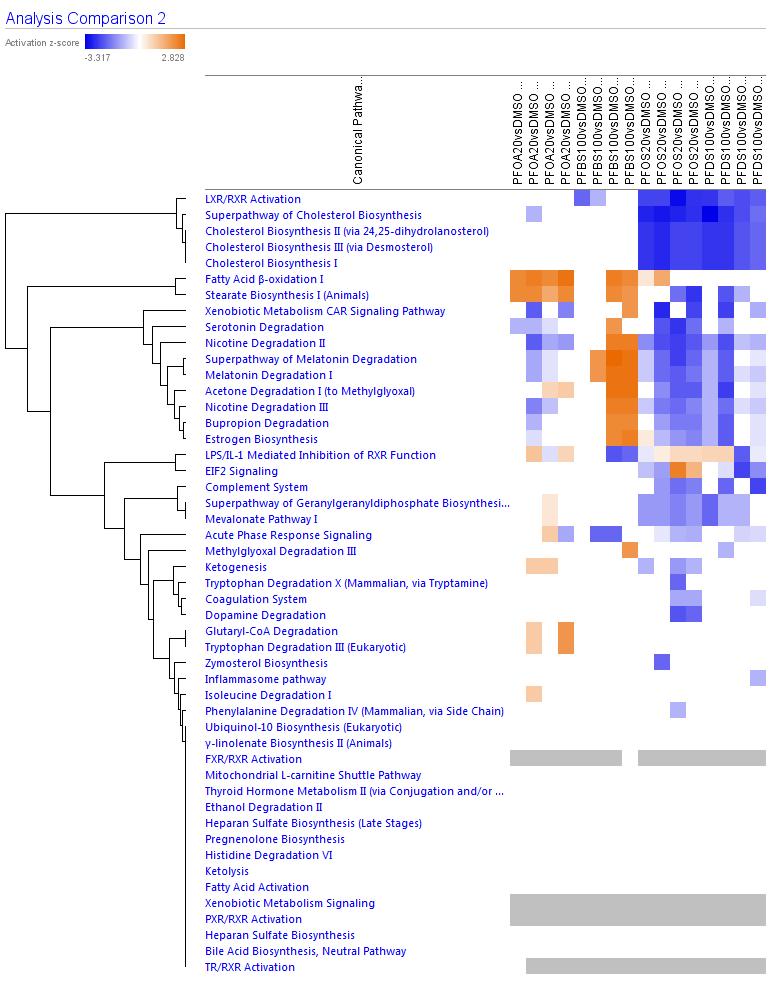

B

Supplemental File 11. Pathway analysis of the differentially expressed genes for all time points at the 20 µM concentration (A) or for the highest non-cytotoxic concentrations (B). For (B) concentrations were 20 µM for PFOA and PFOS, and 100 µM for PFBS and PFDS. Filters for this analysis were less restrictive than in Figure 4 A&B, no z-score filter was applied only a Benjamin–Hochberg adjusted p-value ≤ 0.05. Blue represents inhibited pathways, orange represents activated pathways, grey and white represent pathways that are statistically significant, but do not have a predicted direction (inhibited/activated). n = 3-4 for all exposure groups, except for n = 2 on day-10 PFDS 100 µM. Supplemental Figure 12 illustrates a more detailed pathway analysis for PFOS over all time points and concentrations.

0.02 µM

0.1	µM

0.2	µM

1	µM

2	µM

10	µM

20	µM

PFOS day1

PFOS day10

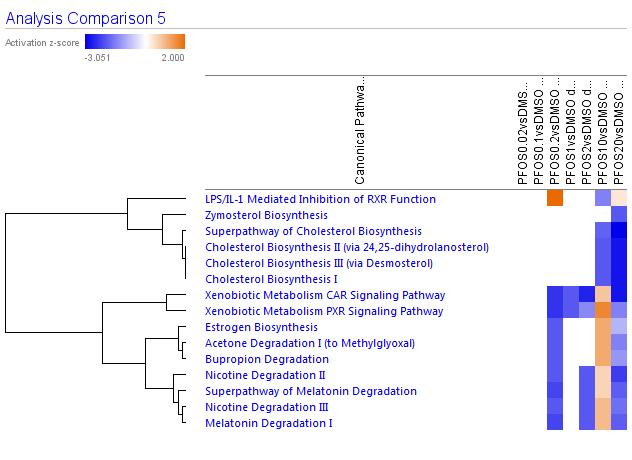

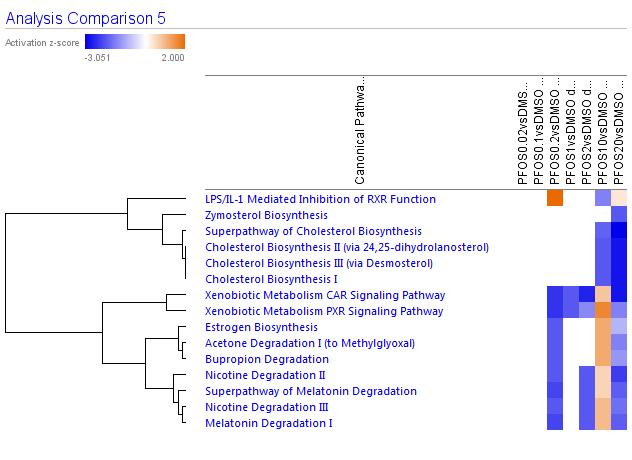

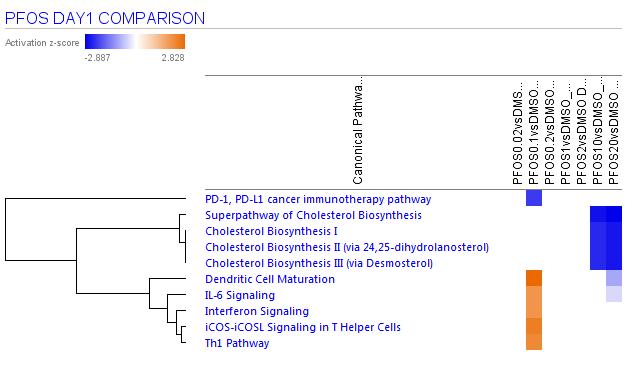

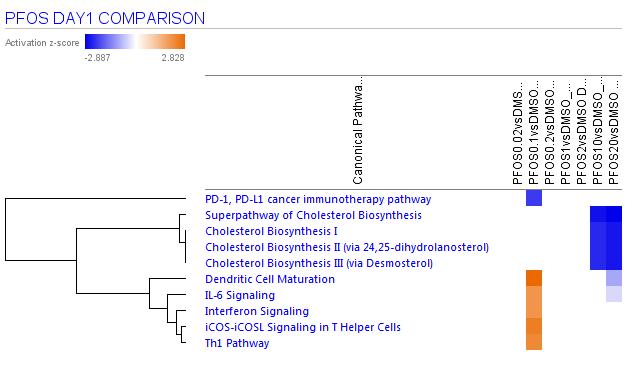

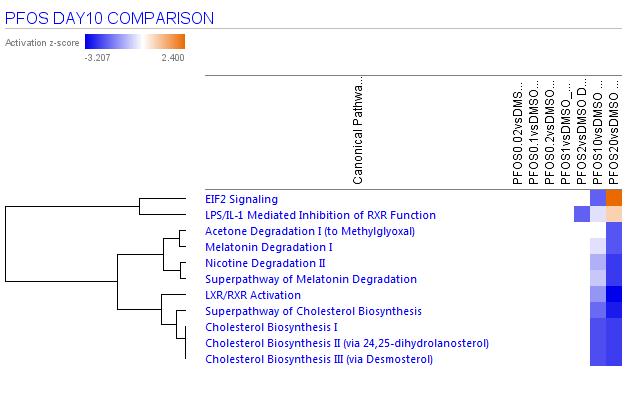

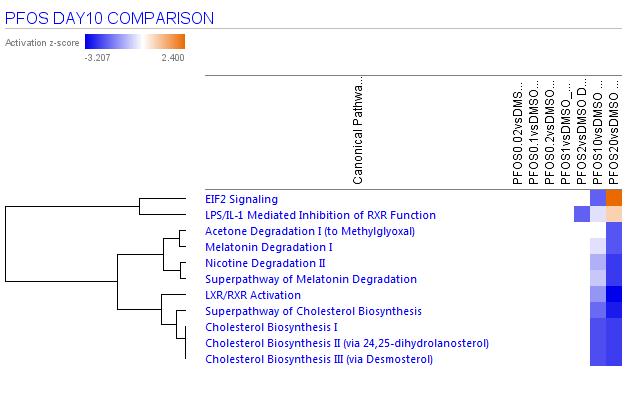

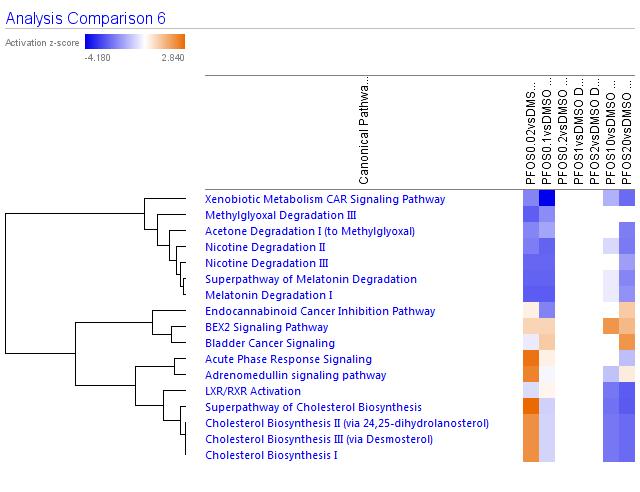

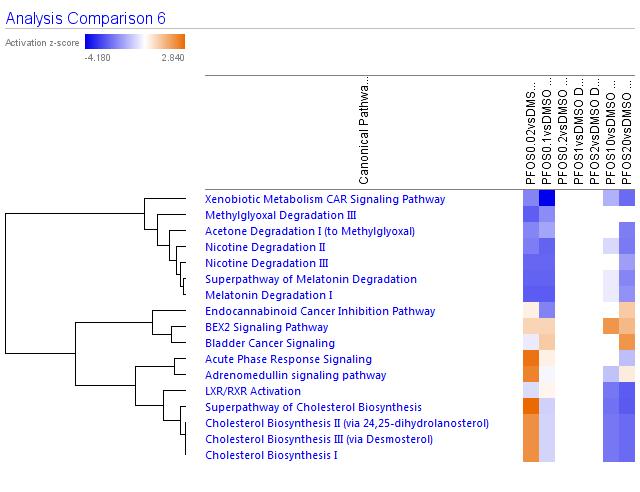

0.02 µM

0.1	µM

0.2	µM

1	µM

2	µM

10	µM

20	µM

PFOS day4

PFOS day14

0.02 µM

0.1	µM

0.2	µM

1	µM

2	µM

10	µM

20	µM

0.02 µM

0.1	µM

0.2	µM

1	µM

2	µM

10	µM

20	µM

Supplemental File 12. Pathway enrichment for all concentrations of PFOS over time points are shown as an example of how pathway enrichment increased with concentration. Filters for this analysis were set to z-score = to 2.0 and Benjamini–Hochberg adjusted p-value ≤ 0.05. Blue represents inhibited pathways and orange represents activated pathways.

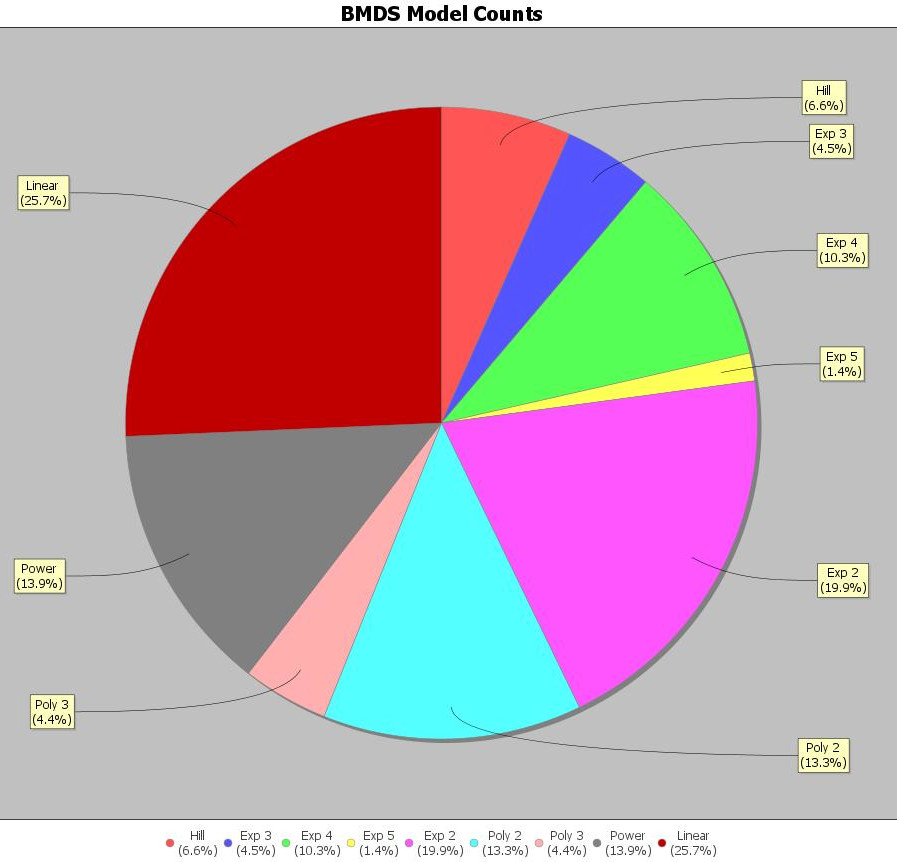

Supplemental File 13. Summary of the models used for the benchmark concentration analysis for PFAS over 1, 4, 10, and 14 days. Analysis conducted in BMDExpress v2.2.

| Day | PFAS | BMC | Lower Confidence Interval | Upper Confidence Interval | # of genes that were modelled |
| --- | --- | --- | --- | --- | --- |
| 1 | PFBS | 53.5 | 49.4 | 59.9 | 191 |
| 1 | PFOA | 39.9 | 35.5 | 43.6 | 403 |
| 1 | PFDS | 24.9 | 19.5 | 32.5 | 278 |
| 1 | PFOS | 8.2 | 7.4 | 8.5 | 309 |
| 4 | PFBS | 69.7 | 64.8 | 74.2 | 136 |
| 4 | PFOA | 56.1 | 54.1 | 58.6 | 905 |
| 4 | PFDS | 11.5 | 9.0 | 14.3 | 571 |
| 4 | PFOS | 9.1 | 7.7 | 9.9 | 364 |
| 10 | PFBS | 46.3 | 42.1 | 50.4 | 223 |
| 10 | PFOA | 32.1 | 29.2 | 33.9 | 630 |
| 10 | PFDS | 12.9 | 10.8 | 16.3 | 341 |
| 10 | PFOS | 8.2 | 7.0 | 9.0 | 459 |
| 14 | PFBS | 60.2 | 55.7 | 65.1 | 170 |
| 14 | PFOA | 6.4 | 2.7 | 7.9 | 131 |
| 14 | PFDS | 17.9 | 12.5 | 25.7 | 404 |
| 14 | PFOS | 8.6 | 7.3 | 9.1 | 537 |

Supplemental File 14. Overall the benchmark concentrations (95 % CI) by day, and the number of genes modelled.
